# Viscoelasticity and Interface Properties of Multi-Component Condensates Govern Protein Sequestration and Suppression of Amyloid Formation

**DOI:** 10.64898/2025.12.29.695806

**Authors:** Tharun Selvam Mahendran, Xinrui Gui, Anne Bremer, Anurag Singh, Fatima K. Zaidi, Giulio Tesei, Joseph L. Basalla, Sagar Chittori, Melissa R. Marzahn, Bhargavi Gindra, Richoo B. Davis, Tapojyoti Das, Kresten Lindorff-Larsen, Davit A. Potoyan, Tanja Mittag, Priya R. Banerjee

## Abstract

Stress granules (SGs) are multi-component biomolecular condensates widely implicated as sites of protein aggregation by virtue of the high concentrations of amyloidogenic RNA-binding proteins they contain. This model, in which SGs are viewed as crucibles for amyloid formation, has not been rigorously tested. Here, we employed twelve multi-component protein–nucleic acid condensate systems as SG-mimics with diverse physicochemical features. Utilizing three fibril-forming proteins, hnRNPA1, Tau, and FUS, which are concentrated more than 50-fold in condensates, we report that multi-component biomolecular condensates robustly suppress, rather than promote, amyloid formation. Multiscale experimental analyses, including quantitative kinetic measurements, rheology, and microscopy, combined with computational modelling, reveal that condensates serve as sinks for soluble protein, and fibrils form in the dilute phase, although interfaces can promote nucleation. Three key physicochemical properties of condensates govern suppression of fibril formation: First, condensate-mediated sequestration lowers the concentration of fibril-forming proteins in the dilute phase. Second, condensate viscoelasticity constrains efflux-driven fibril growth in the dilute phase. And third, dilution of fibril-forming proteins at condensate interfaces mitigates fibril nucleation. The sink potential of SG-mimics is recapitulated in G3BP1–RNA condensates and SGs reconstituted in mammalian cell lysate, suggesting that SGs may have evolved to suppress stress-induced protein aggregation.

## Introduction

Stress-responsive phase separation of G3BP1/G3BP2 and other RNA-binding proteins (RBPs) with untranslated mRNAs mediates the formation of stress granules (SGs) and is important for cell survival^1–6^. Yet, multiple lines of genetic, histopathological, and cell-biological evidence suggest that the prolonged assembly of SGs drives the pathogenesis of neurodegenerative diseases such as amyotrophic lateral sclerosis (ALS), frontotemporal dementia, and multisystem proteinopathy^7–10^. The shared pathological hallmark of these diseases is the presence of solid, fibrillar deposits formed by specific RBPs in patient cells. Because fibril-forming RBPs such as hnRNPA1, TIA1, FUS, and TDP-43 are enriched in SGs, these assemblies are often regarded as potential sites^11^, or even crucibles^12^, for amyloid fibril formation^13^. The link between SGs and protein fibril formation is supported by the fact that fibrils typically appear at the sites of condensates *in vitro*, an observation that has been interpreted to indicate that the high local concentration of protein in condensates promotes nucleation and feeds growth of fibrils^14–16^. Furthermore, fibrils appear earlier in samples containing condensates than in samples without condensates^16^. These insights have led to a model where SGs are thought to promote fibril formation. By contrast, several cell biological lines of evidence support an alternative view that SGs may be protective. For example, TDP-43 is relatively better protected against aggregation when located in SGs than in other types of condensates^17^. Furthermore, pathological inclusions of cytoplasmic TDP-43 can arise independently of SGs, suggesting that protein aggregation may proceed more readily in the dilute phase outside condensates^18–20^.

Employing mono-component systems, our recent characterization of the influence of phase separation of the prion-like low-complexity domain (LCD) of hnRNPA1 (A1-LCD) on its fibril formation suggested that condensates are metastable assemblies that can act as off-pathway sinks against fibril formation^21^. However, some aspects of these homotypic condensates can nevertheless promote fibril formation. For example, the condensate interfaces can accelerate fibril nucleation^21–24^, explaining earlier observations that fibrils appear earlier in the presence of condensates^16^ and are apparently attached to them^15^. A limitation of mono-component systems in which the same fibril-forming protein also drives condensate formation is that they do not recapitulate the multi-component nature of SGs.

The contrasting views of condensates as crucibles or sinks and the potentially opposing roles of condensate interiors versus interfaces raise several fundamental questions that must be addressed to understand the role SGs play in neurodegenerative diseases. (1) Do multi-component condensates that resemble SGs (i.e., SG mimics) enhance or suppress fibril formation? (2) Does the presence of heterotypic interactions in multi-component condensates inhibit fibril-promoting interactions more effectively compared to the purely homotypic interactions in mono-component condensates? (3) How do condensate interiors and interfaces contribute to the effects of multi-component SG-like condensates on fibril formation? (4) Condensates are viscoelastic fluids with sequence- and composition-encoded physicochemical properties^25^. Are there discernible physical rules that govern how distinct condensate properties, such as viscoelasticity, protein partitioning, and transport across the condensate interface, contribute to their modulation of fibril formation?

Herein, we leverage multi-component peptide–nucleic acid condensates whose components are facsimiles of relevant SG building blocks, and which have programmable physical properties, to characterize systematically, quantitatively, and mechanistically how multi-component condensates influence amyloid formation of proteins that are implicated in neurodegenerative diseases. SGs host an intricate network of homotypic and heterotypic interactions between nucleic acids and proteins containing LCDs^26^. Among distinct LCDs, prion-like domains (PLDs) and arginine/glycine-rich repeat (RGG) domains are of interest due to their large abundance in the SG proteome^26–29^. Some PLD-containing proteins carry disease-related mutations and form fibrillar inclusions in patient tissues^30^. The D262V mutation in hnRNPA1 causes highly penetrant disease in affected families^30^. Therefore, we selected the wild-type (WT) A1-LCD and its pathogenic variant, D262V, as one of the fibril-forming proteins of interest. To mimic the SG microenvironment, we employed RGRGG-repeat peptides, which, together with single-stranded DNA or RNA, form condensates that favorably recruit A1-LCD. By rationally altering the peptide sequence or the nucleic acid chain length, we tune the viscoelastic properties of the resulting condensates by nearly three orders of magnitude, enabling the mechanistic dissection of their effects on A1-LCD fibril formation. We demonstrate that condensate-mediated impact on fibril assembly can be extended to orthogonal fibril-forming protein systems as well as full-length counterparts. Furthermore, we show that our findings directly extend to condensates formed by poly-A RNA and the major SG hub protein G3BP1^2,6^, and to stress granules reconstituted from human cell lysate. Together, our results demonstrate that SGs may not be crucibles for neurodegenerative disease but instead protect proteins that preferentially partition into them from fibril formation and mitigate fibril nucleation at interfaces by limiting the density of fibril-forming proteins.

### Multi-component biomolecular condensates suppress amyloid formation

To determine how physical properties of SG-like multi-component condensates influence amyloid fibril formation, we first employed condensates formed by single-stranded (ss) poly-thymine DNA (dT_40_) and a multivalent R/G-rich polypeptide [(RGRGG)_5_] as a model system to mimic the heterotypic SG microenvironment^31^ (**Fig. 1a**). An advantage of this multi-component condensate system is that it offers precise control over condensate material properties through peptide sequence design^31^ and nucleic acid chain length^32^ (**Fig. 1a, Tables S1-S3**). To probe condensate impact on protein fibril formation, we tested a number of different fibril-forming protein clients, including WT A1-LCD, its disease variant D262V, and enhanced fibril-forming variants of Tau and FUS denoted as SynTag-Tau^33^ and enhanced FUS-LCD (eFUS-LCD; see **Supplementary Note 1** for details), respectively. These fibril-forming protein clients partition favorably into the SG-like condensates (**Fig. 1b**).

**Figure 1.**
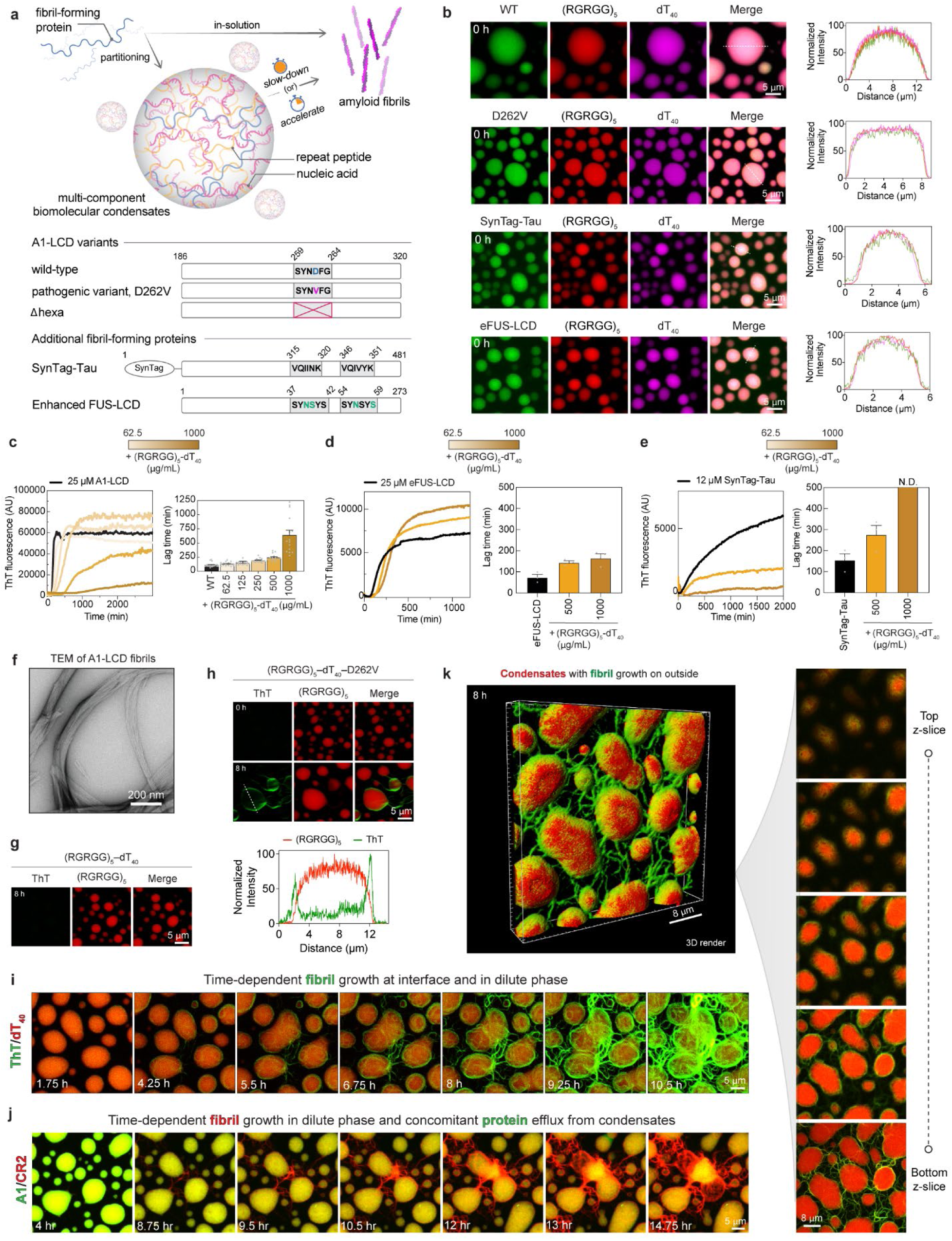
Multi-component biomolecular condensates robustly suppress fibril formation. **(a)** (top) Schematic of the experimental system designed to test whether multi-component peptide–nucleic acid condensates accelerate or suppress fibril formation kinetics. (bottom) Schematic highlighting the sequence architecture and amyloid-forming steric zipper motif (represented as a grey rectangle) of wild-type (WT) A1-LCD, encompassing residues 186 to 320 of the full-length hnRNPA1 protein, and variants of A1-LCD: D262V, a pathogenic variant, and Δhexa, a variant lacking the steric zipper. In addition, the sequence architecture and amyloid-forming steric zipper motifs of SynTag-Tau and Enhanced FUS-LCD (eFUS-LCD) are highlighted (cyan colored letters indicate amino acid substitutions that enhance fibrillization propensity). **(b)** Fluorescence images and corresponding line profiles show partitioning of individual fibril-forming proteins [with 250 nM Alexa488-labeled protein] into (RGRGG)_5_–dT_40_ condensates [with 250 nM of Alexa594-(RGRGG)_5_ and Cy5-dT_40_]. ‘0 h’ hereafter refers to 15 minutes after sample preparation. The display range of individual image panels was adjusted independently for optimal visualization. Kinetics of fibril formation, monitored by Thioflavin T (ThT) fluorescence, of **(c)** A1-LCD, **(d)** eFUS-LCD, **(e)** SynTag-Tau, either alone or in the presence of different concentrations of (RGRGG)_5_ and dT_40_. N.D. refers to ‘not determinable’. Baseline subtraction was applied to all the ThT curves. The ThT curves reported in (c) were generated using a smoothing window of 10 data points. Respective lag times (*t*_5%_) extracted from the ThT kinetics shown in (**c-e**) represent the time required to reach 5% of the plateau ThT fluorescence intensity. Lag times from replicate experiments (n ≥ 3) are shown along with the mean ± standard error of the mean (SEM). **(f)** Negative-stain transmission electron microscopy (nsTEM) image of A1-LCD fibrils from samples without condensates captured after 6 days of incubation at 20 °C with shaking. Confocal fluorescence images of 1 mg/mL (RGRGG)_5_–dT_40_ condensates, either without **(g)** or with 25 μM A1-LCD D262V **(h)** [with 250 nM Alexa594-(RGRGG)_5_] at the indicated time points after sample preparation. ThT was used to detect amyloid fibrils. **(i)** Time-lapse maximum intensity projection images of (RGRGG)_5_–dT_40_ condensates (visualized with 250 nM Cy5-dT_40_) containing A1-LCD D262V. Fibrils were detected using ThT fluorescence (green). See also **Video S1**. **(j)** Time-lapse images to monitor the progressive loss of A1-LCD D262V [with 250 nM Alexa488-D262V] from the interiors of (RGRGG)_5_–dT_40_ condensates. Fibrils were detected using CRANAD2 amyloid reporter (CR2). Also, see **Video S2**. **(k)** (left) 3D confocal Z-stack render of ThT-stained A1-LCD D262V fibrils and Cy5-dT_40_–labeled (RGRGG)_5_–dT_40_ condensates, imaged 8 hours after sample preparation. (right) Individual Z-slices corresponding to (k). Also, see **Video S3**.

To assess whether the condensates promote or suppress fibril formation, we progressively increased the concentrations of condensate-forming components, i.e., peptide and ssDNA, while keeping the total concentration of the fibril-forming protein constant. We monitored A1-LCD fibril formation through changes in the fluorescence intensity of Thioflavin T (ThT), a dye whose fluorescence intensity increases when it binds to amyloid fibrils^34^. Samples with peptide–ssDNA condensates without A1-LCD remained ThT-negative (**Fig. S1a**). In the absence of condensates, A1-LCD formed fibrils rapidly, as evidenced by a sigmoidal ThT curve with a lag time (*t*_5%_, which we defined as the time to reach 5% of the ThT plateau value) of ∼1.5 hours (**Fig. 1c**). Increasing the concentrations of peptide and ssDNA (i.e., titration of the condensate volume fraction) at a fixed concentration of A1-LCD increased the lag time progressively (**Fig. 1c**). Importantly, peptide or nucleic acid alone at concentrations that do not result in condensates were not able to suppress fibril formation (**Fig. S2**).

Condensates also suppressed fibril formation of A1-LCD D262V (**Fig. S1a, b**), SynTag-Tau, and eFUS-LCD to different protein-specific extents (**Fig. 1d, e**). We further verified that the condensate-mediated suppression effect applies to full-length hnRNPA1 (**Fig. S1c**). The presence of amyloid fibrils was confirmed using negative-stain transmission electron microscopy (nsTEM; **Fig. 1f**; **Fig. S1d**). Considering that ThT binding is subject to variation across different fibrillar species and amyloid polymorphs^35^, we did not quantitatively analyze the slopes and plateau values of the ThT curves; instead, we used the lag time as an empirical parameter to deduce the time required for the onset of fibril assembly across samples. Observation of the process using light microscopy confirmed that the formation of detectable mesoscale aggregates was increasingly suppressed (**Fig. S1e, f; Supplementary Note 2.1**). Since the different fibril-forming proteins used in our experiments possess unique amyloid motifs that drive fibril formation (**Fig. 1a**; bottom), the shared suppression of fibril formation upon partitioning into the multi-component condensates suggests that this is a generalizable effect of the microenvironment of these condensates.

### Amyloid fibrils grow in the dilute phase

To determine whether amyloid assembly occurs within condensates or in the surrounding dilute phase, we used quantitative time-lapse confocal fluorescence microscopy and monitored the spatial distribution of fluorescently labeled RGRGG-repeat peptide and the ThT signal in heterotypic condensates. In the absence of fibril-forming protein, the condensates did not show any ThT signal above the background level (**Fig. 1g**). Samples containing condensates and A1-LCD D262V after 8 hours of incubation resulted in a prominent ThT signal at the perimeter of condensates, and ThT-positive fibrillar structures extended into the dilute phase (**Fig. 1h**). We and others recently demonstrated that interfaces of homotypic protein condensates can promote fibril nucleation^21–24,33,36,37^. When monitoring the spatiotemporal distribution of ThT fluorescence in these multi-component condensates, we observe as early as two hours after sample preparation that ThT fluorescence increased at the interfaces of condensates, but not in the condensate interiors, suggesting that nucleation may also be promoted at the interfaces of multi-component condensates (**Fig. 1i, Video S1**). By imaging the distribution of fluorescently labeled D262V molecules as a function of time, we observe migration of protein molecules from the interior to the dilute phase, seemingly towards clusters of amyloid fibrils (**Fig. 1j, Video S2**). The condensates remain stable despite the efflux of A1-LCD molecules (**Fig. S1g**).

To test whether any fibrils were detectable in the dense phase, we performed confocal Z-stack imaging, which revealed that the fibrils localized in the dilute phase and in the interface (**Fig. 1k**; **Video S3**). Utilizing the amyloid reporter dye CRANAD2 and Alexa488-labeled A1-LCD D262V simultaneously, we confirmed that the fibrillar structures at the condensate perimeter are indeed formed by A1-LCD (**Fig. S1h**). Overall, our observations suggest that A1-LCD fibrils do not form in condensate interiors. These findings contrast with the prevailing view that fibril formation occurs in condensates owing to the high local concentration of fibril-forming protein within them ^38,39^, but are in agreement with studies that used homotypic condensates of A1-LCD variants^21^ and Tau^33^, which showed that condensates suppress fibril formation and that fibrils are nucleated at interfaces and form in the dilute phase.

### Condensates with higher viscoelasticity dampen fibril formation kinetics more effectively

Next, we sought to decipher the role the physical properties of condensates play in dampening fibril formation kinetics. Recent reports show that single and multi-component biomolecular condensates that form via phase separation of associative multivalent polymers are network fluids with sequence-encoded viscoelastic material properties^25,32,40,41^. An appealing feature of the synthetic peptide–nucleic acid condensates is their programmable viscoelasticity^31,42^. Subtly altering the sequence of the (RGRGG)_5_ polypeptide while keeping the ssDNA or RNA component fixed tunes condensate viscoelasticity over three orders of magnitude^31^. Substituting arginine with lysine or proline residues or glycine with proline residues attenuates condensate viscoelasticity. Conversely, substituting the second arginine with a tyrosine residue strengthens condensate viscoelasticity.

First, to directly determine the frequency-dependent linear viscoelastic moduli of the three-component peptide–ssDNA–A1-LCD condensates, we used passive microrheology with optical tweezers (pMOT) (**Fig. 2a**)^31^. The rank order of viscoelasticity for these condensates, determined by frequency-dependent storage (G’) and loss (G”) moduli and their crossover frequency (the inverse of which is the terminal relaxation timescale, *τ_M_*, of the condensate viscoelastic network) was the same as in the two-component peptide–ssDNA systems we previously reported^31,32^ (**Fig. 2b, c**). Specifically, condensates scaffolded by RGPGG- and KGKGG-repeat peptides were least viscoelastic, followed by those with RPRPP- and RGRGG-repeat peptides, while RGYGG-repeat peptide formed the most viscoelastic condensates with ssDNA (**Table S4**). This trend is reflected by (i) a progressive decrease in crossover frequency from ∼ 39 Hz to ∼ 1.0 Hz, with condensates in the order (KGKGG)_5_ > (RGPGG)_5_ > (RPRPP)_5_ > (RGRGG)_5_ > (RGYGG)_5_, and (ii) a monotonic increase in G’ and G” at a constant frequency (1.0 Hz), with the order (KGKGG)_5_ < (RGPGG)_5_ < (RPRPP)_5_ < (RGRGG)_5_ < (RGYGG)_5_. Notably, we find that the viscosities of the three-component condensates were consistently higher than those of the two-component peptide–ssDNA systems (**Table S5**).

**Figure 2.**
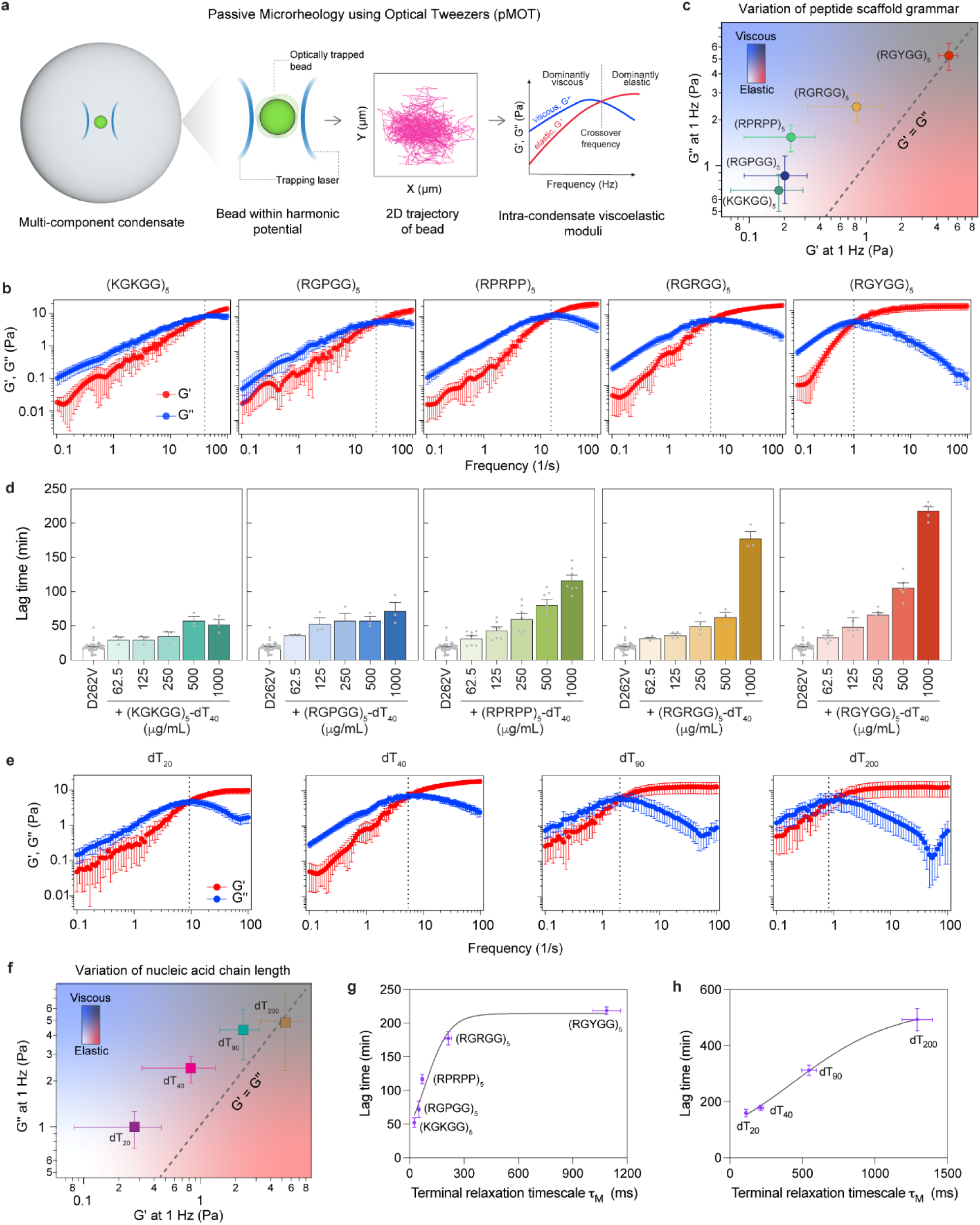
Higher viscoelasticity of multi-component biomolecular condensates prolongs lag times for fibril formation. **(a)** Schematic of passive microrheology using optical tweezers (pMOT) assay for measuring frequency-dependent viscous and elastic moduli of condensates. **(b)** Dynamical moduli for repeat peptide–nucleic acid condensates containing WT A1-LCD, wherein only the identity of the peptide component is varied. G′ and G′′, measured using pMOT, are the frequency-dependent storage and loss moduli, respectively. The data is represented as mean ± standard deviation (SD). The dashed line represents the crossover line. **(c)** Material state diagram of peptide–nucleic acid–WT A1-LCD condensates wherein the peptide sequence grammar is modulated. The data is represented as mean ± SD. **(d)** Lag times (*t*_5%_) extracted from the ThT kinetics corresponding to **Fig. S3d** for A1-LCD D262V either alone or in the presence of multi-component condensate systems, and from replicate experiments (n ≥ 3) are shown along with the mean ± SEM. **(e)** Dynamical moduli for repeat peptide–nucleic acid condensates containing WT A1-LCD, wherein only the length of the nucleic acid component is varied. The data is represented as mean ± SD. The plot corresponding to the ‘dT_40_’ condition is reproduced from that of ‘(RGRGG)_5_’ reported in panel (b). **(f)** Material state diagram of peptide–nucleic acid–WT A1-LCD condensates wherein the length of the nucleic acid component is varied. The data is represented as mean ± SD. **(g)** Lag times (*t*_5%_) for the highest peptide–ssDNA concentrations extracted from the ThT kinetics shown in (d) and from replicate experiments (n ≥ 3) are represented as mean ± SEM as a function of terminal relaxation timescale of the multi-component condensate system. **(h)** Lag times (*t*_5%_) for A1-LCD D262V fibril formation across peptide–ssDNA condensate systems [extracted from the ThT kinetics shown in **Fig. S5**; from replicate experiments (n ≥ 3)], wherein nucleic acid length was varied, are represented as mean ± SEM as a function of terminal relaxation timescale of the multi-component condensate system. The data in (g) and (h) are fitted with a logistic growth model as a guide to the eye.

Next, we compared the ability of these peptide–ssDNA–A1-LCD condensates to suppress fibril formation of WT A1-LCD (**Fig. S3a-c**) and of the pathogenic D262V variant (**Fig. 2d, Fig. S3d, e**). For each type of condensate, increasing peptide and ssDNA concentrations progressively increased lag times on a system-specific timescale (**Fig. S3b, d**). Notably, at a fixed peptide–ssDNA concentration, lag time increased with condensate viscoelasticity (**Fig. S3c, e**). The lag time for fibril formation of the pathogenic variant D262V in the presence of highly viscoelastic (RGYGG)_5_–dT_40_ condensates (viscosity, η, 95 ± 5 Pa·s; terminal relaxation timescale, *τ_M_*, at which G’/G” = 1, is 1087 ± 77 ms) is at least ∼4 times longer relative to that of the least viscoelastic condensates, (KGKGG)_5_–dT_40_ (η = 2.4 ± 0.4 Pa·s; *τ_M_* = 26 ± 3 ms) (**Fig. S3e; Fig. 2g**). Similarly, increasing the viscoelasticity of peptide–ssDNA condensates resulted in progressive dampening of WT A1-LCD fibril formation (**Fig. S3c**).

SGs sequester untranslated RNA and are therefore classified as ribonucleoprotein (RNP) granules^4^. Therefore, we next assessed whether condensates formed by repeat peptides and rU_40_ RNA, in place of ssDNA, similarly suppress fibril formation. Indeed, analogous to the samples containing peptide–ssDNA condensates, the lag times for fibril formation increase with increasing peptide–RNA concentrations (**Fig. S4a-d**). At a fixed peptide–RNA concentration, increasing condensate viscoelasticity similarly dampens fibril formation kinetics of the pathogenic D262V variant (**Fig. S4d**). A range of simple RNP granule mimics can therefore robustly suppress fibril formation of A1-LCD.

Because the material properties of the condensate systems employed here are tuned by using peptides with different chemical makeup (**Fig. 2b**), the observed changes in A1-LCD fibril formation kinetics may be influenced by peptide−A1-LCD interactions rather than condensate viscoelasticity alone. To test this possibility, we used an orthogonal approach to tune the viscoelasticity of multi-component condensates, achieved by increasing the length of the condensate-scaffolding nucleic acid without changing the peptide or nucleic acid chemistry^32^. By varying the poly-dT chain length from 20 to 200 nucleobases, while keeping the peptide sequence grammar constant (i.e., RGRGG repeats), the viscoelastic spectra of the resulting condensate systems spanned a range of terminal relaxation timescales of *τ_M_* = 111 ± 4 ms to *τ_M_* = 1291 ± 106 ms (**Fig. 2e**). Furthermore, the linear viscoelastic properties of these condensates show a steady increase in G’ and G” as a function of increasing chain length (**Fig. 2f**). When we tested A1-LCD D262V fibril formation in the presence of these heterotypic condensates, we observed that condensates with longer ssDNA strongly prolong the lag time of fibril formation, similar to what was observed earlier with changes to the peptide grammar (**Fig. 2g, h**; **Figs. S5, S6**). Notably, consistent with our previous observations, the peptide and ssDNA constructs used here were unable to suppress fibril formation when tested individually at concentrations that did not result in condensate formation (**Fig. S2**). Overall, these results demonstrate that differences in condensate physical properties, such as viscoelasticity, correlate with the suppression of fibril formation by condensate-partitioning proteins, although the mechanistic basis underlying this relationship remains to be established.

### Dilute-phase protein concentration does not explain lag time across multicomponent condensates

A simple explanation for the observed decrease in fibril formation is that the fibril-forming protein is depleted from the dilute phase and progressively sequestered into condensates as peptide and ssDNA concentrations increase. To test this hypothesis, we determined dilute phase concentrations (*c*_dilute_) of A1-LCD in the presence of condensates using analytical HPLC measurements^43^ (**Fig. 3a**; *see* Methods). At 62.5 μg/mL (RGRGG)_5_ peptide and dT_40_ ssDNA, *c*_dilute_ of A1-LCD was 5.4 μM, whereas it was reduced to 1.6 μM at 125 μg/mL peptide/ssDNA concentrations (**Fig. 3b**). *c*_dilute_ of A1-LCD was below the detection limit at greater than 125 μg/mL peptide/ssDNA concentrations. These data show that A1-LCD is indeed progressively depleted from the dilute phase, thereby protecting against A1-LCD fibril formation (**Fig. S1b**). We note that the saturation concentration for fibril formation, *c*_sf_, for WT A1-LCD is ∼0.3 μM in these buffer conditions (**Fig. 3b**). The presence of condensate components does not change *c*_sf_. However, fibril formation below *c*_sf_ may occur in the ternary system because the presence of peptide–ssDNA condensates creates spatially heterogeneous microenvironments that can alter local monomer activities and, consequently, the effective *c*_sf_. Of note, TEM analysis of the samples reveals assemblies that are shorter than typical amyloid fibrils with seeding capacity to promote fibril formation, albeit markedly less potent than bona fide A1-LCD seeds (**Fig. S7; Supplementary Note 2.2**).

**Figure 3.**
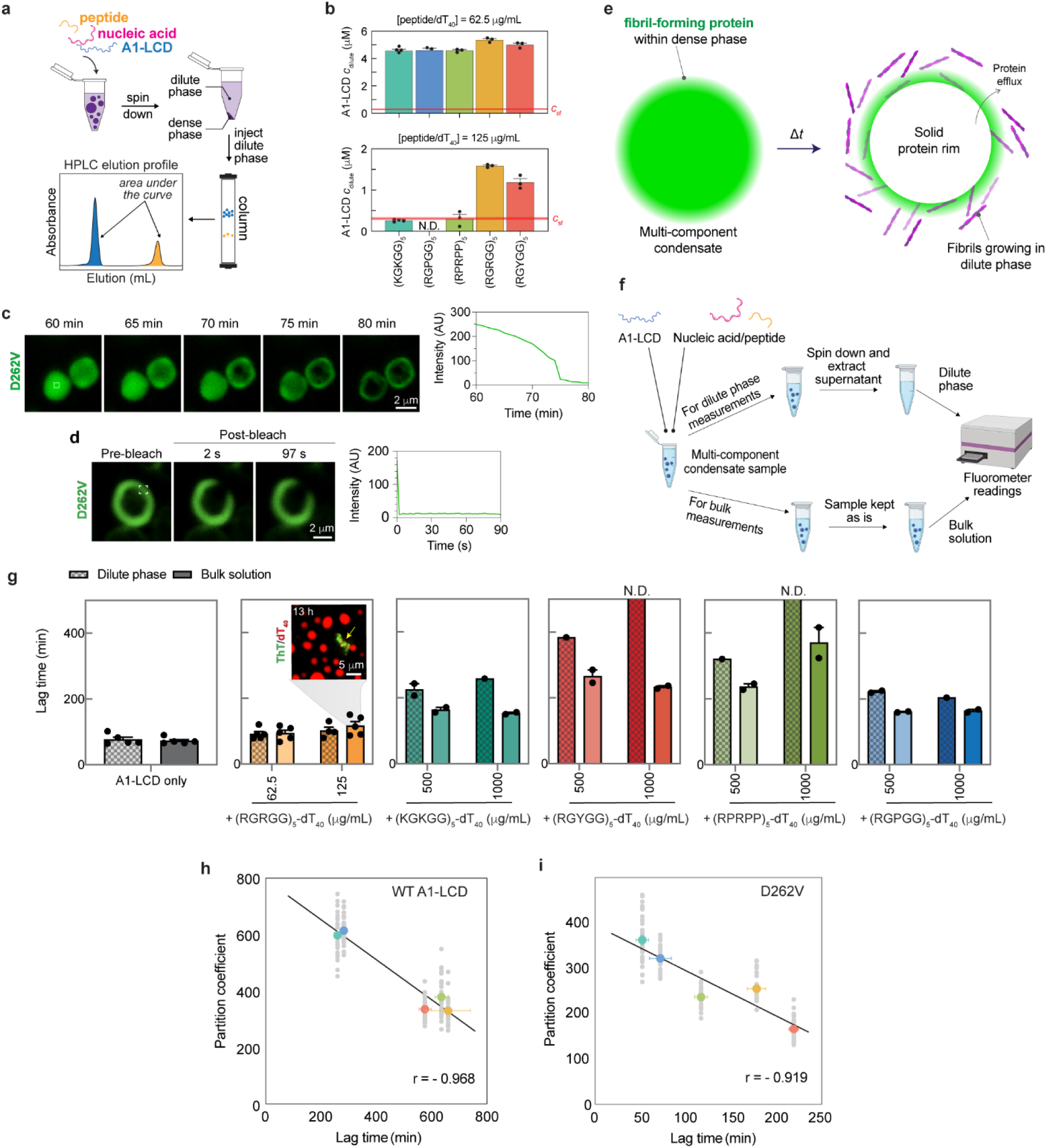
Reconciling the roles of dilute phase and condensate interfaces in amyloid fibril formation. **(a)** Schematic of analytical HPLC assay^43^ to determine the dilute phase concentration (*c*_dilute_) of A1-LCD in samples containing protein–nucleic acid condensates. **(b)** *c*_dilute_ of WT A1-LCD in samples containing peptide–dT_40_ condensates at two different volume fractions. The data is represented as mean ± SEM, and the points indicate individual measurements. Red dashed lines indicate the *c*_sf_ value for WT A1-LCD. **(c)** Time-lapse images to monitor the progressive loss of A1-LCD D262V (visualized with 250 nM Alexa488-D262V) from the interiors of (RGPGG)_5_–dT_40_ condensates. The time-dependent fluorescence intensity change in the area of the white box is also shown. See also **Video S4**. **(d)** Fluorescence recovery after photobleaching (FRAP) profile of an A1-LCD corona (visualized using 250 nM Alexa488-D262V) from a (KGKGG)_5_–dT_40_ condensate. This sample was imaged 1.5 hours after sample preparation. **(e)** Schematic summarizing our experimental observation from **(c)**: ThT-positive protein rim formation coupled with A1-LCD efflux from the peptide–dT_40_ condensates. **(f)** Schematic showing fractionation of dense and dilute phases for ThT fluorescence readings. **(g)** Estimated lag times from measurements of ThT kinetics. Inset depicts a representative fluorescence image of 125 μg/mL (RGRGG)_5_–dT_40_ condensates with 25 μM A1-LCD [with 250 nM Cy5-dT_40_] at the indicated time point after sample preparation. ThT was used to detect amyloid fibrils. The yellow arrow indicates amyloid fibrils. N.D. refers to ‘not determinable’. **(h, i)** Correlation analysis between partition coefficients of WT A1-LCD (**Fig. S8c**) and D262V (**Fig. S8d**) in different protein–nucleic acid condensates (individual measurements are shown along with the mean) versus the lag time (*t*_5%_ as shown in **Fig. S3c, e**; the mean is shown along with the SEM). A straight line shows the best fit for the correlation, and the Pearson correlation coefficient (r) is indicated.

Why do condensates with higher viscoelasticity suppress fibril formation more effectively? One possibility is that condensates that suppress fibril formation recruit more A1-LCD at a fixed peptide–ssDNA concentration. Across all peptide–ssDNA condensates tested, increasing peptide and ssDNA concentrations progressively depleted A1-LCD from the dilute phase (**Fig. 3b**). Notably, *c*_dilute_ was highest for (RGRGG)_5_- and (RGYGG)_5_-based condensate systems, which are the strongest suppressors of A1-LCD fibril formation. Conversely, samples with the lowest A1-LCD dilute phase concentrations, such as those with (RGPGG)_5_–ssDNA condensates, in which the A1-LCD dilute phase concentration was too low to be measurable by analytical HPLC, showed the shortest lag times. Therefore, although dilute-phase concentration is a key determinant of fibril growth, it does not fully explain the differences in lag times observed across different peptide–ssDNA condensates with distinct physical properties.

### Extent of protein sequestration defines the role condensate interfaces play in nucleation

Condensate interfaces represent vulnerabilities because they can promote nucleation^21–24^. Indeed, in condensates that were the least effective at lengthening lag times, including those formed with (RGPGG)_5_, we observed migration of A1-LCD to a rim on a timescale of minutes (**Fig. 3c, Video S4**), as opposed to the timescale of hours in condensates scaffolded by (RGRGG)_5_ (**Fig. 1j**; **Fig. S1g, Video S2**). Notably, after protein loss from the condensate interior, a corona enriched in A1-LCD D262V remained (**Fig. 3d, e**). Fluorescence recovery after photobleaching (FRAP) measurements showed low fluorescence recovery of the D262V corona, suggesting that amyloid fibrils at the condensate interface result in the formation of a solid-like interfacial shell (**Fig. 3d**), reminiscent of observations of solid shells around FUS condensates^23,37^. These observations support the view that interfaces can promote nucleation. But are condensate interfaces always the dominant sites for fibril nucleation, or does the dilute phase dominate nucleation under some conditions? To answer this question, we introduced a fractionation approach to separate the dilute from the dense phase using centrifugation (**Fig. 3f**) and compared time-dependent ThT intensities for the fractionated dilute phase and for the total sample containing dilute and dense phase (called “bulk samples” henceforth) for samples with (RGRGG)_5_-dT_40_ condensates and WT A1-LCD. While the ThT plateau values reached in bulk samples are higher due to the larger total concentration of fibril-forming protein, we found that the estimated lag times do not differ substantially between dilute phase–only versus bulk samples at concentrations up to 125 μg/mL (RGRGG)_5_–dT_40_ (**Fig. 3g; Fig. S8a**). These observations suggest that condensate interfaces do not contribute strongly to nucleation under these conditions.

Given that the (RGRGG)_5_-based systems substantially dampen fibril formation at high concentrations (**Fig. 1c**), we also assessed the behavior of A1-LCD with the other peptide systems. At high peptide–dT_40_ concentrations (i.e., at 500 and 1000 μg/mL), the lag phases were shorter in bulk than in dilute phase–only samples (**Fig. 3g; Fig. S8b**), suggesting that condensate interfaces catalyze fibril nucleation under these conditions. Overall, comparing the ThT kinetics of dilute phase–only and bulk samples at low and high peptide/ssDNA concentrations across different peptides showed consistent trends: At low peptide/ssDNA concentrations, the lag phases were indistinguishable in the presence and absence of condensates. At high peptide/ssDNA concentrations, the presence of condensates shortened the lag phases, pointing to nucleation at interfaces. The relevant difference between these conditions is likely that the dilute phase concentration of A1-LCD is below *c_sf_* at the high peptide/ssDNA concentrations (**Fig. 3b**). Our observations suggest that the presence of condensates promotes fibril nucleation under conditions where *c_dil_* < *c_sf_* for A1-LCD, likely because the dilute phase concentrations are too low to mediate effective nucleation. Consistent with this inference, we find that samples containing 125 μg/mL (RGRGG)_5_/dT_40_ and A1-LCD, wherein *c*^A1−LCD^ > *c_sf_* (**Fig. 3b**), fibrils formed in the dilute phase without any detectable ThT fluorescence at interfaces (inset shown in **Fig. 3g**). In conclusion, interfaces dominate fibril nucleation under conditions wherein the dilute phase is not conducive to nucleation.

Under conditions wherein condensate interfaces promote fibril nucleation, which factors determine the resulting lag phases? We wondered whether A1-LCD concentrations at interfaces, as opposed to concentrations in the dilute phase, could be a decisive factor. We determined the partition coefficients of WT A1-LCD or D262V into peptide–ssDNA condensates (**Fig. S8c, d**) at early time points, which we take as a proxy for A1-LCD concentrations at condensate interfaces (**Supplementary Note 3**). Notably, the partition coefficients show an inverse correlation with the lag time (**Fig. 3h, i**), which reports on nucleation rates. This means that condensates exhibiting higher A1-LCD partitioning also showed shorter fibril nucleation lag times, and vice versa. Hence, high densities of fibril-forming proteins at interfaces represent vulnerabilities in disease processes.

### Kinetic modeling unmasks the role of protein density at condensate interfaces

Leveraging our quantitative experimental observations, we developed a compartmental kinetic model that enables dissection of the effects of coupling condensate-mediated sequestration to amyloid assembly (**Fig. 4a**; see **Supplementary Note 4** for model description, parameters, and numerical implementation). We consider that the fibril-forming protein partitions into three reaction compartments: condensate interior (with concentration *c_c_*), the surrounding dilute phase (*c_d_*), and a thin interfacial layer (*c_i_*) that we treat as a distinct compartment whose loading follows a Langmuir isotherm with affinity *K_int_* and finite capacity *c_i_^max^*. The condensate phase is modeled as a single spherical droplet with an area-to-volume ratio *A*/*V*, which explicitly enters into appropriate rate expressions and couples interfacial amyloid assembly kinetics to droplet geometry. Exchange between condensate and dilute phases is set by the influx rate (*k_in_*) into the condensate and the efflux rate (*k_out_*) out of the condensate, with the partition coefficient defined as *K_part_* = *k_in_*/*k_out_*. Fibril nucleation and growth follow a standard amyloid chemical-kinetics framework: nucleation can occur in the dilute phase and at the condensate interface^44,45^, and we have explored scenarios with and without dilute phase nucleation (see **Supplementary Note 4**; **Fig. S9**). Primary nucleation at the interface scales as 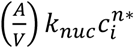, and existing fibrils template further nucleation via a secondary, mass-dependent reaction pathway 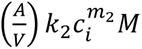. Here, *n\** and *m*_2_ are the reaction orders of primary and secondary nucleation, respectively, with respect to the interfacial protein concentration *c_i_*; *M* denotes the existing fibril mass; and the *k* terms are the corresponding rate constants. Based on our previous work^46^, the model prescribes that elongation, which occurs in the dilute phase, draws monomers from the dilute pool at a rate *k*_+_*c_d_N*, so that the locations of nucleation and fibril growth occur in separate compartments. These model choices result in the following dichotomy: Condensates simultaneously concentrate monomers, accelerate nucleation at their interfaces, and throttle elongation by efflux of monomers into the dilute phase. From the resulting trajectories, we track the total fibril mass *M*(*t*) and the lag time *t*_5%_ defined as the time at which *M*(*t*) reaches 5% of its plateau value.

**Figure 4.**
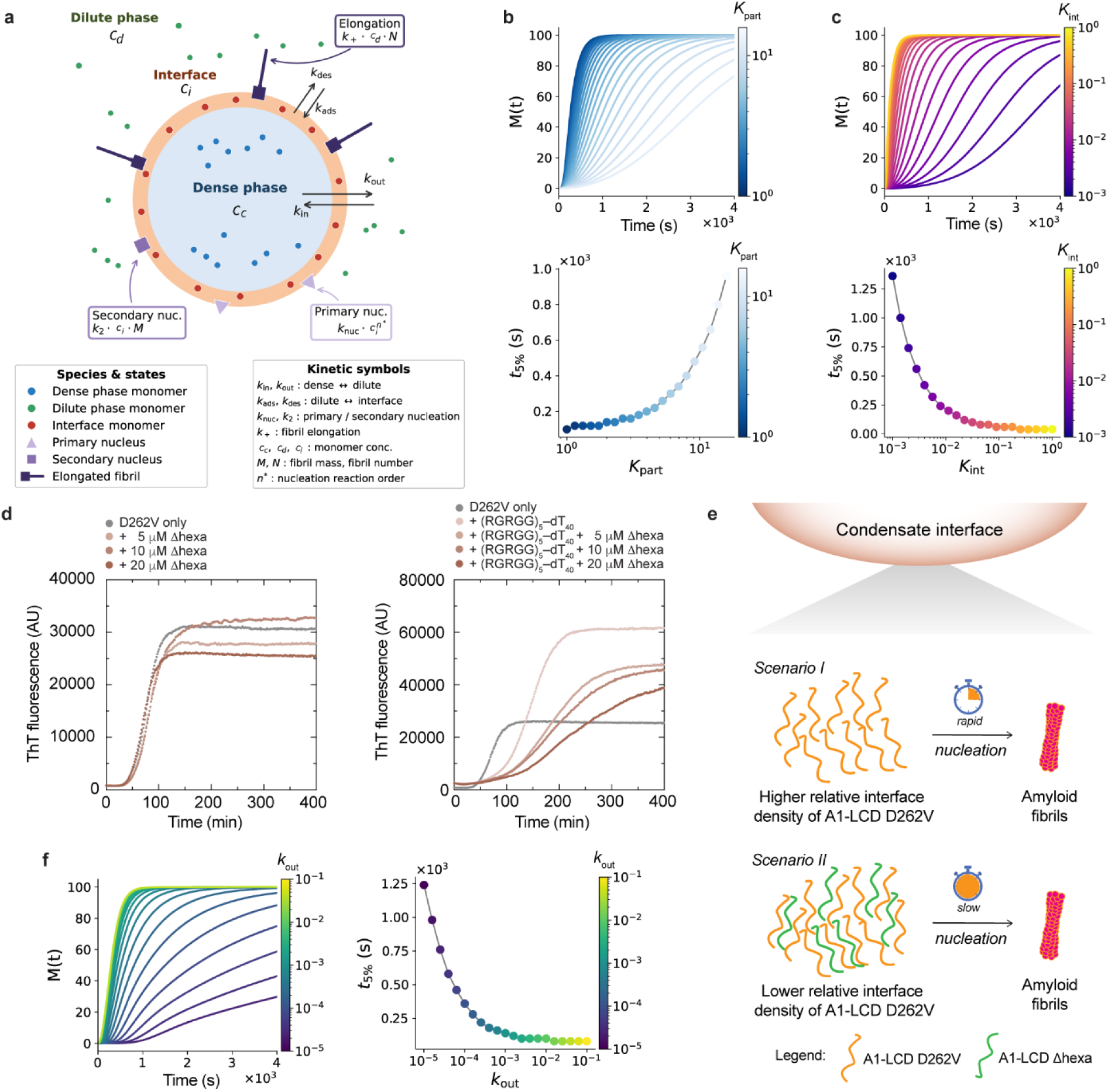
Quantitative modeling reveals key mechanisms of condensate–mediated suppression of amyloid assembly. **(a)** Schematic illustrating the quantitative kinetic model of fibril assembly in the presence of condensates and a brief overview of the relevant parameters. **(b)** Fibril assembly kinetics [as total fibril mass *M*(*t*) and estimated lag times *t*_5%_ for fibril assembly] as a function of the fibril-forming protein sequestration into the dense phase (*K_part_*). **(c)** Total fibril mass *M*(*t*) and estimated lag times *t*_5%_ for fibril assembly as a function of fibril-forming protein enrichment at the condensate interface (*K*_int_). **(d)** Kinetics of fibril formation monitored using ThT fluorescence of 25 μM A1-LCD D262V either alone or with increasing concentrations of A1-LCD Δhexa either in the absence (left) or presence (right) of (RGRGG)_5_–dT_40_ condensates. **(e)** Schematic representing the observed effect of A1-LCD interface density on fibril formation, either in the absence or presence of A1-LCD Δhexa. **(f)** Fibril assembly kinetics upon titration of fibril-forming protein efflux from dense phase and into the dilute phase (*k_out_*), as well as the estimated lag times for fibril assembly.

Using this framework, we first asked whether the model captures how sequestration of monomers within the dense phase impacts fibril formation. Consistent with our experiments (**Fig. 1c-e**; **Fig. 2d; Fig. S1**), the model shows that increasing partitioning into the condensate and the associated depletion of dilute phase protein lengthens *t*_5%_ and lowers the resulting fibril mass (**Fig. 4b**).

Next, we introduced the possibility of nucleation in the dilute phase using the same nucleation kinetics to determine how the concentration of fibril-forming protein at the interface affects the kinetics and location of fibril assembly. In agreement with our experimental results (**Fig. 3g**), kinetic modeling suggests that interfaces strongly increase fibril nucleation at low dilute phase concentrations, where bulk nucleation is ineffective (**Fig. S9**). Furthermore, the model predicts that the lag time shortens exponentially with increasing enrichment of fibril-forming protein at condensate interfaces, parameterized by *K*_int_ (**Fig. 4c**; **Fig. S9; Supplementary Note 4**). Because A1-LCD is homogeneously distributed within condensates at early time points, the experimentally measured partition coefficient also reflects the local protein density at the interface (**Supplementary Note 4.1**); Under these conditions, the measured partition coefficient can therefore be interpreted as a proxy for the model parameter *K*_int_ (**Fig. 4c**), rather than for the bulk partition coefficient *K_part_* (**Fig. 4b**), which governs protein sequestration into the condensate interior. Importantly, *K*_int_ and *K_part_* act oppositely on the lag time, i.e., stronger interior partitioning lengthens it by depleting the dilute phase (**Fig. 4b**), whereas higher interface enrichment shortens it by accelerating nucleation (**Fig. 4c**). Thus, the inverse correlation between the measured partition coefficient and nucleation lag time (**Fig. 3h, i**) is consistent with the *K*_int_ −dependent regime of the model.

Based on these insights, we hypothesized that dilution of A1-LCD density at the condensate interface through the addition of additional protein species may extend lag times and suppress fibril formation further. Given that SGs contain many different RBPs with PLDs, we tested the effect of adding a PLD variant that lacks the propensity to form fibrils but retains the ability to partition into peptide–ssDNA condensates. To this end, we chose a variant of A1-LCD termed Δhexa, in which the primary steric zipper (^259^SYNDFG^2^^64^), which includes the site of the disease variant, is deleted (**Fig. 1a**)^30^. When A1-LCD Δhexa was titrated into dilute solutions containing only A1-LCD D262V without peptide–ssDNA condensates, the kinetics of fibril formation remained unaffected (**Fig. 4d**, *left*). By contrast, the addition of increasing concentrations of A1-LCD Δhexa to condensate samples consisting of A1-LCD D262V, (RGRGG)_5_, and dT_40_ progressively prolonged the lag time (**Fig. 4d**, *right*), although the partition coefficient of D262V in condensates did not change substantially (**Fig. S10a, b**). Video particle tracking-based nanorheology measurements revealed that the Δhexa variant does not alter condensate viscosity (**Fig. S10c**), suggesting that the increase in the lag time is mediated by the altered interfacial protein density. These results support our conclusion that the relative density of fibril-forming protein at the condensate interface is a key determinant of the fibril assembly lag time (**Fig. 4e**). Experiments and modeling therefore support the existence of two regimes for nucleation: (i) At high dilute-phase concentrations, nucleation in the dilute phase is effective and dominates; and (ii) With strong sequestration of protein into condensates, resulting in low dilute phase concentrations, the interfaces are the only place where nucleation can occur.

### Protein efflux down a chemical potential gradient governs fibril growth in the dilute phase

Once fibril nuclei are present, our experimental system comprises three coexisting phases: the dilute phase, the condensate, and fibrils. If the fibrils, which are localized at the interface or in the dilute phase, represent the phase with the lowest chemical potential (*µ*), i.e., are the most stable, then fibril growth will be driven by chemical-potential gradient-assisted mass transport of A1-LCD, analogous to Ostwald ripening^47^. This gradient therefore drives monomer efflux from the condensate into the dilute phase and ultimately toward fibril ends. Earlier experimental observations using time-resolved fluorescence microscopy alluded to the prominent role of protein efflux from the condensate interior during fibril growth (**Fig. 1j, 3c**; **Fig. S1g**). Furthermore, time-lapse microscopy images with condensates formed with ssDNA of different lengths reveal the following notable feature of highly viscoelastic condensates: Condensates with the longest ssDNA displayed CRANAD2-positive rims for many hours without obvious signs of microscopically detectable fibrils in the dilute phase, whereas fibrils were detected around condensates with shorter ssDNA within a few hours (**Fig. S11a**). The differences in lag times were not caused by different protein densities at interfaces because the partition coefficients were similar across the different nucleic acid lengths (**Fig. S11b, c**). Instead, the observations suggest that fibril nucleation had occurred, yet fibrils did not effectively grow in the dilute phase of heterotypic condensates scaffolded by longer ssDNA (**Fig. S11**), potentially because of slower efflux of A1-LCD from more viscoelastic condensates, which would limit fibril growth in the dilute phase.

To test this idea, we used the kinetic model to interrogate how the condensate efflux rate *k_out_* impacts nucleation and elongation across compartments (**Fig. 4f**; **Supplementary Note 4**). Lowering *k_out_*, while keeping *K_part_* constant, leaves the interfacial concentration *c_i_* largely intact, since it is set by the Langmuir equilibrium with the dilute phase, so primary nucleation at rate 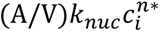 describes generation of nuclei at the condensate interface. Elongation, however, proceeds from the dilute phase monomer population through *r_elonee_* = *k*_+_*c_d_N*, where *N* is the total concentration of fibril-forming protein. Slower efflux is unable to replenish *c_d_*, and the production of fibril mass becomes slower, even if *N* is appreciable. Consequently, *M*(*t*) plateaus well below saturation, and *t*_5%_ rises sharply as *k_out_* decreases (**Fig. 4f**), reproducing the stalled fibril growth phenotype observed in viscoelastic condensates formed with long ssDNA (**Fig. S11a**). The model thus predicts that viscoelastic condensates decouple nucleation from growth by accumulating nuclei at the interface (consistent with the CRANAD2-positive rims [**Fig. S11a**] and short non-fibrillar assemblies observed by TEM [**Fig. S7a**]) while suppressing dilute-phase fibril growth. Furthermore, condensates with similar A1-LCD partitioning (**Fig. S11c**) but higher viscoelasticity are predicted to exhibit reduced efflux of A1-LCD into the dilute phase, thereby limiting the availability of monomers for the growth of ThT-positive fibrils.

### Condensate viscoelasticity determines the efflux rate of fibril-forming protein

To directly track the kinetics of A1-LCD recruitment into and efflux from peptide–ssDNA condensates of varying viscoelasticity, we developed a real-time quantitative assay that takes advantage of microfluidics, correlative confocal fluorescence microscopy, and a dual trap laser tweezer system (**Fig. 5a**). Briefly, we optically trapped a single peptide–ssDNA condensate under laminar flow in one channel [Channel 1 (Ch1)] of a microfluidic chamber and moved it at a controlled velocity to an adjacent channel containing buffer only (Ch2), and eventually to a third adjoining channel (Ch3) containing fluorescently labeled A1-LCD D262V under laminar flow. Continuous fluorescence imaging allowed us to capture the kinetics of client recruitment to the optically trapped condensate while in Ch3. The condensate was kept in this channel for 100 s to measure the extent and speed of client recruitment. Subsequently, the condensate was moved back to the buffer channel (Ch2) to measure the extent and speed of client efflux (**Fig. 5a**).

**Figure 5.**
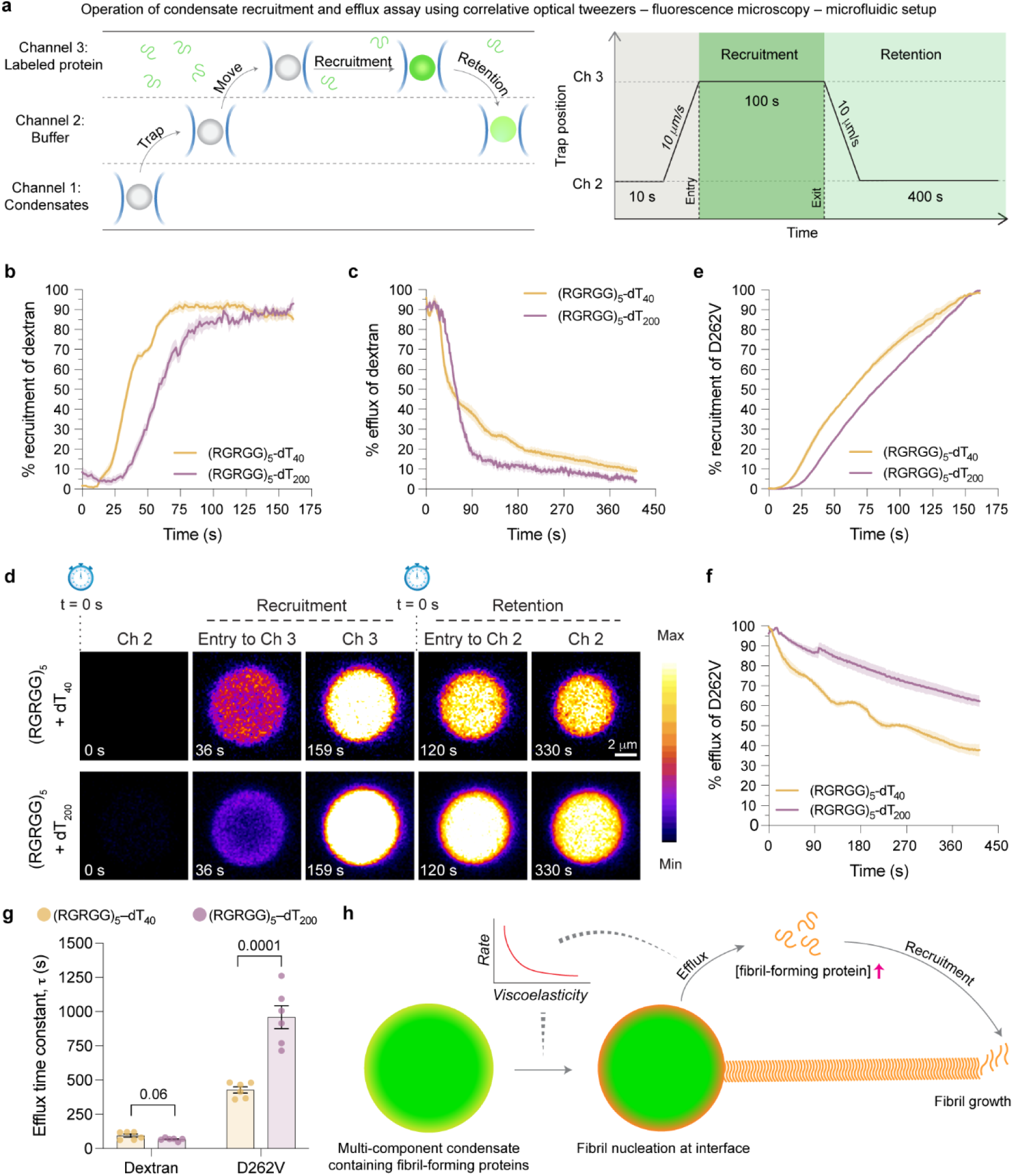
Higher condensate viscoelasticity slows down the efflux of fibril-forming protein and limits fibril growth. **(a)** (top) Schematic depicting the principle of the condensate recruitment and retention assay involving the simultaneous use of microfluidics, optical trapping, and fluorescence microscopy to determine the influx and efflux kinetics of fluorescently labeled client molecules into and out of multi-component condensates. (bottom) Schematic of the trajectory followed by the optical trap in the microfluidic chamber across different channels (Ch). **(b)** Recruitment kinetics of TMR-labeled dextran-4.4K into (RGRGG)_5_–dT_40_ and (RGRGG)_5_–dT_200_ condensates reported as mean ± SEM. **(c)** Retention kinetics of TMR-labeled dextran-4.4K in (RGRGG)_5_–dT_40_ and (RGRGG)_5_–dT_200_ condensates reported as mean ± SEM. **(d)** Time-lapse fluorescence images of Alexa488-labeled D262V showing partitioning at different stages of the recruitment and retention assay, in (RGRGG)_5_–dT_40_ and (RGRGG)_5_–dT_200_ condensates. Also see **Videos S5** and **S6**. The displayed maximum intensity threshold value of the image panels of (RGRGG)_5_–dT_40_ and (RGRGG)_5_–dT_200_ was adjusted independently for optimal visualization. **(e)** Recruitment kinetics of Alexa488-labeled D262V into (RGRGG)_5_–dT_40_ and (RGRGG)_5_–dT_200_ condensates reported as mean ± SEM. **(f)** Retention kinetics of Alexa488-labeled D262V in (RGRGG)_5_–dT_40_ and (RGRGG)_5_–dT_200_ condensates reported as mean ± SEM. **(g)** Efflux time constants of either dextran-4.4K or D262V in (RGRGG)_5_–dT_40_ and (RGRGG)_5_–dT_200_ condensates, extracted from the corresponding retention kinetics by fitting to a stretched exponential decay. Statistical significance was determined by performing unpaired *t* tests between the highlighted pairs of data, with the estimated *p* value displayed above. The points indicate individual measurements from three independent replicates and are shown along with mean ± SEM. **(h)** Schematic highlighting the impact of condensate viscoelasticity in determining the rate of efflux of fibril-forming protein from condensate interiors, serving as a rate-limiting step in fibril growth.

We first tracked recruitment and retention of tetramethylrhodamine (TMR)-labeled dextran-4.4K, a non-specific client that favorably partitions into peptide–ssDNA condensates (**Fig. S11d**), in two condensate systems of distinct viscoelasticity. Recruitment of dextran-4.4K into (RGRGG)_5_–dT_40_ (*τ_M_* = 213 ± 19 ms) and (RGRGG)_5_–dT_200_ condensates (*τ_M_* = 1291 ± 106 ms) plateaued at ∼100 s (**Fig. 5b**; **Fig. 2h**). The recruitment kinetics into the dT_40_ condensate system with lower viscoelasticity were somewhat faster than into the dT_200_ condensates with higher viscoelasticity. Moving the condensate back to the buffer-only channel resulted in efflux of dextran on similar timescales for both types of condensates (**Fig. 5c**). Therefore, the efflux kinetics of dextran are not dependent on the bulk viscoelasticity differences of the condensates, indicating (i) the molecular size of dextran is lower than the correlation length or the average mesh size of the condensates^48,49^, and (ii) the client does not engage in dominant interactions with the viscoelastic network of peptide-ssDNA condensates.

Next, we performed the recruitment and retention assay with Alexa488-labeled A1-LCD D262V. Although A1-LCD D262V was recruited into both condensates on comparable timescales, recruitment was noticeably slower in the more viscoelastic condensates. (**Fig. 5d, e, Videos S5, S6**). By contrast, the protein efflux from highly viscoelastic (RGRGG)_5_–dT_200_ condensates was markedly slower than the efflux from less viscoelastic (RGRGG)_5_–dT_40_ condensates (**Fig. 5d, f**; **Videos S5, S6**). The efflux time constants (*τ*) extracted from time-dependent fluorescence decay curves indicate that the more viscoelastic condensates retain A1-LCD D262V molecules for a significantly longer period (*τ* ∼ 960 s) than the less viscoelastic condensates (*τ* ∼ 428 s) (**Fig. 5g**). Hence, the difference in the strength of the underlying condensate-spanning network, which manifests as condensate viscoelasticity, correlates with the A1-LCD efflux rate (**Fig. 5h**). Of note, the efflux assay performed here is an active measurement under flow, which greatly accelerates molecular exchange and therefore underestimates the timescale relevant to quiescent fibril-formation conditions lacking flow. This is illustrated by a comparison of **Videos S2** and **S4**, which shows that condensates with higher viscoelasticity exhibit efflux on the timescale of multiple hours, in contrast to the substantially shorter timescale observed for condensates with lower viscoelasticity. Furthermore, **Fig. S11a** qualitatively reveals passive protein efflux across condensate systems of different nucleic acid lengths, which underscores the difference in efflux timescales between active (**Fig. 5d**) and passive modes (**Fig. S11a**). Overall, these results show that multi-component condensates with higher viscoelastic network strength release fibril-forming proteins more slowly than the typical rate of protein incorporation into fibrils in the dilute phase, thereby creating a kinetic bottleneck in fibril assembly.

### Reconstituted stress granules protect against amyloid fibril formation

Our observations so far suggest three key features by which multi-component peptide–nucleic acid condensates influence the timescale of fibril formation: (1) Fibrils form in the dilute phase, and strong sequestration of the fibril-forming protein into condensates lowers its concentration in the dilute phase and slows down or even abrogates fibril growth. (2) Lower effective densities of the fibril-forming protein at the interfaces of multi-component condensates reduce nucleation and lengthen lag times. (3) Higher condensate viscoelasticity reduces the efflux of fibril-forming proteins into the dilute phase and limits fibril growth. To what extent are these observations generalizable to biologically relevant SG mimics?

Given that G3BP and poly-A tail–containing mRNA are essential for SG assembly and form the core network that recruits other RBPs^2^, we tested whether multi-component condensates consisting of G3BP1, poly-rA, and A1-LCD could similarly suppress A1-LCD fibril formation (**Fig. 6a**). Using fluorescence microscopy, we first confirmed that A1-LCD D262V strongly partitions into G3BP1–poly-rA condensates (**Fig. S12a**). We then used ThT fluorescence to monitor A1-LCD fibril formation both alone and in the presence of increasing concentrations of G3BP1 and poly-rA at a constant ratio and at different ratios. Higher concentrations of condensate-forming components indeed led to progressively longer lag times (**Fig. 6b**, **Fig. S12b**). Notably, we observed an increase in the fluorescence intensity of the amyloid dye CRANAD2 at condensate interfaces, consistent with the interpretation that fibril nucleation occurs at the interfaces of condensates. This is followed by redistribution of A1-LCD from the condensate interior to A1-LCD fibrils in the dilute phase and the interface over time (**Fig. S12c**). To probe whether reduced interface density can mitigate fibril nucleation, we titrated increasing concentrations of the A1-LCD Δhexa variant into D262V–G3BP1–poly-rA mixtures and found a progressive delay in fibril formation (**Fig. S12d**). Consistent with our findings in peptide–nucleic acid condensates, we did not observe typical fibrils in the presence of G3BP1–poly-rA condensates but rather small, irregular assemblies that may be accumulated nuclei or protofibrils (**Fig. S12e**). We note that these assemblies had lower seeding capacity (**Fig. S12f**) relative to bona fide A1-LCD seeds (**Fig. S7c**).

**Figure 6.**
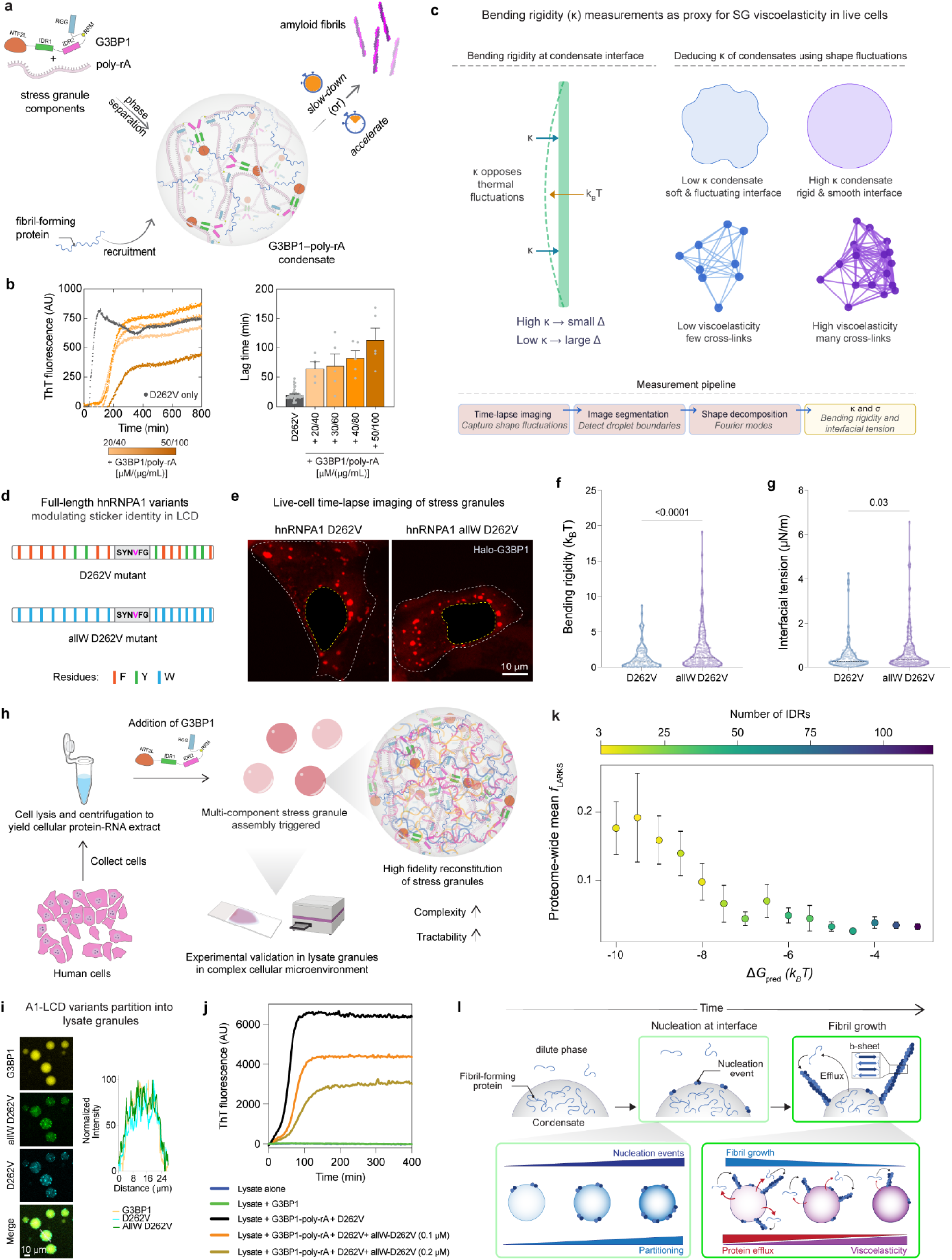
Reconstituted stress granules suppress amyloid fibril formation. **(a)** Schematic illustrating the experimental system to test whether condensates formed by primary SG components, G3BP1 and poly-rA, either promote or suppress amyloid fibril formation of A1-LCD. G3BP1 domains: NTF2L (NTF2-like dimerization domain), IDR1/IDR2 (intrinsically disordered regions), RRM (RNA recognition motif), RGG (arginine/glycine-rich domain). **(b)** (left) Kinetics of fibril formation monitored by ThT fluorescence of 25 μM A1-LCD D262V either alone or in the presence of G3BP1–poly-rA condensates at a range of volume fractions. Baseline subtraction was applied. (right) Lag times (*t*_5%_) extracted from ThT kinetics (shown on the *left*) and replicate experiments (n ≥ 3) are shown along with the mean ± SEM. **(c)** Schematic showing the conceptual framework of bending rigidity and its link to condensate viscoelasticity. The workflow for flicker spectroscopy is shown at the bottom. **(d)** Schematic showing full-length hnRNPA1 mutants with aromatic amino acid substitutions to the LCD. Only the aromatic residue patterning of LCD is captured in this diagram. **(e)** Fluorescence images of live U2OS cells treated with sodium arsenite, expressing HaloTag-G3BP1 along with either D262V or allW D262V variants of hnRNPA1. Fluorescence visualization is aided by JF646 HaloTag ligand. **(f)** Flicker spectroscopy-based SG bending rigidity measurements of D262V and allW D262V conditions are represented as a truncated violin plot with individual measurements indicated, from independent replicate experiments, along with the median (SG sample size: D262V, N=111; allW D262V, N=220). **(g)** Flicker spectroscopy-based SG interfacial tension measurements of D262V and allW D262V conditions are represented as a truncated violin plot with individual measurements indicated, from independent replicate experiments, along with the median (SG sample size: D262V, N=111; allW D262V, N=220). The *P* values indicated in (f) and (g) are estimated using the non-parametric Mann-Whitney test. **(h)** Schematic illustrating the high-fidelity stress granule reconstitution method from human cell lysate. **(i)** Fluorescence images showing colocalization of the indicated A1-LCD variants in stress granules reconstituted from cell lysate, along with the corresponding line profile. **(j)** ThT kinetics of various conditions, as indicated, using the reconstituted stress granule system from cell lysate. All ThT curves were baseline-subtracted. **(k)** Proteome-wide fraction of LARKS (*f*_LARKS_) as a function of the predicted phase separation propensity (*ΔG*_pred_). Data are binned in *ΔG*_pred_ intervals of 0.5 *k_B_T* and shown as mean ± SEM. **(l)** Schematic summarizing the main findings of this study, highlighting three key roles of condensates: sequestration of fibril-forming proteins, regulation of fibril nucleation through interface density, and control of protein efflux, and thus fibril growth, in the dilute phase via condensate viscoelasticity. Each experiment was repeated at least three times with consistent results.

To test whether these observations extend directly to even more complex granules, we utilized a combination of two experimental systems: (i) SGs induced by sodium arsenite stress in live human cells (**Fig. 6c-g**), and (ii) cell lysate–based, reconstituted SGs^50^, which recapitulate the complex compositional features of SGs in living systems while allowing for tractable experimental measurements that are not feasible in living cells (**Fig. 6h-j**). To examine how the expression of protein variants may alter the viscoelasticity in complex granules, we measured the bending rigidity (*κ*) and interfacial tension (σ) of SGs in cells by tracking condensate shape fluctuations using flicker spectroscopy^51,52^ (**Fig. 6c**). Bending rigidity provides a proxy for condensate viscoelasticity by reporting on the resistance of the condensate network to deformation. Therefore, bending rigidity measurements enable relative comparisons of mechanical stiffness across condensate systems for which direct rheological measurements are not feasible. Leveraging sticker variants in the background of the pathogenic D262V variant in full-length hnRNPA1, we tested whether a variant in which all phenylalanine and tyrosine residues outside the steric zipper were replaced with tryptophan residues (allW D262V; **Fig. 6d**) can increase SG viscoelasticity. We find that both constructs support sodium arsenite–induced SG assembly, consistent with our earlier work, where the D262V mutant was observed to delay SG disassembly and the allW D262V variant rescued this phenotype^21^ (**Fig. 6e**). Notably, flicker spectroscopy measurements of SGs in live cells either in the presence of D262V or allW D262V show that expression of the tryptophan variant leads to increase in both bending rigidity (*κ*) and interfacial tension (σ) of SGs (**Fig. 6f, g**). This is consistent with the expectation that increasing the strength of the intra-condensate network results in higher viscoelasticity^25^. To obtain a readout for changes in protein fibril assembly kinetics in response to increased SG viscoelasticity, we next reconstituted SGs by adding G3BP1 to cell lysates, which triggers the formation of lysate-based SGs^50^ (**Fig. 6h**). We confirmed that the tryptophan-containing variant of A1-LCD D262V (allW-D262V) and D262V partition into the lysate-based SGs (**Fig. 6i**). ThT kinetics reveal that the addition of allW-D262V to samples containing lysate-based SGs dampens fibril assembly of D262V in a dose-dependent manner (**Fig. 6j, Fig. S12h**). However, under conditions without condensates, fibril assembly progresses unperturbed (**Fig. S12g**). Together, these results indicate that increasing the viscoelasticity of complex stress granules slows fibril assembly.

If SGs were to serve as protective sinks that suppress amyloid formation, an expectation would be that various fibril-forming proteins across the human proteome, such as those containing steric zippers or low-complexity aromatic-rich kinked segments (LARKS), would favor recruitment into the condensate microenvironment and/or undergo phase separation to avoid aggregation under cellular stress. Conversely, condensate-forming proteins could better tolerate the accumulation of such aggregation-prone motifs. To test these ideas, we identified disordered regions containing LARKS across the entire human proteome and estimated the free energy for phase separation of each protein [*ΔG* (*k_B_T*)] using a machine-learning model trained on simulation data from the CALVADOS 2 model^53,54^. The resulting analyses show that the proteins that harbor phase separation propensity most strongly are enriched in LARKS (**Fig. 6k; Fig. S13**), suggesting that condensates may suppress negative consequences of LARKS-mediated protein aggregation across the human proteome. An alternative explanation is that proteins that spend part of their lifetime within cellular condensates experience reduced selective pressure against accumulating steric zippers and LARKS.

Collectively, these findings suggest that SGs and similar condensates evolved to suppress fibril formation of aggregation-prone proteins and that their physical properties, such as sequestration of soluble protein in the interior of condensates, interface-mediated nucleation, and efflux-driven fibril growth in the dilute phase, govern amyloid assembly kinetics in the presence of multi-component condensates (**Fig. 6l**).

### Multi-component biomolecular condensates as programmable biomaterials that suppress amyloid fibril assembly

Our quantitative and systematic measurements and the use of a heterotypic peptide–nucleic acid condensate platform with physicochemical tunability provide direct mechanistic insights into the influence of multi-component SG mimics on the fibril formation of the disease-relevant protein A1-LCD (**Fig. 6l**): (1) Fibrils grow in the dilute phase. Therefore, sequestration of fibril-forming protein in condensates suppresses fibril assembly. Multi-component condensates can lower the dilute phase concentration of fibril-forming proteins below *c*_sf_, thereby effectively eliminating fibril growth. (2) At low dilute phase concentrations of the fibril-forming protein, condensate interfaces are sites of fibril nucleation. Dilution of fibril-forming protein at interfaces in multi-component condensates lengthens lag times and therefore strongly delays fibril formation. (3) The protein efflux rate from condensates governs fibril growth in the dilute phase. (4) Highly viscoelastic condensates are characterized by slow protein efflux, which contributes to suppression of fibril growth in the dilute phase. Importantly, these detailed insights gained from mechanistic interrogation of the tunable repeat peptide–nucleic acid condensates are generalizable to the effect of condensates on several fibril-forming proteins that play relevant roles in the pathogenesis of neurodegenerative diseases, including FUS and Tau. SG mimics formed by G3BP1 and poly-rA, as well as complex SGs reconstituted in mammalian cell lysate, also suppress fibril formation.

Our results show decisive benefits of multi-component over mono-component condensates towards suppressing fibril formation, highlighting their advantageous emergent material properties. Mono-component condensates are metastable assemblies compared to fibrils and can serve as kinetic sinks for soluble proteins^21^. However, in mono-component condensates, enhanced metastability and the resultant high sink potential are inherently coupled to a high density of the fibril-forming protein at interfaces, which in turn promotes nucleation. Consequently, these condensates exhibit a duality in which the condensate interior suppresses fibril formation while the interface facilitates it. Multi-component condensates also sequester fibril-forming proteins and lower their dilute phase concentration below *c*_sf_, preventing fibril growth, but they offer additional degrees of freedom. Specifically, multi-component condensates enable decoupling of sink potential from interfacial protein density. As a result, condensates with highly viscoelastic network properties but relatively low densities of fibril-forming protein at interfaces are most effective at suppressing fibril formation. Importantly, the ability of multi-component condensates to suppress amyloid assembly cannot be reproduced by the behavior of individual peptide and DNA components, underscoring the emergent properties of condensates (**Supplementary Note 5; Fig. S2**).

Our results suggest that RNP granules optimally suppress pathological fibril assembly if they (a) lower the dilute phase concentration of fibril-forming protein below *c*_sf_, (b) dilute the relative density of any single fibril-forming protein at interfaces by recruiting multiple proteins through favorable homotypic and heterotypic interactions, and (c) engage in strong networking interactions with the fibril-forming protein within the condensate microenvironment to control the rate of protein efflux. We posit that the multi-component nature of SGs achieves these properties and promotes heterotypic buffering^55,56^, through thermodynamic frustration^57,58^. In support of point (c), our experiments utilizing lysate granules aimed at enhancing the strength of the PLD network by mutagenesis successfully delayed the onset of fibril formation. Given that fibril nucleation can be facilitated at condensate interfaces, the rheology of the condensate-dilute phase interface is likely to play a critical role in modulating nucleation processes. This is an important avenue for future investigation.

The interiors of mono-component condensates can suppress fibril formation, presumably because condensate and fibril formation often involve overlapping sequence determinants in fibril-forming proteins, creating competing interactions that can hinder fibril assembly^21,33^. However, we expect that not all condensates will suppress fibril formation of all fibril-forming proteins, because the extent of suppression will depend on the specific overlap between the sequence features that drive condensate formation and those that promote fibril assembly. Therefore, some condensates may be permissive, or even facilitative, of fibril formation if the fibril-forming sequence features are not engaged in the condensate network^59^. This nuanced view is supported by the observation that condensates formed by the intrinsically disordered region of CAPRIN1 suppress fibril formation of the RRM domain of FUS but apparently promote fibril formation of SOD1, although the CAPRIN1 condensates destabilize the folded state of both proteins^60,61^. Nevertheless, our work establishes the suppressive function of a wide array of condensates towards fibril formation of several classical proteins involved in neurodegenerative disorders. We also speculate that the highly viscoelastic interior of condensates can impose a mechanical barrier to fibril nucleation and growth.

In summary, our findings suggest that SGs and other RNP condensates do not typically promote protein fibril formation^62–64^, and therefore do not serve as crucibles for numerous neurodegenerative disorders^12^. SGs may instead buffer against stress-induced protein aggregation. Mutations that reduce the strength of incorporation into the condensate network or stabilize fibrils may overcome this buffering function and nevertheless drive disease processes. Future research will further test the hypothesis that the stabilization of SG interiors can rescue key pathogenic processes driven by RBP fibril formation.

## Acknowledgments

We thank Dr. Rohit Pappu (Washington University in St. Louis) for valuable discussions at various stages of the manuscript preparation, and Dr. Sören von Bülow for discussions regarding predictions of phase separation. The authors acknowledge the use of AI tools to enhance the readability and clarity of certain sections of the manuscript text. The authors reviewed and edited all content generated with the assistance of AI tools and take full responsibility for the content of this manuscript.

## Funding

National Institutes of Health grant R01NS121114 (TM)

National Institutes of Health grant R35GM138186 (PRB)

National Institutes of Health grant R35GM138243 (DAP)

St. Jude Children’s Research Collaborative on the Biology and Biophysics of RNP Granules (PRB, TM)

American Lebanese Syrian Associated Charities (TM)

Novo Nordisk Foundation PRISM (Protein Interactions and Stability in Medicine and Genomics centre NNF18OC0033950 (KL-L)

Any opinions, findings, conclusions, or recommendations expressed in this material are those of the author(s) and do not necessarily reflect the views of the funding agency or any other bodies.

## Author contributions

Conceptualization: TM, PRB

Methodology: TSM, XG, AB, AS, FKZ, GT, JLB, KL-L, DAP, TM, PRB

Investigation: all authors

Visualization: TSM, XG, AB, AS, FKZ, GT, JLB, DAP, TM, PRB

Project administration: TM, PRB

Supervision: KL-L, TM, PRB

Resources: KL-L, DAP, TM, PRB

Writing – original draft and revision: TSM, TM, PRB

Writing – reviewing and editing - all authors

Funding acquisition: KL-L, DAP, TM, PRB

## Data availability

All data are available in the manuscript or the supplementary materials.

## Code availability

The scripts used for Thioflavin T kinetics and rheology data analyses in the current study are available on GitHub (see github.com/BanerjeeLab-repertoire/Interface-density-and-viscoelasticity-of-heterotypic-condensates-determine-their-ability-to-suppress). Data and code to reproduce the proteome-wide analyses of the association between fraction of LARKS and phase separation propensity are available at github.com/KULL-Centre/_2026_Mahendran_suppression_amyloid.

## Supplementary Materials

Materials and Methods

Figs. S1 to S13

Supplementary Notes

Tables S1 to S5

Captions for Videos S1 to S6

Videos S1 to S6

