## Supplementary Information for "Viscoelasticity and Interface Properties of Multi-Component Condensates Govern Protein Sequestration and Suppression of Amyloid Formation"

**The PDF file includes:**

Materials and Methods
Figs. S1 to S13
Supplementary Notes
Tables S1 to S5
Captions for Videos S1 to S6
References

**Other Supplementary Materials for this manuscript include the following:**

Videos S1 to S6

### Materials and Methods

#### Details of protein constructs and protein purification

*Expression and purification of the low-complexity domain (LCD) of human hnRNPA1 (UniProt: P09651; Isoform A1-A)*: The construct spans residues 186-320 (A1-LCD) with a 6xHis-tag and an N-terminal TEV protease cleavage site (ENLYFQGS) in a pDEST17 (ThermoFisher) expression vector as previously described [1,2]. Protein expression and purification from inclusion bodies were performed as previously described [1]. Briefly, A1-LCD was purified under denaturing conditions, and the final pure protein was stored in 6 M guanidinium hydrochloride (GdmHCl), 20 mM MES, pH 5.5, at 4 $^{o}$C. The purity and identity of the proteins were confirmed using SDS-PAGE and intact mass spectrometry. For sequence details of all A1-LCD variants used in the study, refer to **Table S1**.

*Expression and purification of MBP-His-tagged full-length G3BP1*: The construct of full-length G3BP1 used in this study was synthesized and cloned into a pET His6 MBP N10 TEV LIC cloning vector (Addgene plasmid #29706) by GenScript USA Inc. (NJ, USA). The MBP-His-G3BP1 construct was transformed into BL21-RIPL cells, which were grown in LB media at 37 $^{o}$C to an $OD_{600}$ of ~0.8, then induced with 0.6 mM IPTG and incubated overnight at 16 $^{o}$C. Cells were harvested by centrifugation at 4520 g for 12 min and lysed with a microfluidizer in lysis buffer containing 50 mM HEPES (pH 7.5), 500 mM NaCl, 1 mM EDTA, protease inhibitor cocktail (Roche), 5 mM β-mercaptoethanol, and 10 mM imidazole. After centrifugation at 27,812 g for 30 min at 4 $^{o}$C, the clear supernatant was loaded onto a HisTrap HP 5 mL column (Cytiva) and eluted with a step gradient of imidazole. All fractions containing G3BP1 were collected, and 1:100 TEV protease (w/w) was added at room temperature and incubated for 2 h while dialyzing into TEV cleavage buffer containing 50 mM HEPES (pH 7.5), 200 mM NaCl, 2 mM DTT, and 1 mM EDTA. The protein was diluted 5- to 10-fold to reduce the salt concentration, loaded onto a 5 mL Heparin column (Cytiva), and then eluted with a NaCl step gradient in 50 mM HEPES (pH 7.5) and 2 mM DTT. Pure protein fractions eluted in buffer containing 50 mM HEPES (pH 7.5), 450 mM NaCl, 2 mM DTT were pooled, concentrated, aliquoted, and stored at -80 °C for future use.

*Expression and purification of His-SUMO-tagged full-length hnRNPA1*: Protein expression was performed in BL21(DE3)-RIPL cells (Agilent) cultured in LB medium and induced with 0.5 mM IPTG. Following induction, cultures were incubated overnight at 25°C with agitation, and the cells were harvested by centrifugation (4520 g, 20–25 min, 4°C). Cell pellets were collected and stored at -80 °C until further use. For lysis, pellets were resuspended in buffer containing 50 mM HEPES (pH 7.5), 1 M NaCl, 30 mM imidazole, 2 mM β-mercaptoethanol, complete protease inhibitor (Roche), and 0.1 mM PMSF. Cells were disrupted by passing them twice through a microfluidizer, lysate was centrifuged at 27,812 g for 40 min at 4 °C to remove cell debris, and the resulting supernatant was loaded onto a gravity-flow Ni-NTA affinity column after equilibrating the column with buffer A (50 mM HEPES pH 7.5, 500 mM NaCl, 30 mM imidazole, 2 mM β-mercaptoethanol). The resin was washed twice with buffer A, followed by two washes with buffer B (50 mM HEPES pH 7.5, 1 M NaCl, 30 mM imidazole, 2 mM β-mercaptoethanol), and a final wash with buffer A. Bound protein was subsequently eluted in 50 mM HEPES (pH 7.5), 150 mM NaCl, 300 mM imidazole, and 2 mM β-mercaptoethanol. Removal of the His-SUMO tag was achieved by overnight incubation of the protein with Ulp1 protease at 4°C. The sample was then diluted to lower the imidazole concentration to 30 mM and reapplied to a gravity-flow Ni-NTA column. After washing with buffer containing 50 mM HEPES pH 7.5, 150 mM NaCl, 30 mM imidazole, and 2 mM β-mercaptoethanol, the flow-through and wash fractions containing the cleaved protein were collected, pooled, and diluted to a final NaCl concentration of 40 mM. Further purification was performed using anion exchange (HiTrap Q) and cation exchange (HiTrap SP HP) chromatography columns (GE Healthcare) connected in tandem, enabling RNA to bind to the anion exchange column while the protein was bound to and was eluted from the cation exchange column. Elution was achieved using a linear NaCl gradient generated from buffer A (50 mM HEPES pH 7.5, 50 mM NaCl, 5 mM DTT) and buffer B (50 mM HEPES pH 7.5, 500 mM NaCl, 5 mM DTT). Fractions were assessed by SDS-PAGE, pooled, and concentrated. Final purification was carried out by size exclusion chromatography on a Superdex 200 16/60 column (GE Healthcare) equilibrated in 50 mM HEPES (pH 7.5), 300 mM NaCl, and 2 mM TCEP. Peak fractions were analyzed by SDS-PAGE to determine purity, combined, concentrated, flash-frozen in liquid nitrogen, and stored at -80 °C.

*Expression and purification of eFUS-LCD (residues 1–273):* The eFUS-LCD construct (residues 1–273) was cloned into a pET His6 MBP N10 TEV LIC vector (Addgene plasmid #29706) and transformed into BL21(DE3) pLysS cells. Cells were grown in LB medium at 37 °C to an $OD_{600}$ of ~0.8, then induced with 0.5 mM IPTG, and incubated at 30 °C for 6 h. Cells were harvested by centrifugation, and the pellets were stored at −80 °C until protein purification. The protein was expressed with an N-terminal 6×His-tag in E. coli and purified under native conditions by sequential Ni-NTA affinity chromatography and size-exclusion chromatography (SEC). Bacterial cell pellets were resuspended in lysis buffer (300 mM NaCl, 50 mM Tris, 10 mM imidazole, pH 7.5) supplemented with protease inhibitor cocktail (Thermo Scientific), at approximately 4–5 mL per gram of wet pellet, and lysed by probe sonication (55% amplitude, 10 s on/50 s off, 2 min total on-time) on ice. The lysate was clarified by centrifugation at 37,500 × g for 20 min at 4 °C, and the supernatant was incubated with pre-equilibrated Ni-NTA resin (1 mL resin per 500 mL culture) for 1 h at 4 °C with end-over-end rotation. The resin was loaded onto a gravity-flow column and washed sequentially with 2 column volumes (CVs) of wash buffer 1 (1 M NaCl, 50 mM Tris, 20 mM imidazole, pH 7.5) and 1 CV of wash buffer 2 (125 mM NaCl, 25 mM Tris, 20 mM imidazole, pH 7.5). Protein was eluted in 1 mL fractions with elution buffer (125 mM NaCl, 25 mM Tris, 250 mM imidazole, pH 7.5), and fractions with significant $A_{280}$ absorbance were pooled and concentrated to ~1 mL using a 4 mL Amicon centrifugal filter unit (MilliporeSigma, UFC803024; 30 kDa filter; 3,000 × g, 4 °C). The concentrated sample was filter-sterilized through a 0.22 µm membrane prior to injection. SEC was performed on a Superdex 200 Increase 10/300 GL column (Cytiva, 28990944) equilibrated in storage buffer (125 mM NaCl, 25 mM Tris, pH 7.5) at 0.5 mL/min on an ÄKTA purifier system; 0.5 mL fractions were collected across the elution volume, and those corresponding to the primary UV absorbance peak were retained. Protein purity was confirmed by SDS-PAGE. For sequence details, refer to **Table S1**.

*Expression and purification of SynTag-Tau*: For details of the expression and purification of SynTag-Tau, refer to the methods described in a recent work[3]. For sequence details, refer to **Table S1**.

### Buffer exchange of A1-LCD proteins to remove denaturant

Buffer exchange of A1-LCD variants was achieved using a 5 mL desalting column (Cytiva) attached to a peristaltic pump P-1 (Pharmacia Fine Chemicals). The column was equilibrated with 5 column volumes of 25 mM MOPS (pH 7.5), and 5 mM TCEP. Approximately 0.6-1 mL of protein solution stored in 20 mM MES (pH 5.5), 6 M GdmHCl was injected onto the column with a 1 mL syringe. The flow rate of the peristaltic pump was adjusted to 1 mL/min, and 0.5 mL fractions per aliquot were collected. Fractions containing the protein solution were pooled and concentrated using 0.5 mL 3K molecular weight cut-off (MWCO) Amicon centrifugal filters (Merck, UFC500324). For experiments using confocal fluorescence imaging and microrheology, multiple rounds of buffer exchange steps were conducted using 0.5 mL 3K MWCO Amicon centrifugal filters (Merck).

Subsequently, the protein solution was filtered using a 0.5 mL 100K MWCO Amicon centrifugal filter (Merck, UFC510024) to remove any potential aggregates.

#### Fluorophore labeling of A1-LCD variants and other fibril-forming proteins

A1-LCD protein variants were N-terminally labeled with either NHS ester Alexa Fluor 488 (ThermoFisher Catalog# A20000) or NHS-rhodamine (ThermoFisher Catalog# 46406). The labeling reaction was performed in 20 mM HEPES (pH 7.4; HEPES), 2 M GdmHCl denaturant buffer for 1 h in the dark, followed by dialysis at room temperature overnight. The labeled protein solutions were stored in 20 mM MES (pH 5.5), 2 M GdmHCl at 4 °C until usage.

eFUS-LCD and SynTag-Tau were labeled with Alexa488 (Invitrogen Catalog# A10254) via cysteine–maleimide chemistry following the manufacturer-provided instructions. The native cysteine residues of the wild-type Tau sequence were used for labeling, whereas with eFUS-LCD, a cysteine-containing variant (single cysteine residue appended to the N-terminus) of the protein was purified following a similar purification workflow as the non-cysteine variant, outlined in the '*Details of protein constructs and protein purification*' method section, and was used for labeling. Storage buffer conditions, as used in the purification of either protein (except for DTT, which was excluded), were used for the labeling reactions. Subsequently, buffer exchange was conducted using Zeba Spin Desalting Columns (ThermoFisher Scientific, Catalog #89890) to remove free dye (same buffer conditions as before, with 2 mM DTT included).

#### Preparation of polypeptide and single-stranded nucleic acid stocks

All polypeptides used in this study were synthesized by GenScript USA Inc. (NJ, USA). For site-specific fluorophore labeling using cysteine–maleimide chemistry[4], a cysteine residue was incorporated at the C-terminus of the polypeptide. Labeling of the polypeptide with Alexa594 (Invitrogen Catalog# A10256) was performed via cysteine–maleimide chemistry following the manufacturer-provided instructions. Each polypeptide was reconstituted in RNase-free water (Santa Cruz Biotechnology), including 50 mM DTT to prevent cysteine oxidation from occurring during storage. Excess free dye was removed by acetone precipitation as follows. Ice-cold acetone (−20 °C) was added to the labeling reaction at a 4:1 (v/v) ratio, vortexed, and incubated at −20 °C for 60 min. The precipitate was collected by centrifugation at 15,000 × g for 10 min, and the supernatant was carefully decanted. This wash cycle was repeated four times in total. The pellet was air-dried at room temperature for 30 min and resuspended in the required buffer condition. Single-stranded DNA ($dT_{40}$) and RNA ($rU_{40}$), including Cy5-$dT_{40}$ and FAM-$rC_5$, were purchased from Integrated DNA Technologies (IDT). Single-stranded poly-rA was purchased from Sigma (Catalog# 10108626001). Each nucleic acid was reconstituted in RNase-free water. The reconstituted solutions were centrifuged at high speed (23,000 g) for 2 min to remove any potential solid particles or aggregates, and the supernatant was subsequently used. The concentration of nucleic acid was determined using a NanoDrop 1C spectrophotometer. The final stock solutions of polypeptides and nucleic acids were divided into multiple aliquots and stored at −20 °C. For sequence details of all polypeptides and nucleic acids used in the study, refer to **Tables S2 and S3**.

#### Preparation of multi-component biomolecular condensates for fluorescence imaging

Peptide-nucleic acid condensates containing A1-LCD were prepared by mixing the peptide, nucleic acid, and A1-LCD in a buffer containing 25 mM MOPS (pH 7.5), 150 mM NaCl, and 5 mM DTT. As the final component, the nucleic acid was introduced into the sample mixture to facilitate condensate formation. For fluorescence microscopy visualization of multi-component condensates, trace amounts (250 nM) of either Alexa594-labeled polypeptide, Alexa488-labeled A1-LCD, Cy5-labeled $dT_{40}$, or combinations thereof were used. Amyloid-sensitive fluorescent dyes, such as CRANAD2 (Tocris, Catalog# 4803) or thioflavin T (ThT; Thermo Scientific Chemicals, J61043.14), were used at concentrations of 10 μM. For all experiments conducted in this study, the mass ratio of polypeptide to nucleic acid is 1:1, which was shown previously to be favorable for peptide–nucleic acid condensate formation [5]. For fluorescence imaging experiments, 1 mg/mL polypeptide, 1 mg/mL nucleic acid, and 25 μM A1-LCD variant were used. In the case of eFUS-LCD or SynTag-Tau containing ternary systems, the protein concentration used is 25 μM and 12 μM, respectively, and the polypeptide/nucleic acid concentration was kept the same. In the case of eFUS-LCD containing samples, TEV protease was added to MBP-tagged eFUS-LCD at a final concentration of 0.02 mg/mL prior to the addition of ssDNA/peptide, enabling TEV mediated cleavage of MBP tag. Once the final sample was prepared, consisting of a volume of 5 μL, it was sandwiched between a Tween 20-coated coverslip and a glass slide using double-sided tape. Immediately after, mineral oil was used to fill the void space in the sample chamber to prevent sample evaporation during the experimental timeframe.

The G3BP1−poly-rA condensates containing G3BP1, poly-rA, and A1-LCD were reconstituted in a buffer containing 50 mM Tris-HCl (pH 7.5) and 150 mM NaCl. The poly-rA was introduced to the mixture as the final component to induce condensate formation. For the imaging experiments, the concentration of G3BP1 was kept at 80 μM, the poly-rA concentration was kept at 80 ng/μL, and the concentration of A1-LCD was kept at 25 μM. For fluorescence imaging, either Alexa488-labeled A1-LCD, CRANAD2, Cy5-labeled $dT_{40}$ or combinations thereof were used at concentrations similar to those used in experiments involving peptide–nucleic acid–A1-LCD condensate systems.

#### Confocal fluorescence imaging and post-processing

Fluorescence imaging of condensate samples was carried out either using a Q2 laser scanning confocal microscope (ISS Inc., 63× objective, 1.33 NA) or an Andor Dragonfly 600 confocal platform (Oxford Instruments) installed on a Leica DMi8 inverted microscope equipped with an Andor Zyla 4.2 plus sCMOS camera and a Leica HC PL APO 63x/1.40-0.60 oil objective. For correlative optical tweezer and fluorescence microscopy measurements, a Lumicks C-trap featuring a dual-trap optical tweezer and confocal fluorescence microscope system, equipped with a 60× water immersion objective (1.33 NA), was used. The acquired fluorescence images were visualized using Fiji [6] (v1.54f). For illustrative purposes, brightness and contrast were adjusted individually for each image based on its histogram to visualize relevant features such as the coexistence of droplets and fibrils. These adjustments were applied uniformly within each image. This information is also noted in the appropriate figure legends. These adjustments were made solely for presentation purposes and not for downstream quantification. In all cases, quantitative analysis was performed on unmodified raw data. Additionally, for images acquired using the Andor Dragonfly 600 confocal microscope, the gain was adjusted using Imaris (v10.1, Bitplane) in order to enhance the visibility of lower-intensity ThT-positive fibrillar structures in the dilute phase. These gain-adjusted images appeared as the following panels: **Fig. 1i-k**. For generating a 3D render from a Z-stack acquired using the Andor Dragonfly 600 confocal microscope, the 'blend' mode of Imaris, a volume rendering technique, was used. In the case of Video S3, the raw time-lapse video was upscaled via bilinear interpolation using Fiji[6] to enhance clarity for better visualization. Additional details, if any, on image processing are mentioned within the respective method section.

**Thioflavin T (ThT) kinetics assay and seed generation**

Fibril formation kinetics were monitored using a Thioflavin T (ThT; Calbiochem, Cat# 596200) fluorescence assay performed on a CLARIOstar (BMG Labtech) plate reader using black 384-well plates (Greiner Bio-One Catalog# 784900). The plate was sealed with optically transparent film (HD Clear tape from ShurTech Brands) to prevent evaporation, and the plate was incubated with shaking at 500 rpm between readings. Each well contained 30 µL and a final ThT concentration of 20 µM. Fluorescence excitation was performed at 450 nm, and emission spectra were recorded at 510 nm. Fibril formation was monitored until the ThT intensity reached a plateau value wherever possible. Data plotting was performed in GraphPad Prism 10 (Version 10.2.3). All measurements with either nucleic acid–peptide condensate systems were carried out in 25 mM MOPS (pH 7.5), 150 mM NaCl, 5 mM TCEP unless specified otherwise. Distinct preparations for different systems were carried out as follows:

For ThT assays with SynTag-Tau, SynTag-Tau was buffer-exchanged into 20 mM MOPS (pH 7.5), 200 mM NaCl, and 5 mM TCEP using 2 mL Zeba Spin Desalting Columns (ThermoFisher Scientific, Catalog #89890). PEG 8000 was mixed with buffer (20 mM MOPS, pH 7.5, 5 mM TCEP) to a final concentration of 5% prior to the addition of ssDNA, peptide, and SynTag-Tau. The final NaCl concentration was 80 mM.

For ThT kinetics experiments with eFUS-LCD (1–273), TEV protease was added to MBP-tagged eFUS-LCD at a final concentration of 0.02 mg/mL prior to the addition of ssDNA/peptide, enabling initial TEV cleavage followed by fibril formation. The final salt concentration was adjusted to 150 mM NaCl.

For ThT kinetics with A1-LCD, we performed a 2x dilution series for the different nucleic acid/peptide condensates, while keeping the final A1-LCD protein concentration at 25 µM.

For ThT kinetics experiments involving comparison of dilute phase–only samples with bulk samples, A1-LCD and peptide-ssDNA samples were thoroughly mixed with 20 µM ThT and equally divided into two Eppendorf tubes. One tube was incubated at 26 °C for 10 min before transfer to a 384-well plate. The other tube was centrifuged at 17,000 × g for 10 min, and the resulting supernatant was transferred to the same plate.

For the ThT assays with A1-LCD D262V and additions of A1-LCD ∆hexa, A1-LCD D262V was premixed with A1-LCD ∆hexa before adding ssDNA and peptide.

For the ThT assays with G3BP1–poly-rA condensates, G3BP1-poly-rA were pre-mixed before adding A1-LCD and ThT. The assays were carried out in buffer containing 50 mM HEPES (pH 7.5), 150 mM NaCl, 2 mM DTT, and a final ThT concentration of 50 µM.

Analysis of ThT fluorescence data was performed using a custom script written in R. The data were baseline subtracted, and some curves were smoothed using a Gaussian kernel smoother function with a bandwidth of 15 minutes if indicated in the figure captions. The locations of baseline, growth phase, and plateau for each curve were manually defined. The lag times ($t_{5\%}$) were calculated as the time to reach 5% of the plateau value.

To generate seeds, we incubated A1-LCD protein either alone or in the presence of $(RGRGG)_5$–$dT_{40}$ condensates/G3BP-poly-rA condensates at 20 °C on a bench-top incubator shaker with shaking at 600 rpm for 2 days until fibril formation was complete, as indicated by ThT curves (see **Fig. 1**, **Fig. S1**, and **Fig. S3**). Fibrils were collected using ultracentrifugation in a Beckman Coulter Ultracentrifuge (Optima) at 218,000 g at 20 °C for 40 min. After removing the majority of the supernatant, the fibrils were resuspended in the remaining solution. The fibrils were subsequently sonicated for 60 min in a water bath. Varying concentrations of seeds were added to freshly buffer-exchanged A1-LCD protein, and ThT fluorescence was monitored.

#### Differential interference contrast (DIC) microscopy

For DIC microscopy images, 2.5 µL of the sample, which was prepared following the protocol outlined in the '*Thioflavin T kinetics assay and seed generation*' method section, was sandwiched between a coverslip and a glass slide using 3M 300 LSE high-temperature double-sided tape (0.34 mm). DIC images were obtained at room temperature using a Nikon Eclipse Ni Widefield microscope with a 20x 0.75 NA DIC N2 objective.

#### Negative-stain transmission electron microscopy (nsTEM)

Protein, peptide, and ssDNA were mixed together in a 1.5 mL ultracentrifuge tube (Beckman Coulter). The total volume was 100 µl with a final protein concentration of 100 µM and final peptide and DNA concentrations of 4 mg/mL in 25 mM MOPS (pH 7.5), 150 mM NaCl, and 5 mM TCEP. The samples were incubated at 20 °C and with 600 rpm shaking on a bench-top incubator shaker for 6 days until fibril formation was complete. To pellet fibrils that were formed over time, the samples were centrifuged using a Beckman Coulter Ultracentrifuge (Optima) at 218,000 g at 20 °C for 40 min. The supernatant was removed, and 4 µL volume of fibrils and/or higher oligomeric assemblies was transferred onto 300-mesh copper grids (Catalog# CF300-Cu-50, Electron Microscopy Sciences), freshly plasma cleaned with an Ar/$O_2$ gas mixture for 10 sec using Solarus plasma cleaner (Gatan). The samples were allowed to adsorb for 30 sec before blotting away the excess liquid, followed by staining using three successive applications of 2% uranyl acetate (Catalog# 22400-2, Electron Microscopy Sciences). The last round of stain application was allowed to sit for 45 sec before blotting away the excess stain. The grids were air-dried prior to imaging using a 120 kV Talos L120C TEM (ThermoFisher Scientific) equipped with a CETA detector (TFS).

#### Determination of protein dilute phase concentration ($c_{\text{dilute}}$) via analytical HPLC and saturation concentration for fibril formation ($c_{\text{sf}}$)

The dilute phase concentration of the protein ($c_{\text{dilute}}$) in the presence of peptide–ssDNA condensates was measured via analytical HPLC, as the overlapping absorbance spectra of protein, peptide, and DNA necessitated separation of the components [7]. The measurements were carried out in 25 mM MOPS (pH 7.5), 150 mM NaCl, 5 mM TCEP, and the input peptide–ssDNA concentrations are indicated in the respective figures. In these experiments, the bulk concentration of WT A1-LCD was 25 µM. All components were mixed and incubated for 10 min at 20 °C. Following incubation, the dense and dilute phases were separated by centrifugation at 20,000 g for 5 min at 20 °C. The supernatant was carefully removed without disturbing the dense phase. For determination of $c_{\text{dilute}}$, a known amount of the supernatant was removed and mixed 1:1 with 20 mM MES (pH 5.5), 6 M GdmHCl to prevent the onset of fibril formation until measurements are completed. The samples were run over a C4 (Reprosil Gold 200, Dr. Maisch) reverse-phase column attached to a Waters HPLC system with a dual-channel UV/Visible Detector (Waters 2489). Buffer A consisted of dd$H_2O$ with 0.1% TFA, while buffer B was 100% acetonitrile. The proteins were eluted from the column by running a linear gradient from 5% to 80% buffer B.

In order to determine the $c_{\text{dilute}}$ concentration from the HPLC elution profile, a standard curve was generated in parallel by injecting 5 different volumes of the protein with known concentrations.

For further details on the analysis, please refer to previous work [7]. At least three replicate experiments per sample were performed, and the error bars represent the standard error.

For measurement of the saturation concentration for fibril formation ($c_{sf}$), 20 μM WT A1-LCD was prepared in triplicate in buffer containing 25 mM MOPS (pH 7.5), 150 mM NaCl, and incubated in a thermo-mixer (Eppendorf, ThermoMixer C) at 20 °C with 600 rpm shaking for 14 days. The samples were then centrifuged at 20,000 g for 1 h on a bench-top centrifuge. The supernatants were carefully removed without perturbing the pellet. The concentrations of the soluble protein in the supernatants were measured using a micro BCA kit (Thermo Scientific Catalog# 23235).

For the measurement of the saturation concentration for fibril formation in the presence of 1 mg/mL $(RGRGG)_5$-$dT_{40}$, the supernatants were injected onto a HPLC C4 reverse-phase column. The concentration of A1-LCD was analyzed as previously described by comparison to a standard curve[7].

**Passive microrheology with optical tweezers (pMOT)**

Multi-component condensate systems of peptide–$dT_{40}$–A1-LCD consisting of 5 mg/mL peptide [either $(KGKGG)_5$, $(RGPGG)_5$, $(RPRPP)_5$, $(RGRGG)_5$, or $(RGYGG)_5$], 5 mg/mL $dT_{40}$, and 25 μM WT A1-LCD were prepared in a buffer containing 25 mM MOPS (pH 7.5), 150 mM NaCl, and 5 mM DTT. In the case of quaternary condensate systems including A1-LCD ∆hexa, the concentration of the protein included in the sample mixture is noted in the relevant figure legend or corresponding text. In addition, yellow-green carboxylate-modified polystyrene beads (1 μm in diameter, FluoSpheres, Invitrogen, F8823) were introduced into the sample buffer, at a low concentration (0.0005% solids), prior to phase separation, to allow beads to passively partition into condensates once phase separation is initiated. The condensates were prepared in a 10 μL sample volume and were placed on an 18 x 18 mm-sized coverslip, which was coated with Tween20 [20% (v/v)]. The sample was subsequently sandwiched with a 75 x 25 x 1 mm-sized glass slide using five layers of double-sided tape (Scotch 3M). To prevent evaporation, ~175 μL of mineral oil was injected into the chamber to surround the sample. The sample was then loaded onto a correlative optical-tweezer and confocal microscopy system (Lumicks, C-Trap) equipped with a 1.33 NA, 60x water immersion objective. The samples were allowed to equilibrate for ~15 min for the condensates to settle on the coverslip surface and until no more fusion events were observed. Condensates with a single 1 μm-sized polystyrene bead embedded within them were used for pMOT experiments. The optical trap at minimal power (~10-50 μW) was used to trap a bead inside a condensate, and a brightfield camera at 500 Hz was used to track the passive motion of the optically-trapped bead using a template-matching algorithm for 10–30 min. pMOT measurements were performed over 9 to 15 condensates over three independently prepared samples for each multi-component condensate system.

Below, the workflow of the data analysis for the pMOT measurements is presented as previously described [8]. The passive tracking of a bead inside a condensate with an optical trap outputs two-dimensional trajectories of trapped particles. These trajectories were analyzed to calculate the complex modulus [9,10] of the condensate as a function of frequency $\omega$

$$G^*(\omega) = G'(\omega) + i\, G''(\omega) \qquad (1)$$

where $G'$ and $G''$ are the frequency-dependent elastic and viscous moduli, respectively. The possible long-time drifts in the trajectories in X and Y were first removed using a spline-based detrending algorithm. We used the equipartition theorem [11] to obtain the trap stiffness of the optical trap in X and Y using the relation

$$\kappa_x = k_B T/\langle x^2 \rangle \tag{2}$$

where $\kappa_x$ is the optical trap stiffness in the x direction, $T$ is the temperature in Kelvin, and $k_B$ is the Boltzmann constant. Next, a normalized autocorrelation function $g(\tau)$ is calculated. The autocorrelation function value at $\tau = 0$ is extrapolated using a spline algorithm. The autocorrelation function is then transformed into the frequency domain $\hat{g}(\omega)$ using a discrete Fourier transform algorithm

$$-\omega^2 \hat{g}(\omega) = i\omega g(0) + \frac{\left(1 - e^{-i\omega t_1}\right)\left(g_1 - g(0)\right)}{t_1} + \dot{g}(\infty) e^{-i\omega t_N} + \sum_{k=2}^{N} \left(\frac{g_k - g_{k-1}}{t_k - t_{k-1}}\right)\left(e^{-i\omega t_{k-1}} - e^{-i\omega t_k}\right) \tag{3}$$

Lastly, the complex modulus is calculated using

$$G^*(\omega) = G'(\omega) + i\, G''(\omega) = \frac{\kappa}{6\pi a}\left(\frac{i\omega\hat{g}(\omega)}{1 - i\omega\hat{g}(\omega)}\right) \tag{4}$$

where $a$ is the radius of the particle (0.5 µm in the present case). For each trial, we extract two sets of $G'$ and $G''$ values at each frequency. The total number of $G'$ and $G''$ sets is ~18-30 sets for ~9-15 condensates. Finally, at each frequency, we average the values of $G'$ and $G''$and report the average with the error calculated as the standard deviation of the mean.

**Video particle tracking (VPT) nanorheology**

Multi-component peptide–$dT_{40}$–A1-LCD condensate systems consisting of 5 mg/mL peptide [either $(KGKGG)_5$, $(RGPGG)_5$, $(RPRPP)_5$, $(RGRGG)_5$, or $(RGYGG)_5$], 5 mg/mL $dT_{40}$, and 25 µM WT A1-LCD were prepared in the buffer containing 25 mM MOPS (pH 7.5), 150 mM NaCl, and 5 mM DTT. In addition, yellow-green carboxylate-modified polystyrene beads (200 nm in diameter, FluoSpheres, Invitrogen, F8811) were introduced in the sample buffer, at a low concentration (0.005% solids), prior to phase separation, to allow beads to passively partition into condensates once phase separation is initiated. The sample chambers were prepared similarly to what was described for the pMOT measurements. The sample-containing sandwich was then placed onto a Zeiss Primovert inverted microscope with a 100x oil-immersion objective lens. Condensates were allowed to settle on the glass slide for ~20 min prior to the start of imaging. For imaging, a Teledyne FLIR Blackfly S USB3 CMOS camera was used. The focus of the microscope was confined to a particular focal plane of the condensates for imaging. Videos were recorded for condensates containing multiple 200 nm beads for 1000 frames at a rate of 10 frames per second (with exposure time set to 100 ms) to record the motion of the particles within condensates. Three measurements were performed for three independent sample preparations for each multi-component condensate system.

In VPT nanorheology, the movement of the beads partitioned into the condensates is monitored over time. The monitored bead trajectories were analyzed to extract the mean square displacement (MSD) of the beads [12]. The MSDs were further analyzed to obtain the diffusion coefficients, nature of the diffusion, and the viscosity of the condensates. For tracking, we first employed the TrackMate plugin [13] in Fiji [6] to extract the diffusion of beads and to generate two-dimensional trajectories of each bead within condensates. By computing the center of mass vector $\boldsymbol{R}$ at each time point, we calculated the diffusion of the center of mass of the beads. The calculation of the center of mass vector was done using the particle velocities according to the following equation:

$$\boldsymbol{R_{COM}}(k) = \boldsymbol{R_0} + \sum_{k=0}^{k} \boldsymbol{\overline{v}_k} = \boldsymbol{R_0} + \sum_{k=0}^{k} \left( \frac{1}{N_k} \sum_{i=1}^{N} \boldsymbol{v_{i,k}} \right) \Delta k \tag{5}$$

Here, $\boldsymbol{R_0}$ is the initial center of mass vector calculated by averaging the coordinates of all beads in the first frame ($k$=1), $\boldsymbol{\overline{v}_k}$ is the mean velocity of all particles in the frame $k$, $\boldsymbol{v_{i,k}}$ is the velocity of the particle $i$ in frame $k$ in units of μm/frame, and $N_k$ is the number of particles in the frame $k$. The quantity $\Delta k$ is the frame difference, which is 1 in the present case. To ensure the absence of any sample drift in the data, the center of mass was calculated and subtracted from the individual bead trajectories. This was used to calculate the ensemble-averaged mean-squared displacement (MSD) using:

$$MSD(\tau) = \langle \boldsymbol{R}(t+\tau) - \boldsymbol{R}(t) \rangle |_{t,N} \tag{6}$$

In Eq. (6), $\tau$ is the lag time. Next, we extracted the diffusion coefficient of the beads by fitting the MSD to the following equation:

$$MSD(\tau) = 4D\tau^{\alpha} + N \tag{7}$$

Here, $D$ is the diffusion coefficient, $\alpha$ is the diffusivity exponent, and $N$ is a term to account for the tracking noise. For fluids with terminally viscous behavior, $\alpha$ approaches 1 at long lag-times, which allows for the calculation of the terminal viscosity using the Stokes-Einstein equation:

$$\eta = \frac{k_B T}{6\pi D r} \tag{8}$$

Here, $r$ is the particle radius, $k_B$ is the Boltzmann coefficient, and $T$ is the temperature in Kelvin. The averaged value was reported for the viscosity from different condensates over three independently prepared samples for each multi-component condensate system.

**Partition coefficient measurements in biomolecular condensates**

All images used to calculate partitioning of fluorescently labeled A1-LCD in peptide–ssDNA condensates were captured on a Q2 laser scanning confocal microscope (ISS Inc., 63x objective). All samples were prepared in 25 mM MOPS (pH 7.5), 150 mM NaCl, and 5 mM DTT, following the protocol described in the '*Preparation of multi-component biomolecular condensates for fluorescence imaging*' method section. For partition coefficient measurements of A1-LCD (WT or pathogenic variant), samples were prepared with 1 mg/mL of the indicated peptide, 1 mg/mL of the indicated nucleic acid, 25 μM of A1-LCD variant, and 100 nM of A1-LCD labeled with Alexa488. For partition coefficient measurements using A1-LCD Δhexa, 300 nM rhodamine red-labeled Δhexa was used. Partitioning of labeled protein was calculated as the ratio of intensity values within condensates and the average dilute phase intensity determined from 5 randomly chosen regions without condensates. Plots of partition coefficients represent measurements from 10 condensates from each of 3 independently prepared replicates.

**Client recruitment and retention assay**

Client recruitment and retention in biomolecular condensates were measured using a multi-channel microfluidics setup (u-Flux) integrated into the Lumicks C-Trap correlative optical tweezers–confocal fluorescence microscopy system. Briefly, peptide–nucleic acid condensates were prepared according to the protocol outlined in the '*Preparation of multi-component biomolecular condensates for fluorescence imaging*' method section, without including A1-LCD in the sample mixture; 0.5 mg/mL of both peptide and nucleic acid were used to generate condensates. The resultant binary condensates were subjected to laminar flow (~ 1 bar) for entry into a single channel (referred to as Ch 1). The experimental schematic is provided in **Fig. 5a**. The pressure is controlled using the Python-based Bluelake software (Lumicks) package. Condensates in Ch 1 were optically trapped using a laser tweezer operating at 10% trapping laser capacity (1064 nm), which corresponds to ~ 100 µW. The trapped condensate is then moved to the adjacent channel (Ch 2) containing the sample buffer only. At this point, the flow is adjusted to ~ 0.1 bar. This serves as the starting point of the fluorescence microscopy measurements. Using established waypoints, the trapped condensate is moved in a controlled manner at a speed of 10 µm/s from Ch 2 to the adjacent Ch 3, which contains Alexa488-labeled A1-LCD D262V (170 nM). Upon entry of the trapped condensate into Ch 3, fluorescence time-lapse videos were recorded continuously (0.7 frames per second) to monitor the recruitment of the fluorescently labeled protein client into the peptide–nucleic acid condensate system. This is followed for 100 sec after the condensate reaches the destined waypoint in Ch 3 to capture recruitment kinetics. Afterward, the condensate containing A1-LCD D262V is moved back to Ch 2 via the same path and speed. Continuous fluorescence imaging (0.7 frames per second) is conducted throughout to monitor the retention of A1-LCD D262V in the condensate. The retention kinetics were probed for a maximum of ~400 sec after the condensate reached the Ch 2 waypoint, i.e., the starting point of the measurements. In the case of the client recruitment and retention assay conducted using tetramethylrhodamine (TMR)-labeled dextran 4.4K, all steps were performed similarly except for the use of TMR-labeled dextran 4.4K (500 nM) in Ch 3 instead of Alexa488-labeled A1-LCD D262V.

For analysis of client recruitment and retention kinetics, the corresponding fluorescence time-lapse video was imported into Fiji [6] (v1.54f). Using a circular ROI, the fluorescence signal intensity of the client (either Alexa488-labeled A1-LCD D262V or TMR-labeled dextran 4.4K) was measured as a function of time using the 'Plot Z-axis profile' function in Fiji [6], for both the recruitment as well as retention stages of the assay. At either stage, the measured intensities were normalized according to the maximum value, after performing background subtraction. GraphPad Prism 10 was used for generating the plots of fluorescence intensity versus time, representing the recruitment as well as the retention stages of the assay. Further, fitting with a stretched exponential function,

$$Y = Y_0 * e^{-\left(\frac{X}{\tau}\right)^{\beta}} \tag{9}$$

was performed on the data generated from the retention stage to determine the efflux time constant ($\tau$). We note that efflux kinetics were better described by a stretched exponential than by a single exponential. This indicates a distribution of escape times rather than a single characteristic rate constant. Quantitative analyses were performed based on data collected across multiple condensates representing at least 3 independently prepared samples. The lookup table (LUT) Fire was used for the representation of fluorescence time-lapse videos and images showing client recruitment and retention dynamics to better distinguish changes in client partitioning. For preparing representative images and videos of client recruitment and retention, the raw time-lapse videos were upscaled via bilinear interpolation using Fiji [6] to enhance clarity for better visualization.

**Mammalian cell culture and live-cell imaging for flicker spectroscopy**

U2OS cells (a gift from Dr. Shahar Sukenik) were maintained in Dulbecco's Modified Eagle Medium (DMEM; ThermoFisher, 11995-065) supplemented with 10% fetal bovine serum (FBS; ThermoFisher, A5669801) at 37°C in a humidified incubator with 5% $CO_2$. For flicker spectroscopy analysis, cells were seeded into 8-well chambered coverslips and allowed to grow for approximately 24 hours prior to transfection.

Cells were transiently transfected with plasmids encoding HaloTag-G3BP1 together with either hnRNPA1-D262V–IRES-GFP or hnRNPA1-allW-D262V–IRES-GFP. Details of the full-length hnRNPA1 variants are described in our previous work[2]. HaloTag-G3BP1 sequence was synthesized by Genscript and was cloned into mEGFP-C1 vector by swapping out the mEGFP sequence (addgene.org/54759/; mEGFP-C1 was a gift from Michael Davidson). Approximately 24 hours post-transfection, cells were incubated with JF646 HaloTag (Promega, GA1120) ligand in FluoroBrite medium (ThermoFisher, A1896701) for 15 minutes to label Halo-G3BP1, followed by two washes with FluoroBrite medium to remove excess ligand.

Following labeling, cells were treated with 500 µM sodium arsenite (LabChem LC228709) to induce stress granule formation. Imaging was carried out 45 minutes after treatment. During imaging, cells were maintained at 37°C and 5% $CO_2$ in an environmental chamber. Live-cell imaging was performed using an Andor Dragonfly spinning disk confocal microscope equipped with a 100×, 1.49 NA objective and a 1024 × 1024-pixel iXon EMCCD camera. Time-lapse images were acquired in finite burst mode at 50 ms intervals for a total of 1,000 frames. Illumination was performed using the Power Density 1 (PD1) mode with a laser power of 10% and an EM gain of 150. Cells expressing hnRNPA1 variants were identified based on GFP fluorescence intensity and were then selected for finite burst imaging and downstream analyses.

**Flicker spectroscopy image analysis**

Time-lapse image series were analyzed using the FlickerPrint software package[14] to extract condensate interfacial tension (σ) and bending rigidity (κ) from thermal fluctuations of SG boundaries utilizing HaloTag-G3BP1 (JF646) fluorescence. Raw Imaris (.ims) files were converted to ImageJ-format TIFF for subsequent background subtraction. Rolling-ball background subtraction was applied to each frame (radius 50 px ≈ 6.3 µm) to remove fluorescence background before condensate detection.

Stress granules were detected via FlickerPrint using the gradient method with Gaussian smoothing (σ = 1.5 px), restricting detection to objects with radii of 0.3–3.0 µm and a minimum normalized intensity of 0.20, with a fill threshold of 0.6 and a frame-to-frame tracking threshold of 15 px. Fourier-mode spectra were fitted using the corrected experimental spectrum over 15 Fourier modes at 37°C with a pixel size of 0.1254 µm per pixel.

Granules were retained for downstream analysis if they satisfied all of the following quality criteria: $\sigma > 1 \times 10^{-10}$ N/m, spectrum fitting difference > 0.03, pass rate > 0.6, fitting error < 0.5, and all six lowest-order Fourier modes above the pixel-noise floor, $(\text{pixel-size}/15)^2 / \text{radius}^2$.

**Preparation of whole cell lysate and lysate granules, and imaging of reconstituted granules**

U2OS cells expressing G3BP1-tdTomato from the endogenous locus (a gift from Dr. J. Paul Taylor) were maintained in DMEM supplemented with 10% FBS, 2 mM L-glutamine (Gibco, cat # A2916801), and 1% penicillin-streptomycin. Cells were cultured to confluency, harvested using TrypLE (Gibco, cat # 12604021), washed with 1 X PBS, and stored as pellets at −80°C following PBS removal. For the preparation of lysate and lysate granules, the protocol described in an earlier report from Freibaum *et al.* [15], was used with slight modifications. Briefly, frozen cell pellets (three vials containing 5 × $10^6$ cells each) were thawed at room temperature and resuspended in 250–300 µl lysis buffer containing 50 mM Tris-HCl (pH 7.0), 0.5% NP-40, protease inhibitor cocktail, and 2.5% murine RNase inhibitor (Fisher scientific, cat# 50-995-237). Pellets were pooled and homogenized with gentle pipetting until fully resuspended, followed by incubation at room temperature for 3 min. Lysates were clarified by centrifugation at 21,000 × g for 5 min at room temperature, and the supernatant was transferred to a fresh microcentrifuge tube. Protein concentration was measured using a BCA assay with BSA as the standard. Lysate granules were prepared by combining lysate at a final protein concentration of 1 mg/ml with purified G3BP1 (20 µM) to induce phase separation. Depending on the experimental conditions, D262V A1-LCD (20 µM), allW D262V A1-LCD (0.1 µM or 0.2 µM), and/or polyA RNA (200 ng/µl) were added. All samples were prepared in 50 mM Tris buffer (pH 7.0), while maintaining the NaCl concentration at 150 mM throughout.

N-terminal labeling was performed in denaturing conditions (4 M guanidinium chloride, 100 mM phosphate buffer, pH 7.0) using Alexa Fluor 647 NHS ester (Invitrogen, A20006) for D262V A1-LCD and Oregon Green 488 (Invitrogen, O6149) for allW D262V. Desalting steps were conducted to remove excess free dye. For imaging experiments, the labeled protein was mixed with the corresponding unlabeled protein at a 1:100 molar ratio and subsequently buffer exchanged into native buffer to use for sample preparation. Imaging was carried out on Sigmacote (Sigma-Aldrich SL2) coated glass slides using an i4-Nikon AX laser scanning confocal microscope with 488 nm and 633 nm laser excitation for Oregon Green 488 and Alexa Fluor 647, respectively. G3BP1-tdTomato expressed from the endogenous locus of the U2OS cells used for lysate preparation was imaged using excitation at 561 nm. All imaging experiments were performed at room temperature.

**Phase-separation propensity and LARKS in human IDRs**

For each sequence in the LARKS database[16], the human IDR with a contiguous fully matching segment was identified across the human IDRome[17]. Six-residue tiles within the matching segment were predicted to form LARKS if their Rosetta energies in the LARKS database[18] satisfied at least one of the following criteria for the backbone templates: FUS-STGGYG < 4.0, hnRNPA1-GYNGFG < 10.0, or FUS-SYSGYS < −3.0. The number of predicted LARKS tiles was averaged over entries in the LARKS database mapping to the same IDRome sequence, and the LARKS fraction of the human IDR, $f_{\mathrm{LARKS}}$, was calculated as the mean number of LARKS tiles divided by sequence length.

Free energies of transfer from the dilute into the dense phase of homotypic biomolecular condensates ($\Delta G_{\mathrm{pred}}$) were predicted using a machine-learning predictor trained on CALVADOS 2 simulation data[19]. IDRs were grouped into $\Delta G_{\mathrm{pred}}$ bins with a width of 0.5 $k_B T$, and the mean and SEM of $f_{\mathrm{LARKS}}$ were calculated for each bin. Spearman correlation coefficients between $\Delta G_{\mathrm{pred}}$ and $f_{\mathrm{LARKS}}$ were calculated for subsets of IDRs with $\Delta G_{\mathrm{pred}}$ below threshold values ranging from −3 to −6 $k_B T$. The shift between distributions of $f_{\mathrm{LARKS}}$ for IDRs with $\Delta G_{\mathrm{pred}} < -4\ k_B T$ and $\Delta G_{\mathrm{pred}} \geq -4\ k_B T$ was quantified using Cohen's $d$, with standard errors estimated through $10^4$ bootstraps[17]. The statistical significance of LARKS enrichment in IDRs with $\Delta G_{\mathrm{pred}} < -4\ k_B T$ was assessed using a one-sided Brunner–Munzel test, with the *P* value calculated from the *t*-distribution as implemented in SciPy[20] (v1.15.3).

**Software**

For image processing and quantification, Fiji[6] (v1.54f) was used. Python scripts were utilized for analyzing ThT kinetics as well as VPT nanorheology-based MSD measurements and pMOT measurements, which are available on GitHub (github.com/BanerjeeLab-repertoire/Interface-density-and-viscoelasticity-of-heterotypic-condensates-determine-their-ability-to-suppress).

Scripts used for flicker spectroscopy analysis were obtained from Williamson, et al. [14]. Measurements concerning FRAP and condensate recruitment and retention assay, for which fluorescence images were acquired using the Lumicks C-trap correlative laser tweezer and confocal fluorescence microscopy system, Bluelake (v1.6.11; https://lumicks.github.io/bluelake-api/2.4.0/index.html) software was used. For the acquisition of fluorescence images using the ISS Q2 laser scanning confocal microscope, Vistavision (v4.2, ISS Inc.) was used. For the acquisition of fluorescence images using the Andor Dragonfly 600 confocal microscope, Fusion (v2.3) was used. Additionally, Imaris (v10.1, Bitplane) was used for processing images acquired using the Andor Dragonfly 600 confocal microscope. For statistical representation, GraphPad Prism 10 (Version 10.2.3) and IGOR Pro (v6.3 beta 1) were used. For aid in estimation of statistical significance of LARKS enrichment in IDRs, SciPy[20] (v1.15.3) was utilized. Artwork from NIH BIOART was adapted to make certain schematics (NIAID NIH BIOART Source: bioart.niaid.nih.gov/bioart/456). Adobe Illustrator CC (2026) was used for the assembly of all figures. ThT curves were smoothed in RStudio (Version 2023.12.1+402).

### Supplementary Figures

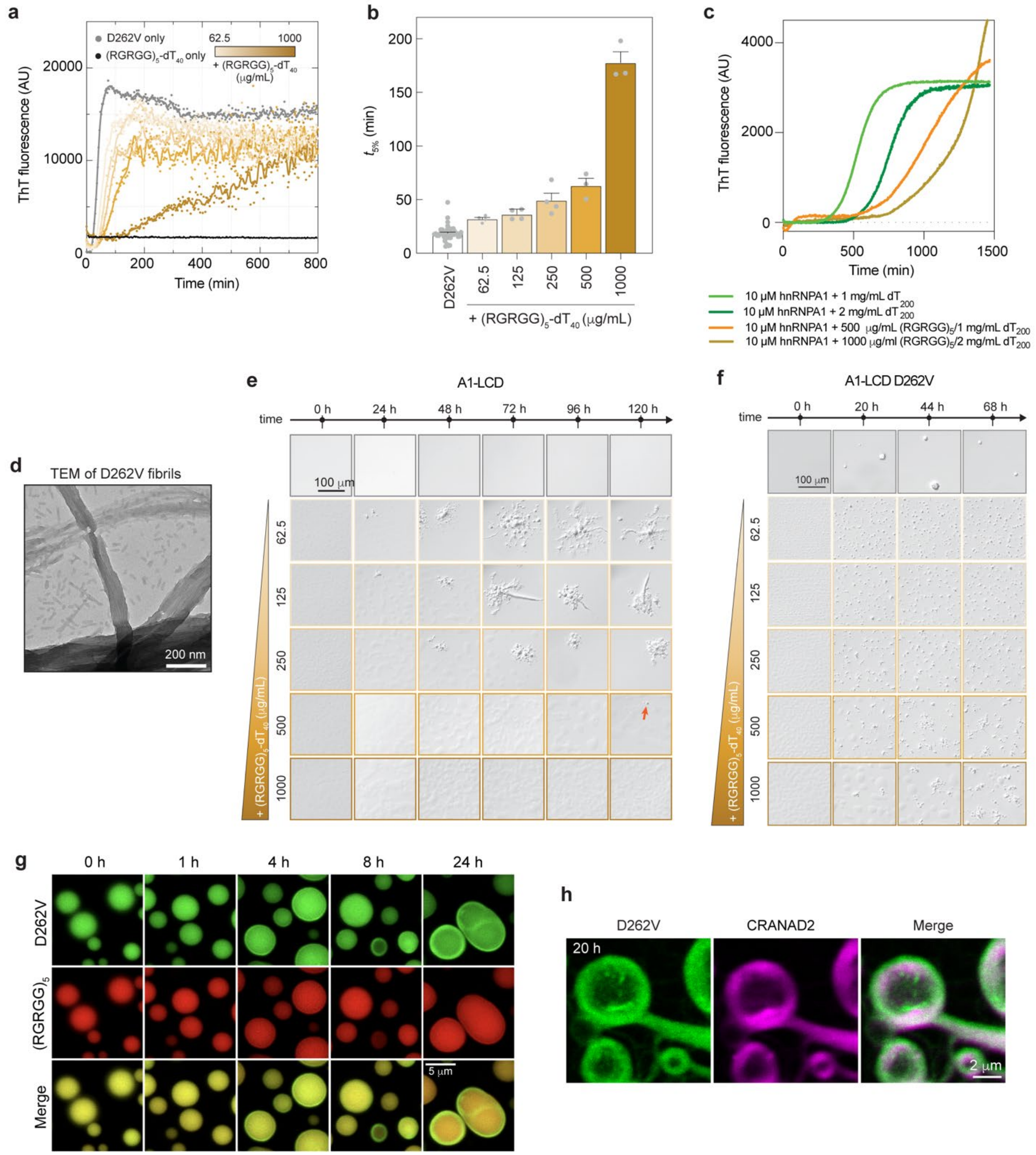


**Figure S1. Multi-component biomolecular condensates suppress fibril formation.** (**a**) Kinetics of fibril formation monitored by ThT fluorescence of 25 μM A1-LCD D262V either alone or in the presence of increasing concentrations of $(RGRGG)_5$ and $dT_{40}$. $(RGRGG)_5$-$dT_{40}$ condensates alone do not exhibit strong enhanced ThT signal as shown by the black curve. A smoothing window of 10 data points was applied to generate a connecting line. **(b)** Lag times ($t_{5\%}$) extracted from the ThT kinetics shown in (**a**) and from replicate experiments ($n \geq 3$) are shown along with the mean ± SEM. **(c)** Kinetics of fibril formation monitored by ThT fluorescence of full-length hnRNPA1 with ssDNA, or in the presence of increasing concentrations of $(RGRGG)_5$ and $dT_{200}$. An excess of $dT_{200}$ was added because full-length hnRNPA1 binds ssDNA and solubilizes condensates at mass equivalents of ssDNA and peptide. **(d)** nsTEM image of A1-LCD D262V fibrils from the sample without condensates captured after 6 days of incubation at 20 ℃ with shaking. Time-course DIC images of **(e)** A1-LCD or **(f)** A1-LCD D262V, either alone or in the presence of increasing volume fractions of $(RGRGG)_5$–$dT_{40}$ condensates. **(g)** Time course fluorescence images to monitor the progressive loss of A1-LCD D262V (visualized with 250 nM Alexa488-D262V) from the interiors of $(RGRGG)_5$–$dT_{40}$ condensates [visualized with 250 nM Alexa594-$(RGRGG)_5$], leaving behind a protein rim enriched in D262V. Images were adjusted independently for optimal visualization of the coexistence of droplets and fibrils in each condition. **(h)** Confocal fluorescence microscopy images of $(RGRGG)_5$-$dT_{40}$ condensates with A1-LCD D262V at a late time point showing formation of CRANAD2-positive amyloid fibrils containing D262V (visualized using 250 nM Alexa488-labeled D262V).

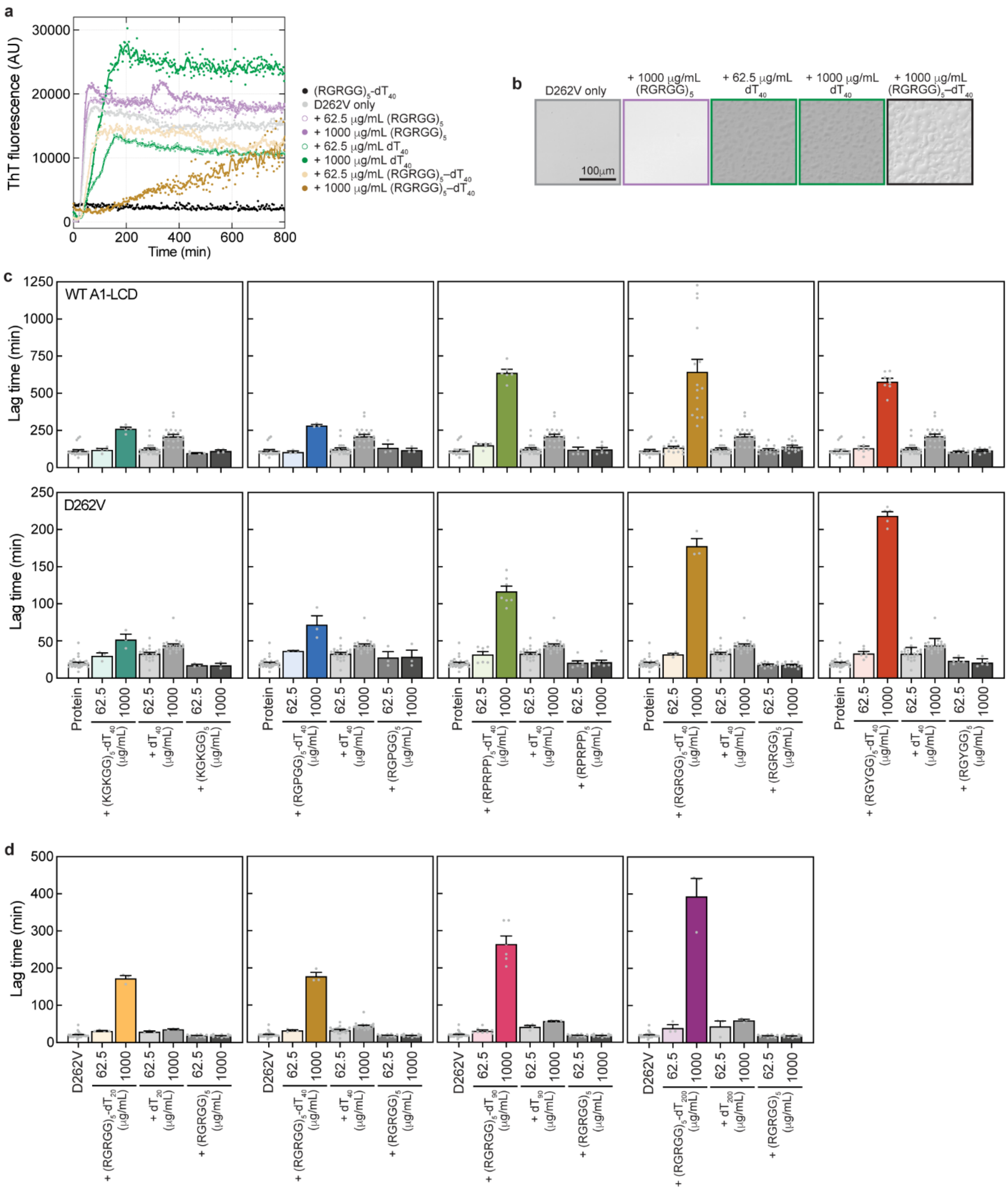


**Figure S2. Increase in A1-LCD fibril formation lag times cannot be reproduced using peptide or DNA alone. (a)** Kinetics of fibril formation monitored by ThT fluorescence of 25 µM A1-LCD D262V either alone or in the presence of either $(RGRGG)_5$–$dT_{40}$ condensates or peptide or ssDNA alone at two different concentrations. Baseline subtraction was applied, and a smoothing window of 10 data points was used to generate a connecting line. **(b)** Brightfield images of 25 µM A1-LCD D262V either alone or in the presence of either $(RGRGG)_5$–$dT_{40}$ condensates or peptide or ssDNA alone at two different concentrations. It is evident that at a concentration of 25 µM, which is much below the saturation concentration for homotypic condensation, A1-LCD and $dT_{40}$ form heterotypic condensates. **(c)** Lag times ($t_{5\%}$) extracted from the ThT kinetics corresponding to either WT A1-LCD (top) or D262V (bottom) alone or in the presence of either multi-component condensate systems of varying peptide sequence grammar or peptide or DNA alone. Lag times ($t_{5\%}$) from replicate experiments ($n \geq 3$) are shown along with the mean ± SEM. **(d)** Lag times ($t_{5\%}$) extracted from the ThT kinetics corresponding to D262V either alone or in the presence of either multi-component condensate systems of varying nucleic acid (dT) length or peptide or DNA alone. Lag times from replicate experiments ($n \geq 3$) are shown along with the mean ± SEM.

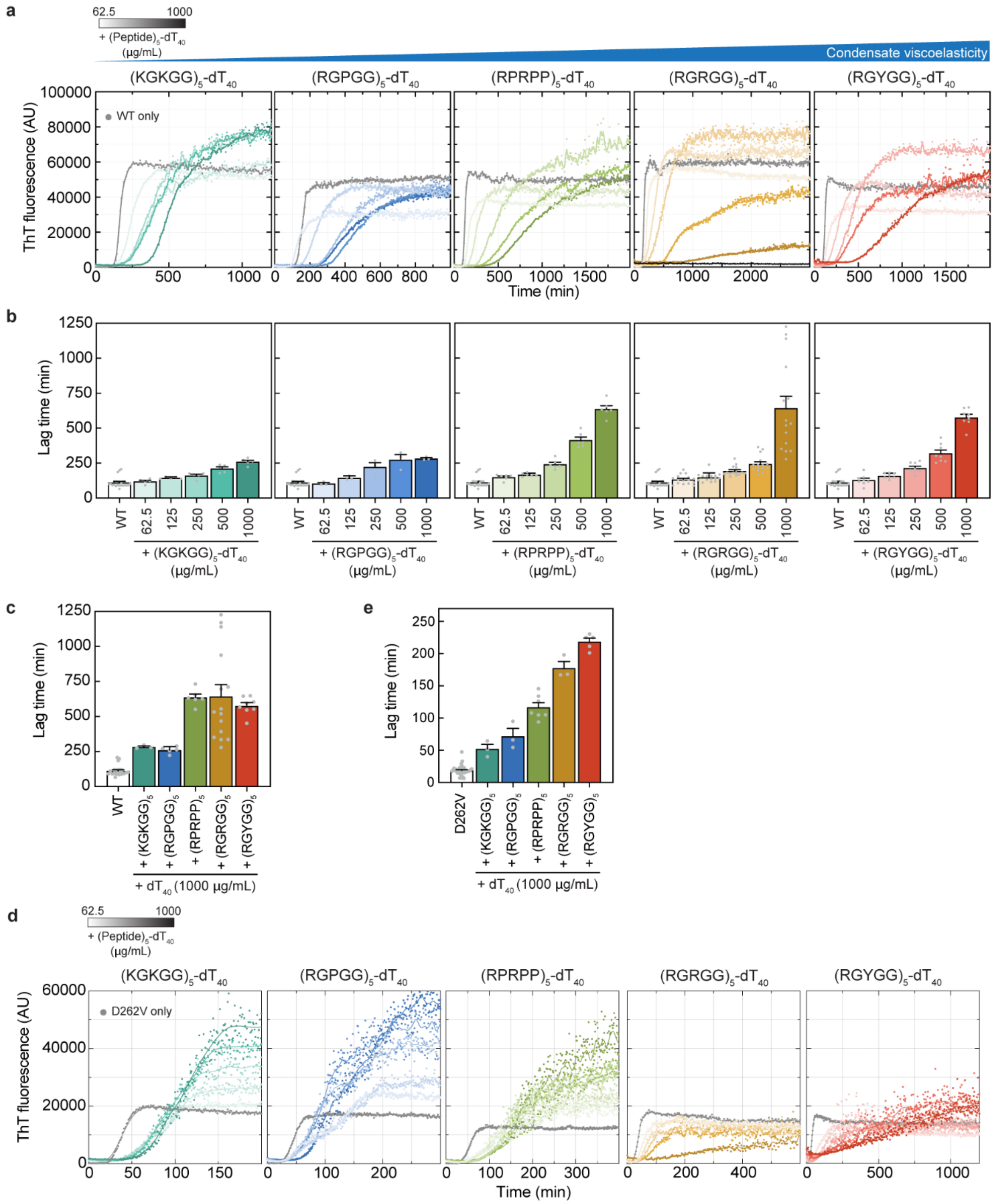


**Figure S3. Multi-component biomolecular condensate systems with higher viscoelasticity result in longer lag times for A1-LCD fibril formation. (a)** Kinetics of fibril formation monitored by ThT fluorescence of 25 µM WT A1-LCD either alone or in the presence of different peptide–ssDNA multi-component condensate systems with distinct viscoelasticity at a range of volume fractions, arranged in order of increasing condensate viscoelasticity from left to right. Across the different condensate systems, the repeat peptides are varied, but $dT_{40}$ is kept constant. A smoothing window of 10 data points was applied to generate a connecting line. **(b)** Lag times ($t_{5\%}$) extracted from the ThT kinetics corresponding to (**a**) for WT A1-LCD either alone or in the presence of multi-component condensate systems. Lag times from replicate experiments ($n \geq 3$) are shown along with the mean ± SEM. **(c)** A subset of the lag times shown in (**b**), corresponding to the highest dense phase volume fraction. **(d)** Kinetics of fibril formation monitored by ThT fluorescence of 25 μM A1-LCD D262V either alone or in the presence of different multi-component condensate systems with distinct viscoelasticity at a range of volume fractions, arranged in order of increasing condensate viscoelasticity from left to right. A smoothing window of 10 data points was applied to generate a connecting line. See the corresponding $t_{5\%}$ plots for all peptide–ssDNA concentrations tested in **Fig. 2d**. **(e)** Lag times ($t_{5\%}$) for the highest peptide–ssDNA concentrations extracted from the ThT kinetics shown in **Fig. S3d** and from replicate experiments ($n \geq 3$) are shown along with the mean ± SEM.

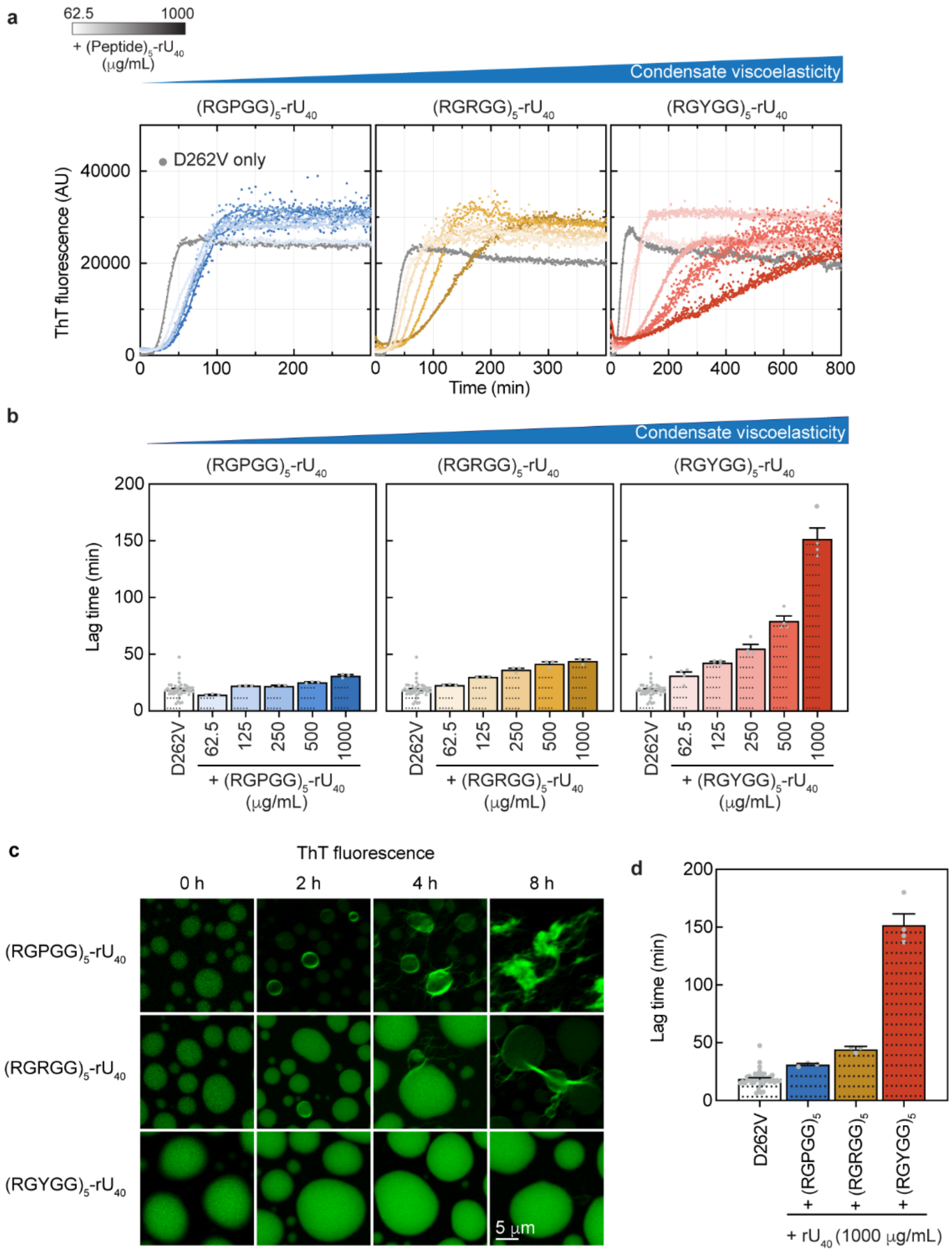


**Figure S4. Multi-component peptide–RNA biomolecular condensate systems with higher viscoelasticity result in longer lag times for A1-LCD fibril formation. (a)** Kinetics of fibril formation monitored using ThT fluorescence of 25 μM A1-LCD D262V either alone or in the presence of different peptide–ssRNA multi-component condensate systems with distinct viscoelasticity at a range of volume fractions, arranged in order of increasing condensate viscoelasticity from left to right. Across the different condensate systems, the repeat peptides were varied, but $rU_{40}$ was kept constant. A smoothing window of 10 data points was applied to generate a connecting line. **(b)** Lag times ($t_{5\%}$) extracted from the ThT kinetics in (**a**) of A1-LCD D262V either alone or in the presence of multi-component repeat peptide–RNA (i.e., $rU_{40}$) condensates. Lag times from replicate experiments ($n \geq 3$) are shown along with the mean ± SEM. **(c)** Time course fluorescence images of different peptide–$rU_{40}$ condensate systems, in which the peptide component is varied. ThT was used for the visualization of amyloid fibrils. Images were adjusted independently for optimal visualization of the coexistence of droplets and fibrils in each condition. **(d)** A subset of lag times ($t_{5\%}$) shown in (**b**), corresponding to the highest dense phase volume fraction from replicate experiments (n ≥ 3), are shown along with the mean ± SEM.

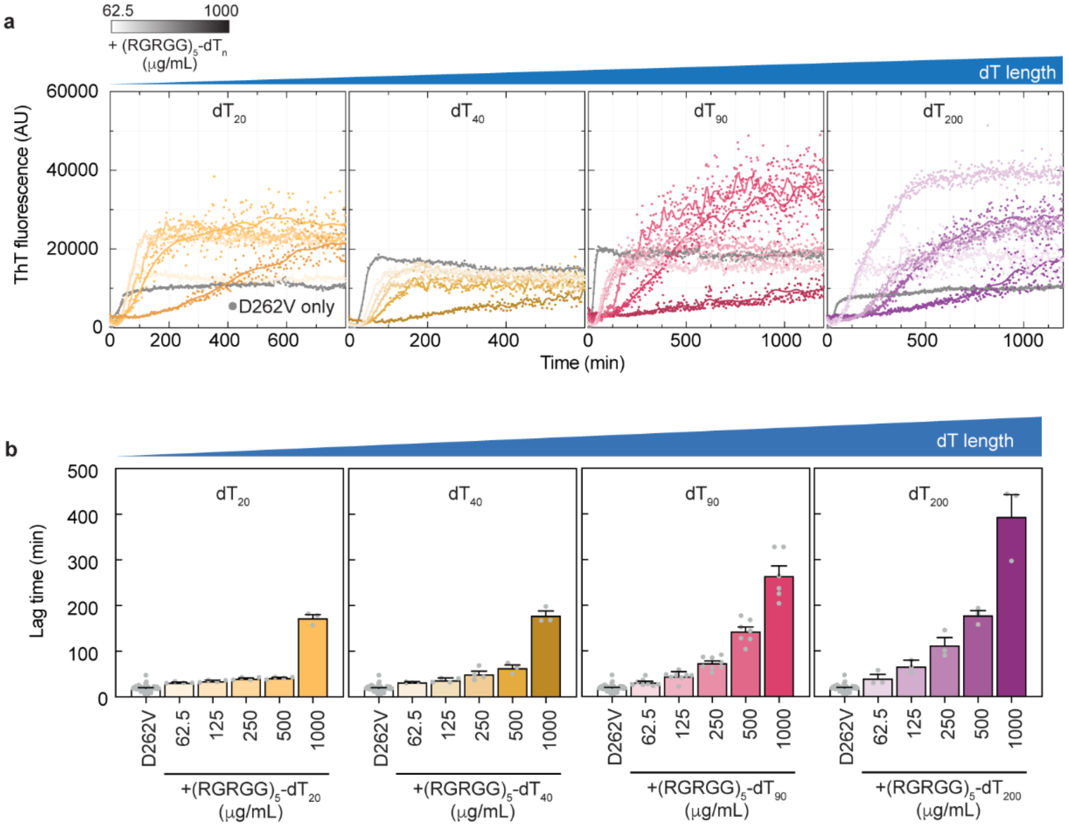


**Figure S5. Multi-component biomolecular condensate systems scaffolded by different nucleic acid lengths show that higher viscoelasticity results in longer lag times for A1-LCD fibril formation. (a)** Kinetics of fibril formation monitored by ThT fluorescence of A1-LCD D262V alone and in the presence of different peptide–ssDNA multi-component condensate systems with distinct viscoelasticity at a range of peptide/ssDNA concentrations, arranged in order of increasing condensate viscoelasticity from left to right. Across the different condensate systems, the identity of the peptide component, $(RGRGG)_5$, was kept constant while varying the chain length of the ssDNA (poly-dT). A smoothing window of 10 data points was applied to generate a connecting line. **(b)** Lag times ($t_{5\%}$) extracted from ThT kinetics monitoring D262V fibril formation either in the absence or presence of increasing concentrations of $(RGRGG)_5$ and poly-dT ssDNA of distinct chain lengths. Data from replicate experiments ($n \geq 3$) are shown along with the mean $\pm$ SEM.

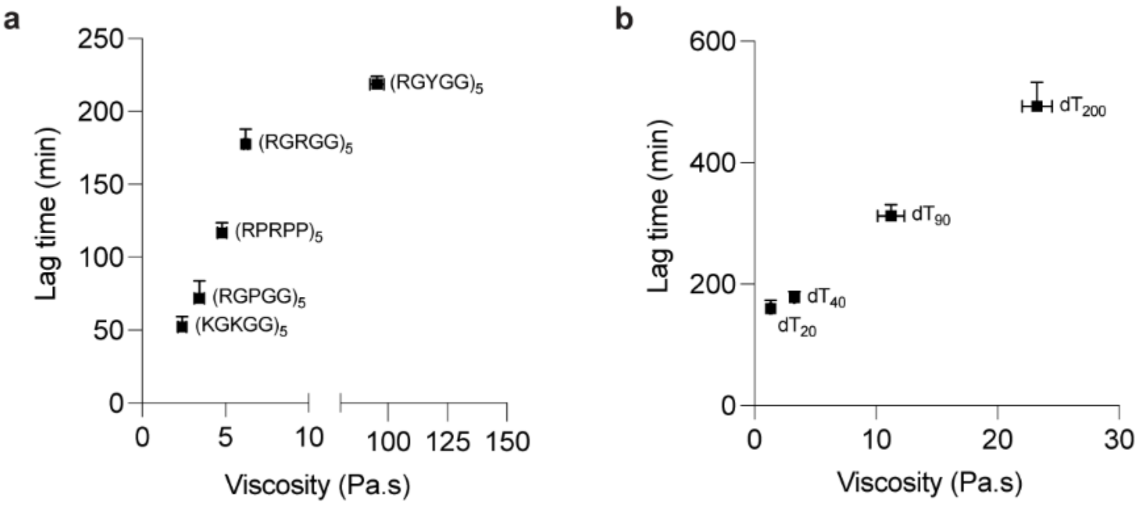


**Figure S6. Lag time for fibril formation scales with condensate viscosity. (a)** Lag times ($t_{5\%}$; mean ± SEM from n ≥ 3 replicate experiments) for the highest peptide–ssDNA concentrations extracted from the ThT kinetics shown in **Fig. S3d** as a function of viscosity based on 3-component condensate systems (peptide–$dT_{40}$–A1-LCD) wherein the peptide sequence grammar is varied. **(b)** Lag times ($t_{5\%}$; mean ± SEM from n ≥ 3 replicate experiments) for the highest peptide–ssDNA concentrations extracted from the ThT kinetics shown in **Fig. S5a** as a function of viscosity based on 2-component condensate systems (peptide–$dT_{40}$) wherein the nucleic acid chain length is varied. The data is represented as mean ± SEM.

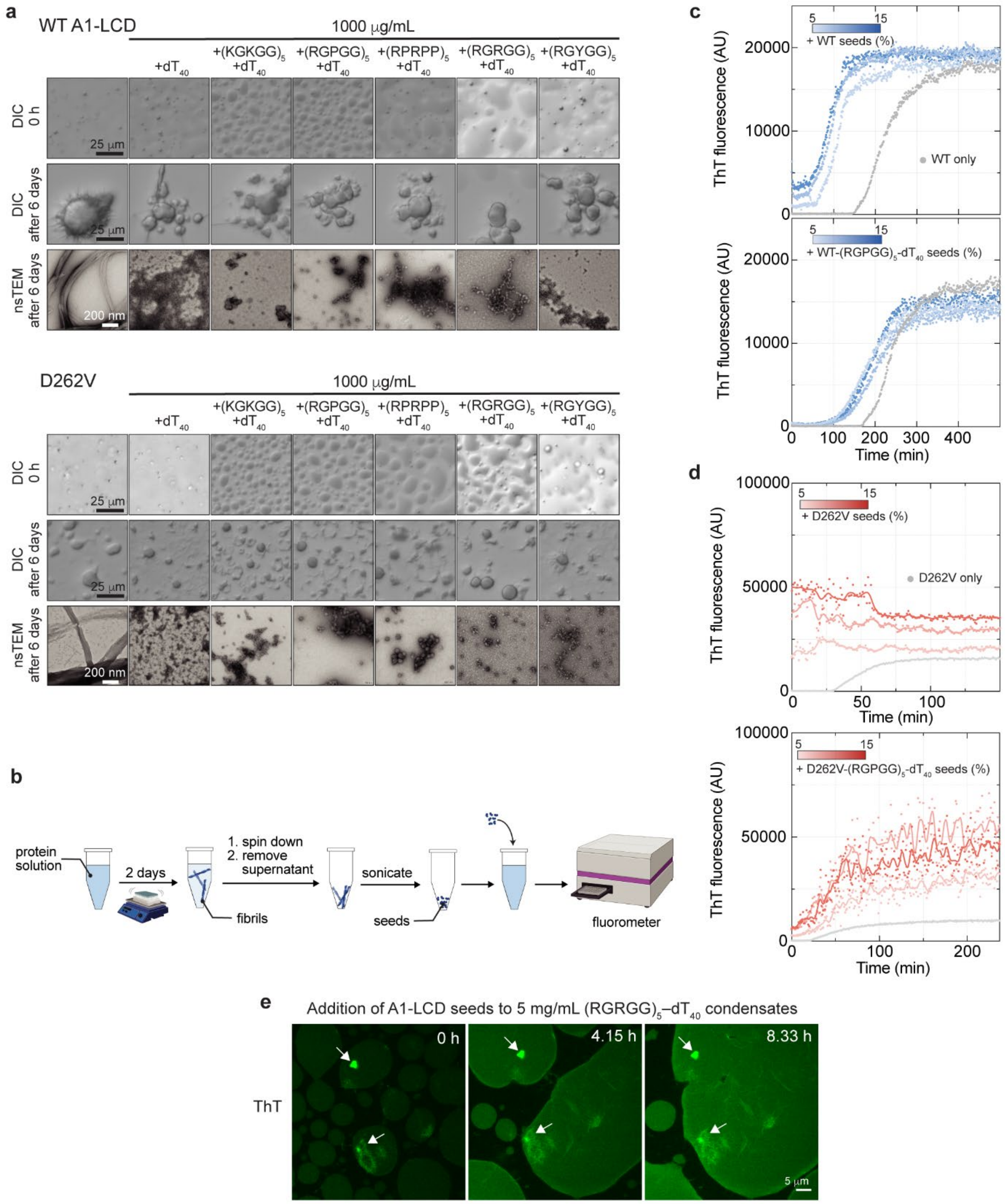


**Figure S7. WT A1-LCD and D262V form small assemblies in the presence of multi-component peptide–nucleic acid condensates, which can seed fibril formation of A1-LCD in the dilute phase but not within condensates**. **(a)** DIC and nsTEM images show morphology of micron-scale fibrillar bundles and nanoscale fibrils, respectively, of A1-LCD (top) and D262V (bottom) either in the absence or presence of condensates, as indicated. nsTEM images were captured after 6 days of incubation at 20 ℃ while shaking. In parallel, light microscopy images were taken at the beginning of the experiment and after 6 days to have a direct comparison with nsTEM images. **(b)** Schematic of seed generation and seeding assay. 'SN' refers to supernatant. ThT kinetics of WT A1-LCD (**c**) and D262V (**d**), either alone or in the presence of seeds at varying concentrations. **(e)** Introducing WT A1-LCD seeds generated from samples without condensates during the sample preparation stage of 5 mg/mL $(RGRGG)_5$–$dT_{40}$ condensates containing 25 μM WT A1-LCD results in passive entry of seeds into the dense phase (visualized using ThT). However, we do not observe the growth of these ThT-positive assemblies (indicated by white arrows) in the dense phase as a function of time. Note the coarsening of condensates in the frame of view as a function of sample age.

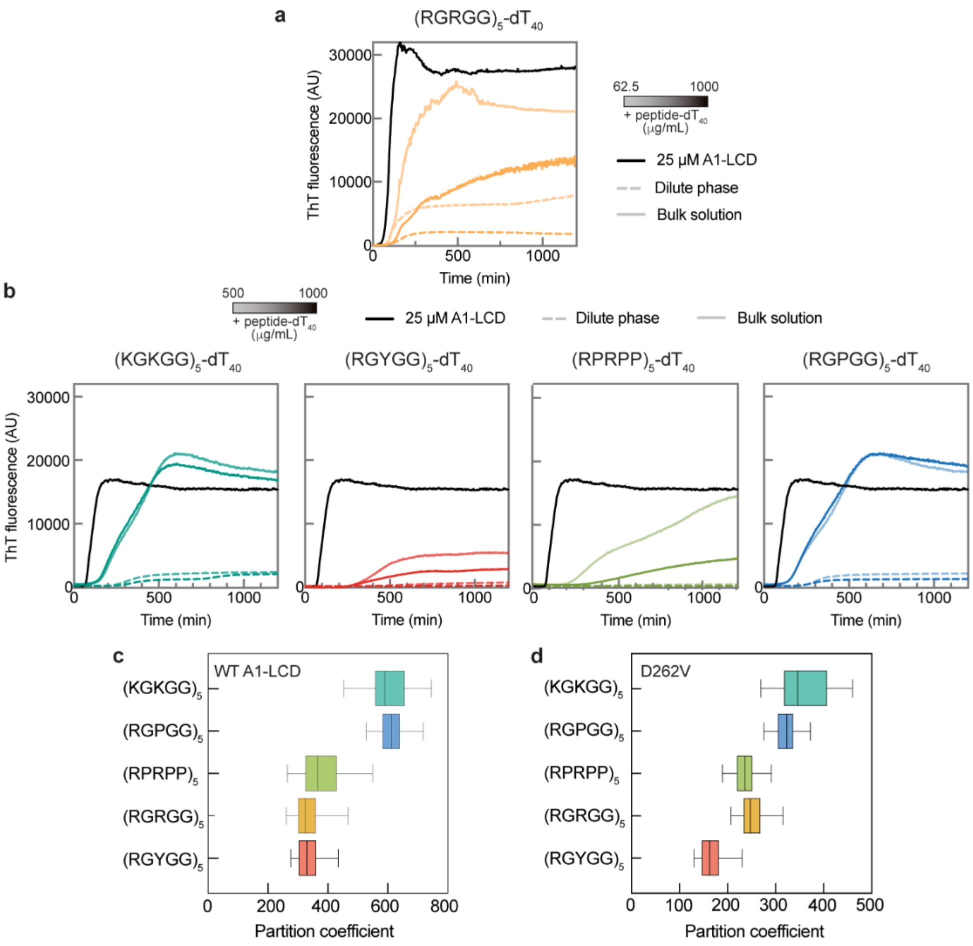


**Figure S8. Kinetics of fibril formation in different fractions and A1-LCD partitioning profiles across different condensate systems.** Kinetics of fibril formation monitored by ThT fluorescence of either the fractionated dilute phase or the total bulk sample consisting of 25 μM A1-LCD, either alone or in the presence of **(a)** (RGRGG)$_5$–dT$_{40}$ condensates, or in the presence of **(b)** different multi-component condensate systems with distinct viscoelasticity at two different volume fractions. The A1-LCD alone ThT curve (black) has been reproduced across all the ThT plots shown in panel (b). Partition coefficients of Alexa488-labeled WT A1-LCD **(c)** and D262V **(d)** into various multi-component condensate systems of distinct viscoelastic properties at peptide/ssDNA concentrations of 1000 μg/mL. The central line of the box plot represents the median, while the whiskers represent the minimum and maximum.

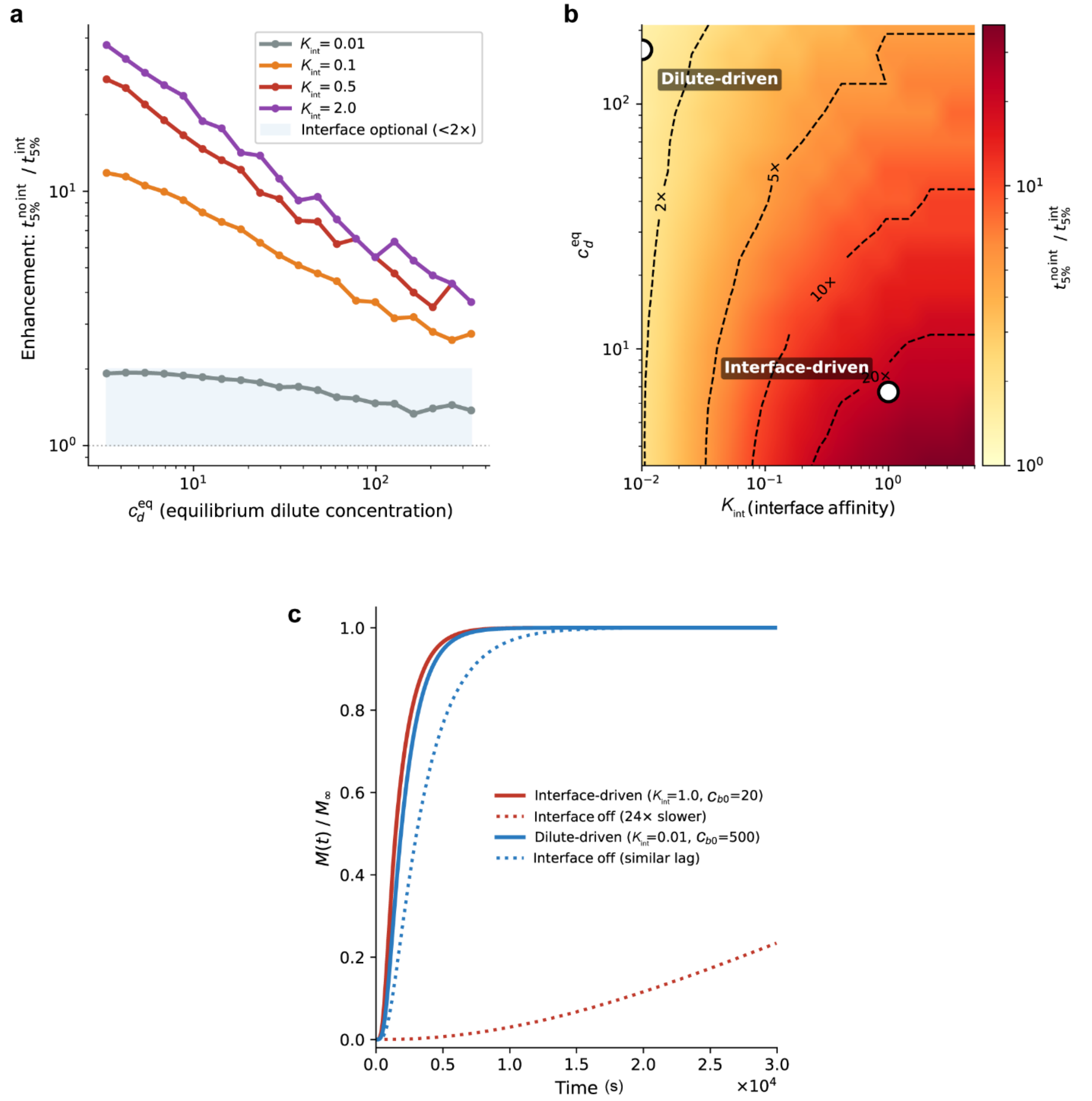


**Figure S9. Separate regimes of fibril formation driven by nucleation at interfaces versus in the dilute phase. (a)** Enhancement of fibril formation $E$ as a function of the equilibrium dilute concentration $c_d^{\text{eq}}$ for four interface affinities $K_{\text{int}}$. At low $c_d^{\text{eq}}$, the interface is essential ($E$ up to $\sim 10^2$); at high $c_d^{\text{eq}}$, nucleation becomes similar to interface nucleation, *i.e.*, $E$ approaches 1. **(b)** Two-dimensional enhancement map in the $(K_{\text{int}}, c_d^{\text{eq}})$ plane with iso-enhancement contours at $E = 2, 5, 10, 20$. The diagonal contour structure delineates the boundary between the interface-essential and dilute phase-driven regimes. **(c)** Two paired $M(t)/M_\infty$ traces with active interface (solid lines) and inactivated interface (dotted lines). In the *interface-driven* trajectory ($c_b(0) = 20$, $K_{\text{int}} = 1.0$), removing the active interface delays fibril formation by an order of magnitude or more. In the *dilute phase–driven* trajectory ($c_b(0) = 500$, $K_{\text{int}} = 0.01$), the curves for which the interface is on vs. off are nearly indistinguishable.

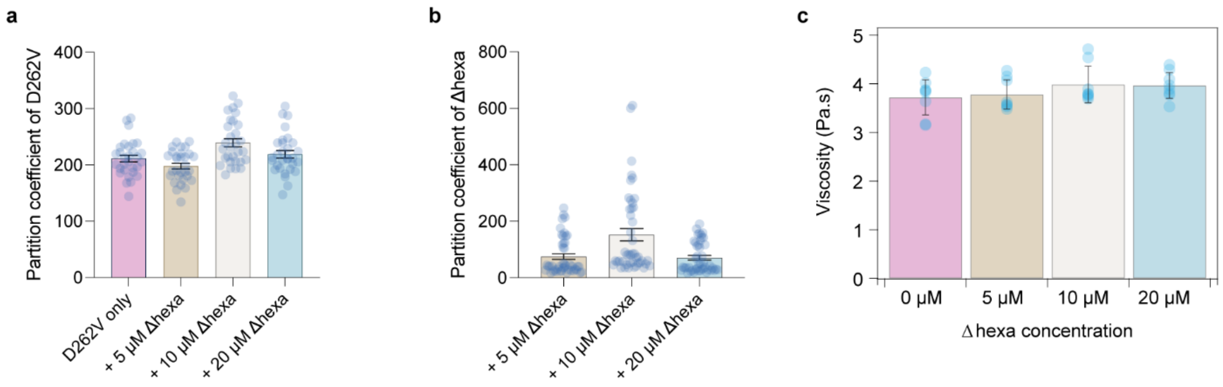


**Figure S10. Partitioning of A1-LCD D262V and ∆hexa in multi-component condensates as a function of ∆hexa concentration and its influence on condensate viscosity**. **(a)** Partition coefficients of Alexa488-labeled A1-LCD D262V in (RGRGG)$_5$–dT$_{40}$-D262V condensates with the addition of different total ∆hexa concentrations. **(b)** Partition coefficients of rhodamine red-labeled ∆hexa in (RGRGG)$_5$–dT$_{40}$-D262V condensates with different total ∆hexa concentrations. In each plot, the mean ± standard deviation (SD) is represented along with points indicating individual measurements based on three independent replicates. **(c)** Viscosity measurements made using VPT nanorheology of multi-component condensate systems from (**a**). In (**a**-**c**), individual points represent data points from three independent replicates except for (**c**), which represents two independent replicates.

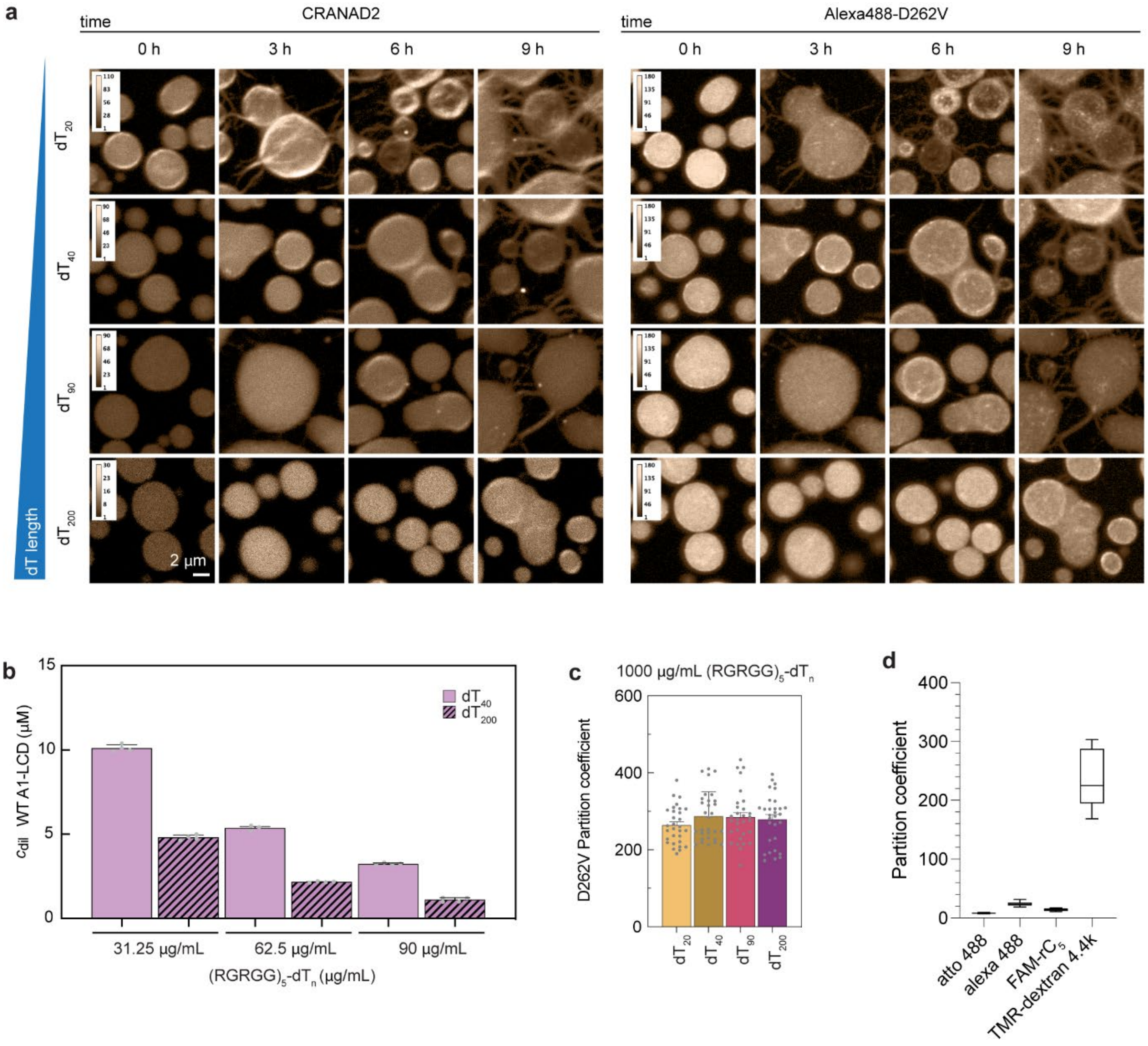


**Figure S11. Higher condensate viscoelasticity, independent of peptide sequence grammar, slows down D262V fibril formation and highlights the role of efflux in fibril assembly. (a)** Fluorescence images of multi-component condensates composed of $(RGRGG)_5$ and poly-dT of variable lengths, containing D262V (visualized with 250 nM Alexa488-D262V). Fibrils were visualized using CRANAD2. In each condition, the display range of the image was adjusted independently for optimal visualization of the morphological features of fibril assemblies, which are dimmer and therefore less visible than the substantially brighter condensates. Such adjustments were made uniformly across a series of images. **(b)** Dilute phase concentrations ($c_{dilute}$) of WT A1-LCD with either $(RGRGG)_5$-$dT_{40}$ or $(RGRGG)_5$-$dT_{200}$ condensate systems, at increasing peptide-ssDNA concentrations, show increased A1-LCD sequestration in condensates. The data is reported as mean ± SEM along with points indicating individual measurements from three independent replicates. **(c)** Partition coefficients of Alexa488-labeled D262V in peptide–ssDNA condensates [$(RGRGG)_5$ and poly-dT of increasing chain length] at fixed concentrations of 1 mg/mL peptide and nucleic acid. The points represent individual measurements from three independent replicates and are reported along with mean ± SD. **(d)** Partition coefficient measurements of different fluorescent clients in $(RGRGG)_5$-$dT_{40}$ condensates. The central line of the box plot represents the median, while the whiskers represent the minimum and maximum, based on measurements from three independent replicates.

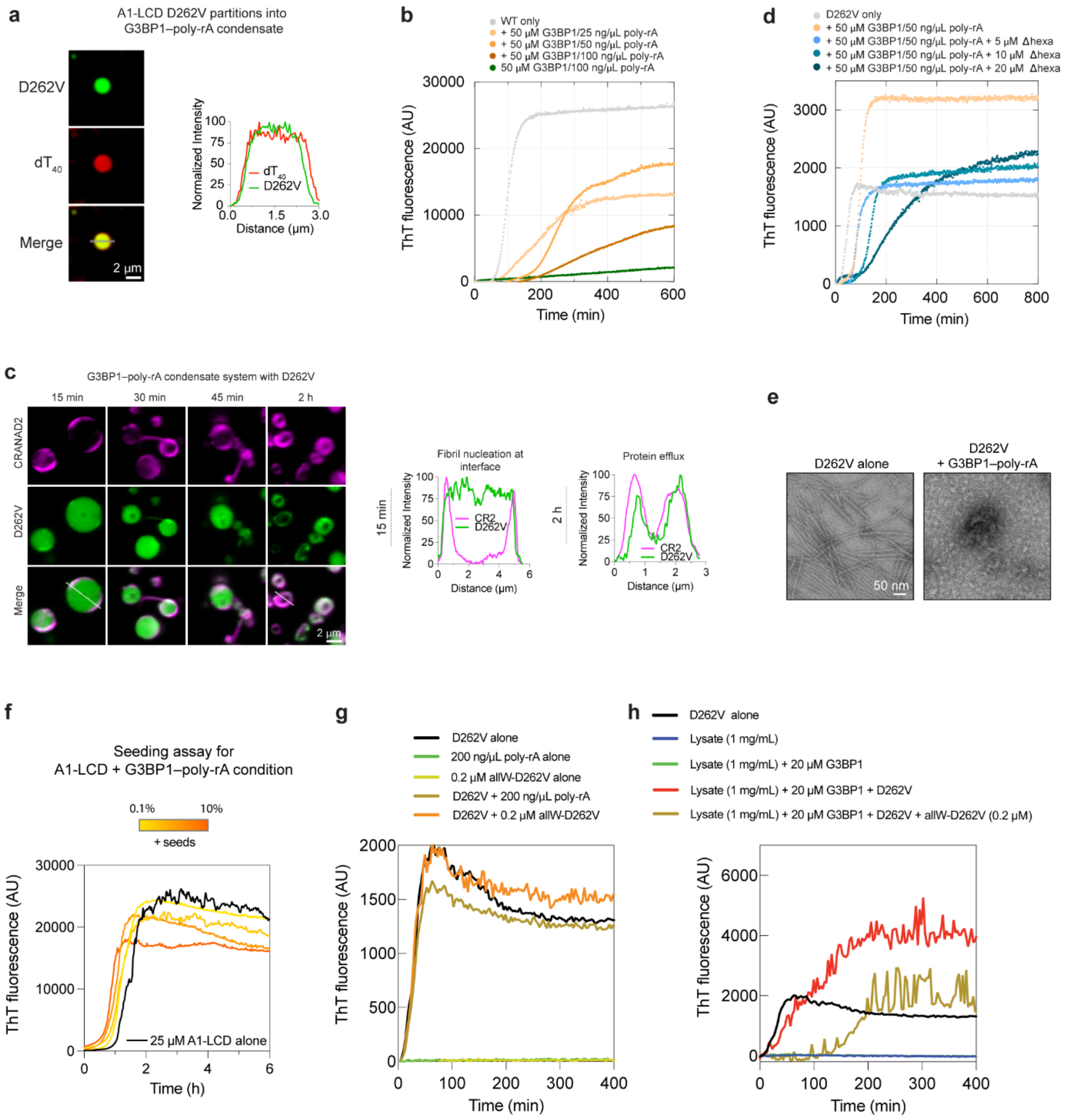


**Figure S12. G3BP1–poly-rA condensates and stress granules reconstituted from mammalian cell lysate suppress A1-LCD fibril formation. (a)** Fluorescence images showing favorable partitioning of D262V (visualized with 250 nM Alexa488-D262V) into G3BP1–poly-rA condensates (visualized with 250 nM of anti-sense Cy5-$dT_{40}$). The corresponding line profiles are shown. **(b)** Kinetics of fibril formation monitored by ThT fluorescence of 25 µM WT A1-LCD either alone or in the presence of G3BP1 and poly-rA at different ratios. Baseline subtraction was applied. G3BP1–poly-rA condensates alone show little ThT signal enhancement under these conditions. **(c)** Fluorescence image time course of G3BP1–poly-rA condensates with D262V (visualized with 250 nM Alexa488-D262V) showing CRANAD2 fluorescence at the interfaces and fibrillar structures appearing in the dilute phase (visualized with CRANAD2), supported by corresponding line profile analyses. Images were adjusted independently for optimal visualization of the coexistence of condensates and fibrils in each condition. **(d)** Kinetics of fibril formation monitored by ThT fluorescence of 25 µM A1-LCD D262V alone, in the presence of G3BP1–poly-rA condensates, and with increasing concentrations of A1-LCD ∆hexa in the presence of G3BP1–poly-rA condensates. Baseline subtraction was applied. **(e)** nsTEM images of the fibrillar assemblies (or a lack thereof) formed by A1-LCD D262V either alone or in the presence of G3BP1–poly-rA condensates. **(f)** ThT kinetics of WT A1-LCD, either alone or in the presence of seeds, from A1-LCD–G3BP1–poly-rA condition, at varying seed concentrations. **(g)** Kinetics of fibril formation monitored by ThT fluorescence in the dilute phase at the indicated concentrations. Addition of allW-D262V does not lengthen the lag phase. **(h)** Kinetics of fibril formation monitored by ThT fluorescence of 20 µM A1-LCD D262V in lysate granules. The lysate concentration is 1 mg/mL and is used either alone or in the presence of G3BP1 (20 µM) and allW-D262V (0.2 µM). In the presence of lysate granules, the addition of allW-D262V lengthens the lag phase. Baseline subtraction was applied to all ThT curves.

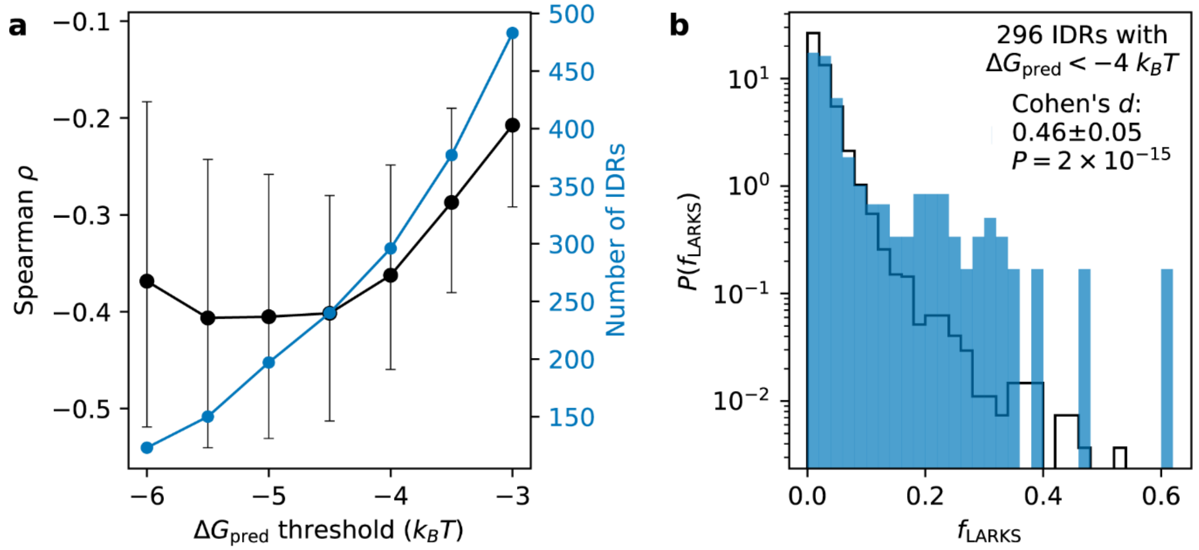


**Figure S13. Association between phase-separation propensity and LARKS fraction in human IDRs.** (**a**) Spearman correlation coefficient between $\Delta G_{\text{pred}}$ and $f_{\text{LARKS}}$ calculated for subsets of human IDRs with predicted free energy of transfer, $\Delta G_{\text{pred}}$, below threshold values ranging from −3 to −6 $k_BT$ (black) and corresponding number of IDRs in the subsets (blue). Error bars represent 95% confidence intervals estimated from $10^4$ bootstraps. (**b**) Normalized distributions of $f_{\text{LARKS}}$ for human IDRs with $\Delta G_{\text{pred}}$< −4 $k_BT$ (296 IDRs; blue bars) and $\Delta G_{\text{pred}} \geq$ −4 $k_BT$ (13,578 IDRs; black line). Cohen's $d$ quantifies the shift between the distributions and the *P* value from a one-sided Brunner–Munzel test indicates the statistical significance of LARKS enrichment in IDRs with $\Delta G_{\text{pred}}$ < −4 $k_BT$.

### Supplementary Notes

#### Note 1: Fibril-forming protein systems

The enhanced FUS-LCD (eFUS-LCD) construct is an engineered variant of the FUS low-complexity domain (LCD) in which the two RAC (reversible amyloid core) motifs, $^{37}$SYSGYS$^{42}$ and $^{54}$SYSSYG$^{59}$, are replaced by the SYNSYS zipper motif (see **Table S1** for complete sequence details). The eFUS-LCD construct is based on the human FUS 'Short' splice isoform (UniProt P35637, VSP_005798; $^{64}$TG$^{65}$→ S relative to the canonical sequence) and comprises residues 1–273 of this isoform. Based on zipper score predictions made by ZipperDB[21], the engineered SYNSYS zipper motif is superior to the native zipper of FUS-LCD driving amyloid formation. The substituted RAC regions contribute to 4 amino acid substitutions to the overall FUS-LCD sequence, thereby largely preserving the intrinsic compositional and physicochemical characteristics of the low-complexity domain.

The SynTag-Tau construct is an engineered variant of full-length microtubule-associated protein Tau (2N4R isoform) in which a 40-amino-acid synthetic prionogenic low-complexity sequence (composed of residue types found in naturally occurring prion-like domains) is appended as an N-terminal tag (see **Table S1** for complete sequence details). In our earlier study, we reported that this synthetic tag substantially lowers the nucleation barrier for Tau condensate-to-fibril conversion[3]. Using ZipperDB analysis, we showed that the putative steric zippers in the tag are outweighed in both score and frequency by those in the native Tau sequence, indicating that the primary drivers of fibril nucleation originate from the Tau core.

These engineered protein variants of FUS and Tau introduce minimal changes to the native protein sequences, which are otherwise recalcitrant to fibril assembly within an experimentally accessible timescale[22,23].

#### Note 2: Sequestration of A1-LCD in the dense phase abrogates the formation of fibrils

2.1 Reconciling light microscopy and nsTEM observations

The observations by light microscopy and nsTEM are seemingly at odds: While ThT kinetics and nsTEM demonstrate the rapid appearance of fibrils in samples without condensates (**Fig. 1c, f**; **Fig. S1a, d**), light microscopy can detect fibrillar structures only in samples that contain condensates (**Fig. S1e, f**). The diameter of single amyloid fibrils is typically too small to be detectable by light microscopy, and we therefore speculate that fibrils are bundled in the presence of condensates, enabling their visualization by light microscopy. These observations demonstrate the challenge of elucidating the underlying processes comprehensively through diffraction-limited light microscopy and point to the requirement of integrating complementary measurements, since no single technique resolves all relevant length scales of fibril assembly.

2.2 Multi-component condensates counteract amyloid fibril assembly and seeding of fibrils

Given that $c_{sf}$ values represent threshold levels for fibril formation, and the dilute phase concentrations could be lower than $c_{sf}$ in samples with high peptide and ssDNA concentrations, A1-LCD fibrils should not grow under these conditions. However, bulk ThT assays and ThT-based fluorescence microscopy suggested fibril formation. To explain this conundrum, we performed nsTEM of sedimentable components in the samples after 6 days of incubation. While WT A1-LCD and the pathogenic variant alone formed fibril bundles, we did not observe typical fibrils when peptide and ssDNA were present in samples (**Fig. S7a**). Instead, we observed smaller, irregular-looking assemblies. We hypothesized that these assemblies were a result of nucleation at interfaces, but that the nuclei had failed to grow into fibrils because of the low dilute–phase concentration. To distinguish this from a model where the structures are off-pathway assemblies, we tested whether they could seed fibril formation. Pure A1-LCD fibrils were sonicated to generate seeds (**Fig. S7b**). We also isolated the sedimentable components of A1-LCD, peptide, and ssDNA–containing samples and treated them in the same way (**Fig. S7b-d**). Each of these preparations was tested for its seeding potential by adding to fresh A1-LCD protein and monitoring fibril growth. WT A1-LCD seeds generated from fibrils shortened the lag phase strongly, and D262V seeds eliminated the lag phase (**Fig. S7c, d**). The sonicated material generated from condensate-containing samples shortened the lag phases, although to a lesser extent, and in a manner that depended on the seed concentration. Overall, these results confirmed that the assemblies formed in condensate-containing samples act as seeds and are thus likely small fibrils or pre-fibrillar in nature. We therefore interpret the ThT fluorescence to stem from nucleation of A1-LCD at the condensate interfaces and aborted fibril growth. Conversely, when we tested whether pre-formed fibril seeds embedded within condensate interiors could promote amyloid growth in the dense phase, we observed no appreciable fibril elongation: the seeds remained largely unchanged in size over several hours, indicating that fibril growth within the dense phase is effectively suppressed or blunted (**Fig. S7e**).

#### Note 3: Partitioning of A1-LCD into different peptide–ssDNA condensates

Partitioning of A1-LCD D262V is expected to be higher in condensate systems in which it can effectively outcompete the peptide for nucleic acid interactions. In our recent work, we showed that $(RGYGG)_5$ forms more stable intermolecular interactions with both ssDNA and ssRNA than $(RGPGG)_5$ or $(KGKGG)_5$ peptides[5,12]. We also demonstrated that, in mono-component systems, stronger inter-chain interactions correlate with higher viscosity[8]. Thus, in $(RGYGG)_5$-$dT_{40}$ condensates, D262V is less effective in competing with the peptide for association with ssDNA as compared to $(RGPGG)_5$ and $(KGKGG)_5$ systems, resulting in lower partitioning. Similarly, weaker peptide–ssDNA interactions in $(RGPGG)_5$ and $(KGKGG)_5$ systems enable greater incorporation of D262V in condensates formed by these peptides.

As a result of the collective favorable three-body interactions, one would expect an increase in condensate viscoelasticity when A1-LCD is added to peptide-ssDNA condensates. This is confirmed by our experiments (**Table S5**), which revealed that the viscosities and the network relaxation time ($\tau_M$: inverse of the cross-over frequency; **Table S4; Fig. 2a**) of the three-component condensates are consistently higher than those of the two-component peptide–ssDNA systems. Importantly, in the presence of A1-LCD, we note that the relative increase in condensate viscosity is greater for weaker nucleic acid–interacting peptide systems, $(RGPGG)_5$ and $(KGKGG)_5$, relative to the stronger nucleic acid-interacting peptides $(RGRGG)_5$ and $(RGYGG)_5$.

#### Note 4: Quantitative modeling of fibril formation at the condensate interface

4.1 Experimental motivation

We asked whether A1-LCD concentrations at interfaces, rather than those in the dilute phase, determine nucleation rates. Using fluorescence microscopy, we determined the partition coefficients of labeled WT A1-LCD or D262V into peptide–ssDNA condensates (**Fig. S8c, d**). The fluorescence intensity of labeled A1-LCD remained homogeneously distributed throughout peptide–ssDNA condensates at early time points. We therefore used the relative partition coefficients of labeled WT and D262V A1-LCD as a proxy for A1-LCD concentrations at condensate interfaces. Strikingly, both A1-LCD variants partitioned more strongly into $(RGPGG)_5$–$dT_{40}$ and $(KGKGG)_5$–$dT_{40}$ condensates, i.e., those with the lowest viscoelasticity, compared to the more viscoelastic $(RGYGG)_5$–$dT_{40}$ condensates. This differential partitioning can be mechanistically explained by competitive interactions between the A1-LCD and the repeat peptides for ssDNA binding (*see* **Supplementary Note 3**). Notably, condensates exhibiting higher WT A1-LCD partitioning also showed shorter fibril nucleation lag times, and vice versa (**Fig. 3h**). This inverse relationship between protein partitioning into condensates and lag time also held true for the D262V variant (**Fig. 3i**). We note that the lag time is an empirical parameter that does not purely represent primary nucleation rate but also includes contributions from secondary nucleation and fibril growth rates. Overall, given that the interiors of condensates suppress fibril formation, our results suggest that the concentration of fibril-forming proteins at interfaces, so-called interface densities, plays an important role in determining the timescale of fibril nucleation.

#### 4.2 Multicompartment kinetic model of fibril formation

To quantitatively test whether the insights into interface densities can help explain the observed nucleation kinetics, we developed a 3-compartment kinetic model. The A1-LCD population partitions across three compartments associated with a spherical condensate of radius $R$: the condensate interior ($c_c$), the surrounding dilute phase ($c_d$), and a thin interfacial layer ($c_i$) at the condensate interface. Two aggregate observables, the fibril number concentration $N$ and the total fibril mass $M$ (in monomer units), complete the description. The kinetic scheme follows the master-equation framework for amyloid assembly developed by Knowles, Dobson and co-workers[24-27], specialized here to explicitly resolve the condensate interface as a distinct compartment in the spirit of recent reports of accelerated nucleation at biomolecular condensate interfaces[28-30]. Interface fluxes are scaled by the area-to-volume ratio $A/V = 3/R$ for a spherical condensate.

##### 4.2.1 Dense to dilute partitioning

Exchange between the condensate interior and the dilute phase is taken to be a first-order process, reflecting fast molecular exchange across a sharp phase boundary[23,31]:

$$J_{c\to d} = k_{\text{out}}\, c_c - k_{\text{in}}\, c_d,$$

so that at equilibrium the partition coefficient is

$$K_{\text{part}} = \frac{c_c^{\text{eq}}}{c_d^{\text{eq}}} = \frac{k_{\text{in}}}{k_{\text{out}}}.$$

##### 4.2.2 Dilute to interface adsorption (Langmuir)

Adsorption to the interface follows a Langmuir form[32] so that detailed balance is satisfied at finite interface capacity. With fractional occupancy $\theta = c_i/c_i^{\max}$,

$$J_{d\to i} = \frac{A}{V}\left[ k_{\text{ads}}\, c_d\,(1-\theta) - k_{\text{des}}\, c_i \right],$$

yielding the standard isotherm at equilibrium,

$$\frac{c_i^{\text{eq}}}{c_i^{\max}} = \frac{K_{int}\, c_d}{1 + K_{int}\, c_d}, \qquad K_{int} = \frac{k_{\text{ads}}}{k_{\text{des}}}.$$

The Langmuir form is preferred over a Hill-type saturation because it gives a simple closed-form expression for the equilibrium interface coverage that depends on just one parameter, the $K_{\text{int}}$. This choice is consistent with prior treatments of amyloid nucleation at two-dimensional surfaces[33,34].

4.2.3 Nucleation

Fibril nuclei form predominantly at the interface from interface-bound monomers, following the primary-then-secondary nucleation scheme[24,25] but with $c_i$ rather than the dilute phase concentration as the kinetic substrate:

$$r_1 = \frac{A}{V} k_{\text{nuc}} c_i^{n^*}, \qquad r_2 = \frac{A}{V} k_2 c_i^{m_2} M$$

Here, $n^*$ is the reaction order of primary nucleation (the kinetic analogue of a critical nucleus size[35,36]), and $m_2$ is the reaction order of secondary nucleation in the existing fibril mass[25,26]. A direct dilute-phase nucleation pathway $r_d = k_{\text{nuc}}^{(d)} c_d^{n^*}$ is available but is ignored for the sake of simplicity ($k_{\text{nuc}}^{(d)} = 0$) for the results described in the main-text panels. This pathway is used only in **Fig. S9** (Sec. 4.4), where we test regimes of dilute versus interface-dominated fibril formation, where this step sets a finite background nucleation rate when the interface is artificially inactivated.

4.2.4 Elongation

Mature fibrils elongate by monomer addition from the dilute pool,

$$r_{\text{elong}} = k_+ c_d N.$$

No explicit lag phase is imposed; the experimentally observed lag phase emerges naturally from the time required to populate the interfacial compartment $c_i$ and reach the nucleation threshold[24,27].

4.2.5 Mass-balance equations

Combining Sec. 4.2.1–4.2.4, the full system reads

$$\begin{aligned}
\dot{c}_c &= -J_{c\to d}, \\
\dot{c}_d &= J_{c\to d} - J_{d\to i} - r_{\text{elong}} - n^* r_d, \\
\dot{c}_i &= J_{d\to i} - n^* (r_1 + r_2), \\
\dot{N} &= r_1 + r_2 + r_d, \\
\dot{M} &= n^* (r_1 + r_2 + r_d) + r_{\text{elong}}.
\end{aligned}$$

Total monomer mass $c_c + c_d + c_i + M$ is conserved by construction and is monitored as a sanity check in every integration step.

4.3 Parameters

The base parameter set used to generate the main-text modeling panels is reported in **Table 1**. The parameter scans (**Table 2**) modify only the indicated parameter while holding all others fixed at the base values. Concentrations and rates are reported in arbitrary units (a.u.); time is converted to seconds in the figures using $1\,\text{a.u.} = 0.1\,\text{s}$, chosen so that simulated lag times fall in the experimentally reported range[30] ($10^3$–$10^4$ s).

**Table 1.** Base parameters.

| Symbol | Value (a.u.) | Description |
|---|---|---|
| $k_{\mathrm{out}}$ | $1 \times 10^{-3}$ | dense to dilute phase efflux |
| $k_{\mathrm{in}}$ | $2 \times 10^{-3}$ | dilute to dense phase uptake |
| $K_{\mathrm{part}} = k_{\mathrm{in}}/k_{\mathrm{out}}$ | 2.0 | partition coefficient $c_c^{\mathrm{eq}}/c_d^{\mathrm{eq}}$ |
| $k_{\mathrm{ads}}$ | $2 \times 10^{-2}$ | dilute to interface adsorption |
| $k_{\mathrm{des}}$ | 1.0 | interface to dilute desorption |
| $K_{int} = k_{\mathrm{ads}}/k_{\mathrm{des}}$ | $2 \times 10^{-2}$ | interface affinity (Langmuir) |
| $c_i^{\mathrm{max}}$ | 10 | maximum interface capacity |
| $k_{\mathrm{nuc}}$ | $1 \times 10^{-5}$ | primary interface nucleation rate constant |
| $n^*$ | 2 | nucleation order in $c_i$ |
| $k_2$ | $1 \times 10^{-7}$ | secondary interface nucleation rate constant |
| $m_2$ | 1 | secondary nucleation order in $M$ |
| $k_+$ | $4.6 \times 10^{-2}$ | elongation rate constant |
| $R$ | 1 | droplet radius (sets $A/V = 3/R$) |
| $c_c(0)$ | 100 | initial monomer in droplet ($c_d = c_i = N = M = 0$) |
| $k_{\mathrm{nuc}}^{(d)}$ | 0 (main); $1 \times 10^{-9}$ (SI) | direct dilute nucleation |

**Table 2.** Parameter sweep ranges.

| Figure | Swept parameter | Range | $N_{\mathrm{pts}}$ |
|---|---|---|---|
| Fig. 4b | $K_{\mathrm{part}}$ (vary $k_{\mathrm{in}}$, fix $k_{\mathrm{out}}$) | $[10^0, 10^{1.2}]$ | 21 |
| Fig. 4c | $K_{\mathrm{int}}$ (vary $k_{\mathrm{ads}}$, fix $k_{\mathrm{des}}$) | $[10^{-3} 10^0]$ | 21 |
| Fig. 4f | $k_{\mathrm{out}}$ (fix $K_{\mathrm{part}} = 2$) | $[10^{-5}, 10^{-1}]$ | 21 |
| Fig. S9a | $c_b(0)$ at $K_{\mathrm{int}} \in \{0.01, 0.1, 0.5, 2\}$ | $[10^1, 10^3]$ | 20 |
| Fig. S9b | $(K_{\mathrm{int}}, c_b(0))$ heatmap | $K_{int} - [10^{-2},\ 10^{0.7}]$, $c_b(0) - [10^1, 10^{2.8}]$ | $16 \times 16$ |
| Fig. S9c | paired traces, interface on/off | $(c_b = 20, K_{int} = 1.0)$, $(c_b = 500, K_{int} = 0.01)$ | n/a |

4.4 Numerical methods and observables

The ODE system is integrated with the Radau implicit Runge–Kutta solver[37] in *scipy.integrate.solve_ivp*, with relative and absolute tolerances $10^{-8}$ and $10^{-10}$, respectively, and a final time $T_1 = 2 \times 10^5$ a.u. ($= 2 \times 10^4$ s). State variables are clamped at zero before each evaluation of the right-hand side to suppress unphysical negative excursions during stiff transients.

Lag-time observable: Each trajectory is summarized by the lag time, which is the time at which fibril mass first reaches 5% of its plateau

$$t_{5\%} = \min\{t : M(t) \geq 0.05\, M(T_1)\},$$

This is a standard kinetic surrogate for the experimentally reported lag phase[25,26] and is used as the dependent variable in the lower row of main-text **Fig. 4** and in the enhancement maps in **Fig. S9**.

Enhancement factor**:** For **Fig. S9**, two trajectories are integrated per parameter point (one with the interface active and one with the interface inactivated by setting $k_{\mathrm{ads}} = 10^{-10}$), and the dimensionless enhancement

$$E = \frac{t_{5\%}^{\mathrm{no\ int}}}{t_{5\%}^{\mathrm{int}}}$$

is reported. $E \gg 1$ indicates that interface-catalyzed nucleation dominates the lag time on the simulated time scale, while $E \approx 1$ identifies the bulk-driven regime in which the interface is dispensable.

4.5 In what kinetic regime does the interface matter?

Here, we set out to determine in which kinetic regime the interface of condensates contributes more significantly to nucleation/elongation than what could take place in the dilute phase. To this end, we make one change to the main model to add a small direct dilute-phase nucleation rate of $k_{\mathrm{nuc}}^{(d)} = 10^{-9}$. This is enabled so that fibril formation remains feasible even when the interface is switched off; all other parameters are as in **Table 1**. **Fig. S9** supports the inference from our experimental data described in **Fig. 3** that the interface is most important when the dilute-phase concentration falls below the spontaneous fibril formation threshold $c_{\mathrm{sf}}$: in our parameterization, this corresponds to the upper-left wedge of panel (b), $c_d^{\mathrm{eq}} \lesssim 30$ a.u. with $K_{\mathrm{int}} \gtrsim 0.1$.

### Note 5: Increase in A1-LCD fibril formation lag times cannot be reproduced by individual condensate-forming components.

We tested whether the presence of individual condensate-forming components, such as the peptide or nucleic acid component alone, can explain the increase in fibril formation lag time that was observed with multi-component condensate systems. To this end, we measured ThT kinetics of A1-LCD D262V in the presence of either peptide–ssDNA condensates [i.e., $(RGRGG)_5$–$dT_{40}$] or the peptide or ssDNA alone [i.e., $(RGRGG)_5$ or $dT_{40}$, respectively]. In the presence of heterotypic condensates, we observe a substantial increase in fibril formation lag times in a volume fraction-dependent manner (**Fig. S2a**). However, in the presence of $(RGRGG)_5$ alone at the same concentrations, the A1-LCD D262V fibril formation kinetics remained unperturbed.

Interestingly, in the presence of $dT_{40}$ alone, we observed a shift in the ThT kinetics towards marginally higher lag times. Upon imaging the various sample conditions using brightfield microscopy, we found that the sample containing A1-LCD D262V and $dT_{40}$ formed condensates, but the sample containing A1-LCD D262V and $(RGRGG)_5$ did not (**Fig. S2b**). Therefore, the mild increment in fibril formation lag time in samples containing A1-LCD D262V and $dT_{40}$ can be attributed to the formation of two-component condensates that sequester A1-LCD in the dense phase and lower the protein concentration in the dilute phase. This is consistent with our proposed model that condensates act as sinks for soluble protein and kinetically suppress their conversion to fibrils [2].

Next, we sought to determine the effect of different condensate-forming peptides and ssDNA of different lengths, which were utilized in our study (**Fig. 2**), on A1-LCD fibril formation lag times. Across different multi-component condensate systems, wherein the peptide sequences were varied, fibril assembly of WT A1-LCD or D262V was markedly suppressed in a volume fraction-dependent manner compared to that of peptide or nucleic acid alone (**Fig. S2c**). With the least viscoelastic condensate system [$(KGKGG)_5$–$dT_{40}$], the protection against fibril assembly offered by the multi-component condensate system relative to nucleic acid ($dT_{40}$) alone appears to be mild (**Fig. S2c**). By contrast, highly viscoelastic condensates [e.g., $(RGRGG)_5$–$dT_{40}$, $(RGYGG)_5$–$dT_{40}$] are superior suppressors of fibril assembly compared to $dT_{40}$ alone (**Fig. S2c**). Similarly, when varying nucleic acid chain length, we observe that the suppression of A1-LCD D262V fibril formation is more pronounced in the presence of multi-component condensate systems than in the presence of A1-LCD D262V and nucleic acid alone (**Fig. S2d**). Hence, the ability of multi-component condensates to suppress amyloid assembly cannot be fully explained by the behavior of individual components, highlighting the emergent properties of multi-component condensate systems.

### Supplementary Tables

**Table S1. List of hnRNPA1 variants and other fibril-forming proteins used in the study.** The proteins have two additional amino acids at the N-terminus (highlighted in italic), which are the remainder after the His-tag is removed using TEV protease. The residue highlighted in red in the A1-LCD D262V sequence corresponds to an ALS-linked disease mutation. In the SynTag-Tau sequence, the synthetic prion-like tag sequence is underlined, whereas the remainder of the sequence comprises the wild-type Tau sequence (2N4R isoform). Residue highlighted in green in the eFUS-LCD sequence corresponds to engineered zipper motifs with a strong propensity to form amyloids, relative to the wild-type zipper motifs.

| S. no. | Name | Sequence |
|---|---|---|
| 1. | WT A1-LCD | *GS*MASASSSQRGRSGSGNFGGGRGGGFGGNDNFGRGGNFSGRGGFGGSRGG<br>GGYGGSGDGYNGFGNDGSNFGGGGSYNDFGNYNNQSSNFGPMKGGNFGGRS<br>SGPYGGGGQYFAKPRNQGGYGGSSSSSSYGSGRRF |
| 2. | A1-LCD D262V | *GS*MASASSSQRGRSGSGNFGGGRGGGFGGNDNFGRGGNFSGRGGFGGSRGG<br>GGYGGSGDGYNGFGNDGSNFGGGGSYN**V**FGNYNNQSSNFGPMKGGNFGGRS<br>SGPYGGGGQYFAKPRNQGGYGGSSSSSSYGSGRRF |
| 3. | A1-LCD Δhexa | *GS*MASASSSQRGRSGSGNFGGGRGGGFGGNDNFGRGGNFSGRGGFGGSRGG<br>GGYGGSGDGYNGFGNDGSNFGGGG------NYNNQSSNFGPMKGG<br>NFGGRSSGPYGGGGQYFAKPRNQGGYGGSSSSSSYGSGRRF |
| 4. | allW D262V A1-LCD | GSMASASSSQRGRSGSGN**W**GGGRGGG**W**GGNDN**W**GRGGN**W**SGRGG**W**GGSRGG<br>GG**W**GGSGDG**W**NG**W**GNDGSN**W**GGGGSYN**V**FGN**W**NNQSSN**W**GPMKGGN**W**GGRS<br>SGP**W**GGGGQ**WW**AKPRNQGG**W**GGSSSSSS**W**GSGRR**W** |
| 5. | hnRNPA1 | MSKSESPKEPEQLRKLFIGGLSFETTDESLRSHFEQWGTLTDCVVMRDPN<br>TKRSRGFGFVTYATVEEVDAAMNARPHKVDGRVVEPKRAVSREDSQRPGA<br>HLTVKKIFVGGIKEDTEEHHLRDYFEQYGKIEVIEIMTDRGSGKKRGFAF<br>VTFDDHDSVDKIVIQKYHTVNGHNCEVRKALSKQEMASASSSQRGRSGSG<br>NFGGGRGGGFGGNDNFGRGGNFSGRGGFGGSRGGGGYGGSGDGYNGFGND<br>GSNFGGGGSYNDFGNYNNQSSNFGPMKGGNFGGRSSGPYGGGGQYFAKPRN<br>QGGYGGSSSSSSYGSGRRF |
| 6. | SynTag-Tau | MKSSHHHHHHGSSNSSSNNNNNNNNNNLGIEENLYFQSNIMAEPRQEFEVM<br>EDHAGTYGLGDRKDQGGYTMHQDQEGDTDAGLKESPLQTPTEDGSEEPGSE<br>TSDAKSTPTAEDVTAPLVDEGAPGKQAAAQPHTEIPEGTTAEEAGIGDTPS<br>LEDEAAGHVTQARMVSKSKDGTGSDDKKAKGADGKTKIATPRGAAPPGQKG<br>QANATRIPAKTPPAPKTPPSSGEPPKSGDRSGYSSPGSPGTPGSRSRTPSL<br>PTPPTREPKKVAVVRTPPKSPSSAKSRLQTAPVPMPDLKNVKSKIGSTENL<br>KHQPGGGKVQIINKKLDLSNVQSKCGSKDNIKHVPGGGSVQIVYKPVDLSK<br>VTSKCGSLGNIHHKPGGGQVEVKSEKLDFKDRVQSKIGSLDNITHVPGGGN<br>KKIETHKLTFRENAKAKTDHGAEIVYKSPVVSGDTSPRHLSNVSSTGSIDM<br>VDSPQLATLADEVSASLAKQGL |
| 7. | eFUS-LCD | *GS*MASNDYTQQATQSYGAYPTQPGQGYSQQSSQPYGQQ**SYNSYS**QSTDTSG<br>YGQS**SYNSYS**QSQNSYGTQSTPQGYGSTGGYGSSQSSQSSYGQQSSYPGYG<br>QQPAPSSTSGSYGSSSQSSSYGQPQSGSYSQQPSYGGQQQSYGQQQSYNPP<br>QGYGQQNQYNSSSGGGGGGGGGNYGQDQSSMSSGGGSGGGYGNQDQSGGG<br>GSGGYGQQDRGGRGRGGSGGGGGGGGGYNRSSGGYEPRGRGGGRGGRGGM<br>GGSDRGGFNKFGGPRDQGSR |

**Table S2. List of polypeptides used in the study.**

| S. no. | Name | Sequence |
|---|---|---|
| 1 | $(RGRGG)_5$ | RGRGGRGRGGRGRGGRGRGGRGRGGC |
| 2 | $(RGPGG)_5$ | RGPGGRGPGGRGPGGRGPGGRGPGGC |
| 3 | $(RGYGG)_5$ | RGYGGRGYGGRGYGGRGYGGRGYGGC |
| 4 | $(KGKGG)_5$ | KGKGGKGKGGKGKGGKGKGGKGKGGC |
| 5 | $(RPRPP)_5$ | RPRPPRPRPPRPRPPRPRPPRPRPPC |

**Table S3. List of nucleic acids used in the study.**

| S. no. | Name | Sequence |
|---|---|---|
| 1 | $dT_{20}$ | dTdTdTdTdTdTdTdTdTdTdTdTdTdTdTdTdTdTdTdT |
| 2 | $dT_{40}$ | dTdTdTdTdTdTdTdTdTdTdTdTdTdTdTdTdTdTdTdTdTdTdTdTdTdTdTdT dTdTdTdTdTdTdTdTdTdTdTdT |
| 3 | $dT_{90}$ | dTdTdTdTdTdTdTdTdTdTdTdTdTdTdTdTdTdTdTdTdTdTdTdTdTdTdTdT dTdTdTdTdTdTdTdTdTdTdTdTdTdTdTdTdTdTdTdTdTdTdTdTdTdTdTdT dTdTdTdTdTdTdTdTdTdTdTdTdTdTdTdTdTdTdTdTdTdTdTdTdTdTdTdT dTdTdTdTdTdTdTdTdTdTdTdTdTdTdTdT |
| 4 | $dT_{200}$ | dTdTdTdTdTdTdTdTdTdTdTdTdTdTdTdTdTdTdTdTdTdTdTdTdTdTdTdT dTdTdTdTdTdTdTdTdTdTdTdTdTdTdTdTdTdTdTdTdTdTdTdTdTdTdTdT dTdTdTdTdTdTdTdTdTdTdTdTdTdTdTdTdTdTdTdTdTdTdTdTdTdTdTdT dTdTdTdTdTdTdTdTdTdTdTdTdTdTdTdTdTdTdTdTdTdTdTdTdTdTdTdT dTdTdTdTdTdTdTdTdTdTdTdTdTdTdTdTdTdTdTdTdTdTdTdTdTdTdTdT dTdTdTdTdTdTdTdTdTdTdTdTdTdTdTdTdTdTdTdTdTdTdTdTdTdTdTdT dTdTdTdTdTdTdTdTdTdTdTdTdTdTdTdTdTdTdTdTdTdTdTdTdTdTdTdT dTdTdTdTdTdTdTdTdTdTdTdTdTdT |
| 5 | $rU_{40}$ | rUrUrUrUrUrUrUrUrUrUrUrUrUrUrUrUrUrUrUrUrUrUrUrUrUrUrU rUrUrUrUrUrUrUrUrUrUrUrU |

**Table S4**. **Crossover frequency and the estimated terminal relaxation time for the peptide–$dT_{40}$–WT A1-LCD condensate systems.** The data is represented as mean ± SEM.

| Peptide–$dT_{40}$–WT A1-LCD | Crossover frequency (Hz) | Terminal relaxation time (ms) |
|---|---|---|
| $(KGKGG)_5$ | 39 ± 3 | 26 ± 3 |
| $(RGPGG)_5$ | 23 ± 2 | 51 ± 5 |
| $(RPRPP)_5$ | 18 ± 2 | 69 ± 7 |
| $(RGRGG)_5$ | 5.4 ± 0.4 | 213 ± 19 |
| $(RGYGG)_5$ | 0.98 ± 0.05 | 1087 ± 77 |

**Table S5. Comparison of viscosities between peptide–$dT_{40}$ and peptide–$dT_{40}$–WT A1-LCD condensate systems.** The viscosity values reported here for the peptide–$dT_{40}$ condensate systems are reproduced from Alshareedah et al. 2024[12]. The data is represented as mean ± SD. 'N.D.' here means "not determined". Note that the viscosity of the two-component $(KGKGG)_5$–$dT_{40}$ and $(RGPGG)_5$–$dT_{40}$ condensates could not be determined because under similar buffer conditions [25 mM MOPS (pH 7.5)] that include 150 mM NaCl, these systems formed very small condensates (diameter ~1 μm) compared to the three-component condensates due to lesser thermodynamic stability.

| Peptide | Viscosity (Pa·s) | |
|---|---|---|
| | Peptide-$dT_{40}$ | Peptide–$dT_{40}$–WT A1-LCD |
| $(KGKGG)_5$ | N.D. | 2.4 ± 0.4 |
| $(RGPGG)_5$ | N.D. | 3.4 ± 0.5 |
| $(RPRPP)_5$ | 0.38 ± 0.01 | 4.8 ± 0.5 |
| $(RGRGG)_5$ | 3.28 ± 0.09 | 6.2 ± 0.2 |
| $(RGYGG)_5$ | 37 ± 1 | 95 ± 5 |

### Supplementary Video Captions

**Video S1:** Maximum intensity projection time-lapse showing ThT-positive fibrils (green) emerge over time from $(RGRGG)_5$–$dT_{40}$ condensates (red; visualized with Cy5-$dT_{40}$); corresponding to Fig. 1i.

**Video S2:** Time-lapse of A1-LCD D262V fibril nucleation at the condensate interface and fibril growth in the dilute phase (red; stained with CRANAD2), and concomitant efflux of A1-LCD D262V (green; visualized with Alexa488-D262V) from $(RGRGG)_5$–$dT_{40}$ condensates; corresponding to Fig. 1j.

**Video S3:** Confocal Z-scan showing fibrils (green; stained with ThT) are present only in the dilute phase and at the interface of the $(RGRGG)_5$–$dT_{40}$ condensates (red; visualized with Cy5-$dT_{40}$); corresponding to Fig. 1k.

**Video S4:** Efflux of A1-LCD D262V (green; visualized with Alexa488-D262V) from the dense phase of $(RGPGG)_5$–$dT_{40}$ condensates; corresponding to Fig. 3c.

**Video S5:** Time-lapse of A1-LCD D262V recruitment and retention in optically-trapped $(RGRGG)_5$–$dT_{40}$ condensates, corresponding to Fig. 5d.

**Video S6:** Time-lapse of A1-LCD D262V recruitment and retention in optically-trapped $(RGRGG)_5$–$dT_{200}$ condensates, corresponding to Fig. 5d.
